# Sample-specific network analysis identifies gene coexpression patterns of immunotherapy response in advanced kidney cancer

**DOI:** 10.1101/2024.11.02.621068

**Authors:** Liangwei Yin, Pietro Traversa, Mohamed Elati, Yamir Moreno, Natalia Marek-Trzonkowska, Christophe Battail

**Affiliations:** Université Grenoble Alpes, IRIG, Laboratoire Biosciences et Bioingénierie pour la Santé, UA 13 INSERM-CEA-UGA, 38000 Grenoble, France; Institute for Biocomputation and Physics of Complex Systems (BIFI), University of Zaragoza, 50018 Zaragoza, Spain; Department of Theoretical Physics, University of Zaragoza, 50018 Zaragoza, Spain; CENTAI Institute, 10138 Turin, Italy; Univ. Lille, CNRS, Inserm, CHU Lille, UMR9020-U1277 – CANTHER – Cancer Heterogeneity Plasticity and Resistance to Therapies, Lille F-59000, France; International Centre for Cancer Vaccine Science, University of Gdansk, Kladki 24, 80-822, Gdansk, Poland; Laboratory of Immunoregulation and Cellular Therapies, Department of Family Medicine, Medical University of Gdańsk, ul. Dębinki 2, 80-811 Gdańsk, Poland

## Abstract

Immunotherapies have recently emerged as a standard of care for advanced cancers, offering remarkable improvements in patient prognosis. However, only a small proportion of patients respond to the treatment, and no definitive molecular hallmark has been identified for clinical use in predicting treatment outcomes. Here, we propose a sample-specific weighted gene network approach to investigate the heterogeneity of patient clinical outcomes by leveraging multiscale network features derived from gene expression data. Our results show that patients exhibiting similar clinical benefits share comparable gene coexpression patterns. Increased gene connectivity and stronger negative gene-gene associations are also pivotal factors in patients with poor prognoses. Moreover, integrating pathway genes with topological measures enables the identification of the perturbed regulation of biological pathways associated with treatment responses. Additionally, sample-level network features enhance the prediction performance of gene expression values-based machine learning models. Collectively, our approach provides valuable guidance on the use of gene network information to stratify cancer patients and to optimize treatment strategies.

## Introduction

Clear cell renal cell carcinoma (ccRCC) is the predominant histological subtype of kidney cancer, with a high mortality rate following metastatic progression [1]. Immune checkpoint inhibitors (ICI) targeting PD-1 and PD-L1, either as monotherapy or in combination with angiogenesis inhibitors, have become the standard of care for metastatic ccRCC in recent years [2, 3]. While these therapies have improved patient survival rates, the objective response rate to nivolumab, an ICI, have been reported to be only 34.1% [4]. Among the response mechanisms to ICI, truncating mutations in *PBRM1* and focal loss of 10q23.31 have been positively associated with patient survival, likely due to the higher expression of angiogenesis genes and the loss of tumor suppressor PTEN, respectively [5–7]. Although immunotherapy aims to enhance immune response against tumors, the proportion of CD8+ T cell infiltration did not correlate with treatment outcome [8]. However, these findings have not been consistently observed in prior studies [9–11], underscoring the complex mechanisms of genomic mutations and T cells in tumor progression and therapy resistance. Therefore, identifying novel predictive markers is crucial for optimizing patient therapies and advancing personalized medicine.

To model the complex system of diseases at the individual level, several methods have been developed to infer sample-specific networks, capturing the unique network structures from multiple samples with different phenotypes. These include single networks based on the partial correlations (P-SSN) [12], sample-specific networks using one sample against a group of given control samples (SSNs) [13], linear interpolation for inferring sample specific network (LIONESS) [14], and sample-specific correlation network (SWEET) [15]. The P-SSN method employs a differential partial correlation analysis between a set of control samples (m) and a specific sample plus the given samples (m + 1). By focusing on direct interactions and excluding indirect interactions, P-SSN’s network distance can distinguish different cancer types or subtypes based on network edges [12]. Similarly, the SSNs method infers sample-specific network using one case sample against a set of control samples as a reference, based on differential Pearson correlation. SSNs validated node and edge markers by literature, function analysis, and pathway enrichment [13], demonstrating the biological significance of network markers. Both P-SSN and SSNs rely on a reference group of healthy samples, which may overlook patient sample heterogeneity across populations. To address this limitation, the LIONESS method used linear interpolation to estimate sample networks by comparing an aggregated network of a group (m) and a perturbed network excluding a case sample (m-1) [14]. While LIONESS may be influenced by population size, the SWEET method introduced genome-wide sample weights in network inference to mitigate this issue. These methods revealed that network degrees of PD1 pathway genes and the *TBC1D* gene were associated with patient’s survival in glioblastoma and lung adenocarcinoma, respectively [15, 16]. However, the inconsistent use of network markers across these methods may restrict the applicability of sample-specific network analysis in cancer research, and sample-specific network features remain to be elucidated as the predictive marker of treatment response in cancer patients.

In our study, we proposed a sample-specific network approach to study the relevance of gene coexpression patterns in the stratification and treatment response of ccRCC patients. We extended the existing SWEET method by incorporating new components, including network similarity and pathway network-based scores, which consider the entire network structure in patients subtyping and integrate network information into signaling pathways. Utilizing transcriptomic profiling data of 309 advanced ccRCC patients collected in clinical trial cohorts, we stratified patients into distinct clusters and identified gene coexpression patterns associated with patient survival through network similarity, network nodes, network edges, and pathway network-based scores. These network features enhanced the prediction performance of gene expression value-based machine learning models. Additionally, we validated the relevance of pathway network-based scores on an independent cohort of advanced ccRCC patients treated with avelumab and axitinib. In summary, our method not only offers a comprehensive strategy to explore gene coexpression patterns from general network structure to specific network markers for patient stratification and treatment prediction, but also complements single sample pathway enrichment analysis in current cancer research.

## Material and methods

### Data source

Gene expression, genomic mutation, and clinical data of 309 ccRCC patients were obtained from the publication by Braun et al. (CheckMate 009, 010, 025) [8]. Sequencing data were generated prior to treatment, and survival data was collected after treatment. The expression data of these samples were split into four groups based on the treatment and cancer site in patients (Table 1). Ensembl gene ID was converted to gene symbol using a human gene annotation file (release v43) from the GENCODE database version 43 [17] and only immune genes, mitochondrial genes, long non-coding RNA genes, and protein-coding genes were kept for gene network inference. To validate the constructed networks, RNA-seq data from 85 normal kidney cortex tissue samples were obtained from the GTEx portal, and cancer-related genes were collected from the Cancer Gene Census database [18]. Additionally, data from an independent ccRCC cohort was downloaded from the publication by Motzer et al. [19], including expression profiles of 354 patients treated with avelumab (anti PD-L1) and axitinib (anti-angiogenic). Gene expression values were normalized using the log_2_ transformation of transcripts per

**Table 1.**
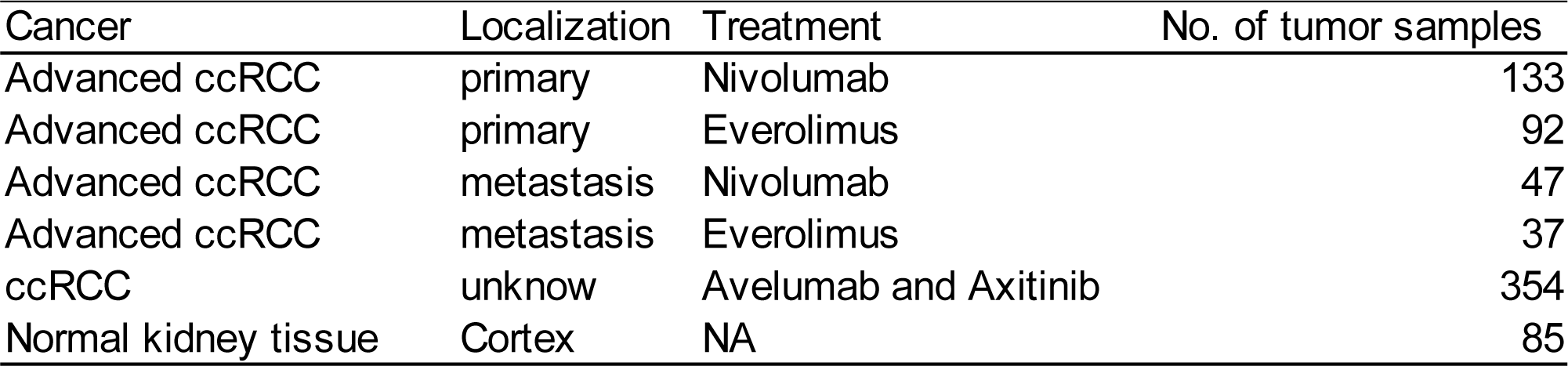
We collected RNA-seq data from several studies, Including the Braun 2020 paper, the Motzer 2020 paper and the GTEx portal.

### Gene network inference

In this study, we used the recently developed SWEET method to construct single-sample weighted gene coexpression networks (ssGCNs) [15]. An aggregated network (*N_ij_^G^*) was first constructed using gene expression of all samples within a specific category. Subsequently, a perturbed network (*N_ij_^G-S^*) was generated by removing one specific sample from the aggregated network (Details in Supplementary Note 1 and Figure 1A). Specifically, a sample-specific network 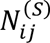 was defined as

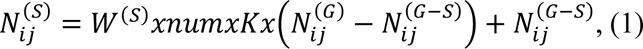

**Figure 1.**
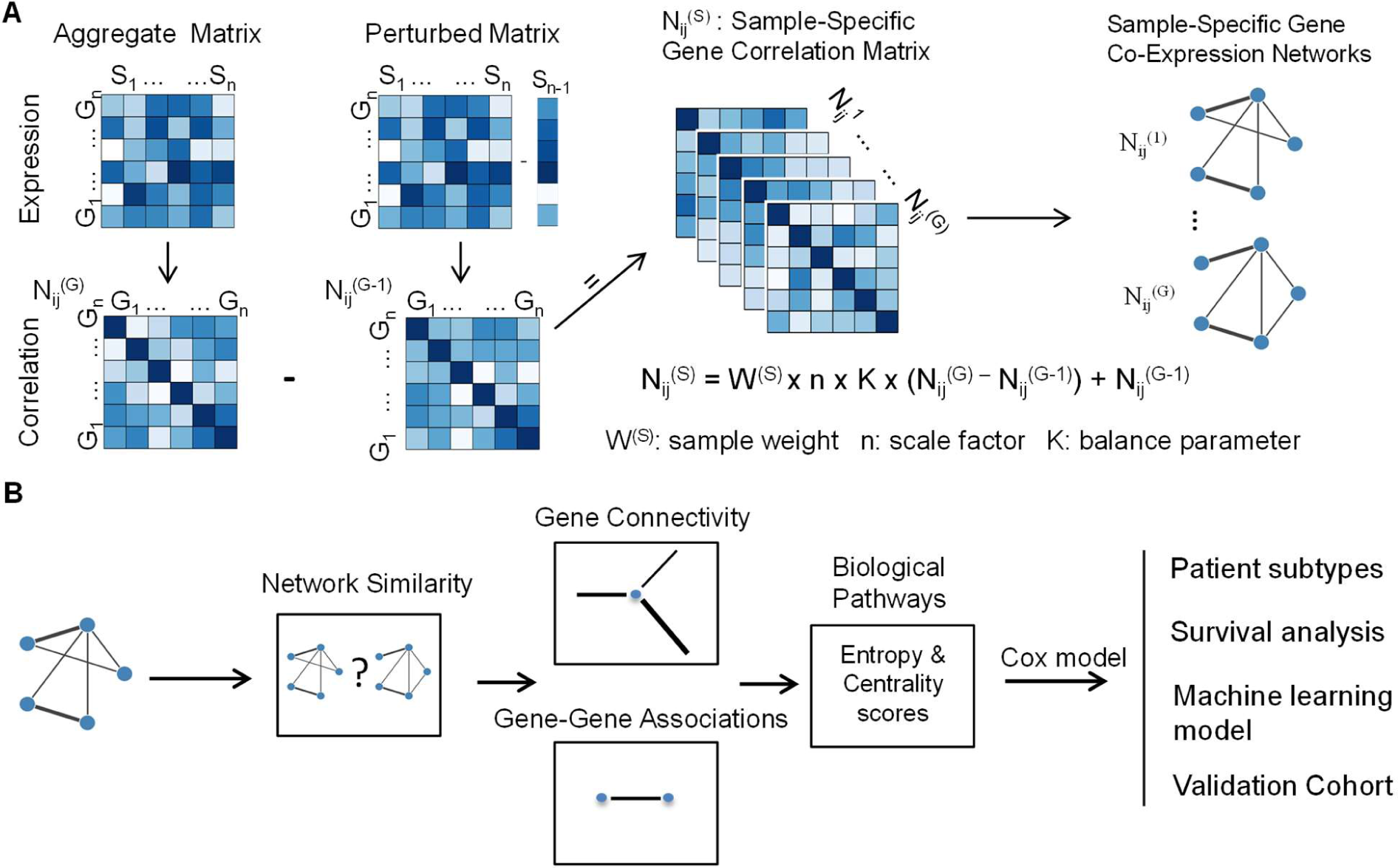
A computational framework for the inference of sample-specific gene coexpression networks and calculation of network features to stratify patients based on their response to anti-tumor therapies. (A). Description of sample-specific gene network construction with the SWEET method. Each sample network was constructed with the difference between a cohort-level aggregate correlation matrix and a sample-specific perturbed correlation matrix. Sample weight (W(^S^)), scale factor (n, i.e. number of samples) and balance parameter (K) were used to adjust for differences in proportions of sample subgroups within a cohort. (B). Description of our pipeline for patient subtyping and survival analysis using network features calculated from sample-specific gene networks. Network similarity was measured by adjusted network distance. Gene connectivity and gene-gene edges were calculated using both the number and the strength of associations between genes. Biological pathway entropy and centrality scores embedded the complexity and the topology of gene network within each pathway.

where *num* was the total number of samples except the target sample in a group, *W*^(S)^ was the sample weight, and *K* was a balance factor ranging from 0 to 1. The best performance was achieved with *K*=10% as shown in Supplementary Figure S1 and by the SWEET paper. The parameter *K* was a scale factor used to enlarge the differential correlation between the aggregated matrix and the perturbed matrix. The sample weight *W*^(S)^ was added to the equation of sample-specific networks to neutralize the network edge number bias [15]. *W*^(S)^ was calculated as

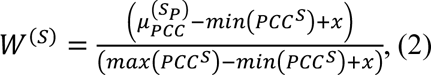

where 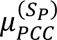 was the mean of Pearson Correlation Coefficient (PCC) between one specific sample S and the other samples, *PCC^S^* was the set of PCCs between two patients and *x* was a constant term added to avoid division by zero that we set to 0.01. To reduce the noise within the networks, the significance level of confidence scores for edges was assessed using z-score normalization, with a z-score threshold of 2.58, corresponding to a two-sided p-value of 0.01 (Details in Supplementary Note 1).

### Network distance

Network distance (*Nd*) was introduced by Huang et al. as a measure of network similarity, primarily used to identify cancer types [12]. *Nd* was defined as the ratio of the number of overlapped edges to the number of union edges between two partial correlation-based sample networks,

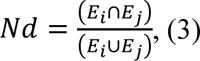

where *E*_i_ and *E*_j_ represented the sets of edges from sample-specific networks. These edges correspond to direct interactions between genes.

However, due to the difference between Pearson correlation and partial correlation, network distance may not be suitable for subtyping patients when using Pearson correlation-based gene networks. Befitting the divergence of treatment response, we proposed a cohort-level adjusted network distance, which calculated the similarity between an individual sample network and a cohort-level network. Cohort-adjusted distances were calculated using clinical benefits (CB) (Equation 4) or non-clinical benefits (NCB) cohort-level networks (Equation 5). Also, the difference between CB distance and NCB distance was used for further analysis (Equation 6).

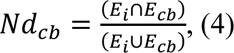

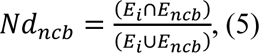

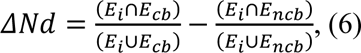

### Network features

We explored two key network features: gene connectivity (weighted node degree) and gene-gene association (edges) (Figure 1B). Gene connectivity, which quantifies the total strength of all the associations of one gene with other genes, was calculated using the Python package igraph (version 0.10.4) [20]. To accurately identify whether the source of gene connectivity was from positive or negative regulations, networks were divided into two: one containing positive correlation edges and the other containing negative correlation edges. To reduce the computation time, the analysis was limited to the top 5,000 most varied genes based on their connectivity (Details in Supplementary Note 2).

To assess the impact of edges, we focused on common edges across all samples within a cohort and selected the top 10,000 most variable edges based on their weights. Furthermore, genes and edges were filtered using a univariate Cox regression model, retaining those with a p-value less than 0.01. To align with the definition of clinical benefits, network features were preserved if they were significantly associated with both overall survival (OS) and progression free survival (PFS).

### Pathway entropy and topology scores

Sample-specific pathway networks were extracted from sample networks using gene sets of pathways from the KEGG database [21]. To enable the calculation of pathway scores, both positive and negative edge values (converted to their absolute values) were considered. Given that the noise elimination within ssGCNs led to a varying number of edges, pathway entropy was calculated based on the distribution of edges [22] as follows:

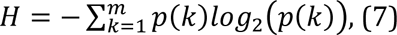

where *p(k)* is the probability of an edge inside a selected pathway network, and *m* is the total number of edges inside that pathway network. The probability *p(k)* of each edge was determined by dividing the weight of the edge by the sum of all the edge weights inside the pathway network.

Additional pathway topological scores were calculated based on the average of gene eigenvector centrality, gene closeness centrality, and edge betweenness centrality (Details provided in Supplementary Note 3) [23]. Alongside our entropy and topology scores, sample-level pathway scores based on gene expression were calculated using the gene set variation analysis (GSVA 1.46.0) method implemented in R (version: 4.2.3) [24].

### Clustering of samples and survival analysis

Unsupervised hierarchical clustering was conducted using the Ward method and cosine metrics. To facilitate comparisons of treatment responses, we divided the samples into two clusters. To assess survival probability between clusters, Kaplan-Meier curves were plotted, and log-rank tests were performed to determine whether the survival distribution of the two clusters were significantly different with a p value of 0.05. These analyses were conducted using the Python packages lifelines 0.27.7 and scikit-survival 0.21.0 [25, 26].

### Gene sets and Transcription Factor (TF) enrichment analysis

The Molecular Signatures Database (MSigDB) hallmark, the KEGG canonical pathways, and the cell marker (augmented 2021) were obtained from the Human MSigDB website and Enrichr libraries [21, 27, 28]. For over-representation analysis, significant pathways were identified using the Python package gseapy 1.0.5 [29] and an adjusted p-value threshold of 0.05 was selected as a threshold. For the enrichment of transcription factors (TF), an online query was conducted on the CHEA3 online tool[30].

### Measuring the performance of machine learning (ML) predictions

To evaluate the prediction performance based on gene expression and network feature values, we used the Logistic Regression (LR) model, and also tested Random Forest (RF) and Support Vector Classifier (SVC) models. Genes (n=64) were selected based on their expression values against survival data using the univariate Cox regression model. The dataset was initially split into a training set and a test set with 30% of the data allocated to the test set. The training set was used to train ML models and the test data was used to evaluate the performance of ML models. The area under the receiver operating characteristic curve (AUC) was used as the main performance metric for machine learning models. We also validated the performance of ML models using Leave One Out Cross Validation (LOOCV) on the whole dataset.

For predictions based on network features and the combination of gene expression and network features, we utilized these features values or their combination to train and test the ML model. For pathway network scores, the number of selected pathways (using the Cox model) was limited and the performance of pathway entropy-based score was compared with the GSVA pathway score. All analyses were implemented in Scikit-learn in Python [31].

### Validation of pathway scores

To assess whether our pathway scores were effective in other cohorts, we selected the data from 354 ccRCC patients treated with the combination of avelumab (anti-PD-L1) and axitinib (anti-angiogenic) [19]. For each sample in this cohort, sample-specific gene networks were constructed and pathway networks were then derived as described above.

## Results

### Inference of sample-specific gene coexpression networks

Sample-specific weighted gene coexpression networks (ssGCNs) were constructed with 20,545 genes using the SWEET method from a cohort of 309 advanced ccRCCs patients included in CheckMate 009, CheckMate 010 and CheckMate 025 clinical trials (Figure 1A) [8, 15]. To accurately study the differences in patient treatment response, the cohort was divided into four subcohorts: 133 and 92 samples from primary tumor sites of patients treated with nivolumab (pN, anti PD1) or everolimus (pE, mTOR inhibitor) respectively, and 47 and 37 samples from tumor metastases of patients treated with nivolumab (mN) or everolimus (mE) respectively. With an optimal balance parameter at 10% and a two-sided z-score threshold of 2.58, the ssGCNs achieved an average network density of 1.6%, comprising 20,357 nodes and 3,320,160 edges, with an average determination coefficients R^2^ of 0.696 for scale-free topology (Supplementary Figure S1). The closer the R^2^ value is to 1, the better the ssGCNs follow the power*-*law node*-*degree distribution expected for biological networks. Notably, the R^2^ coefficients for sample-specific networks from primary tumor sites were higher compared to those from tumor metastases (0.774 vs 0.448, Wilcoxon rank-sum test, p-value < 0.01) (Supplementary Figure S2A). To assess whether subcohort size influenced the R^2^ coefficient of the gene-degree distribution, a simulation was performed on the pN subcohort by randomly selecting 40 to 120 samples. The results showed a decrease in cohort size was associated with a reduction in the mean of R^2^ coefficients, likely reflecting a loss of robustness of the gene network (Supplementary Figure S2B).

Cancers often displayed varied gene network complexities, with acquired network nodes enriched in metabolic and immune related processes such as regulation of immune response, T cell receptor signaling pathway and podosome assembly [32]. To explore whether our ssGCNs exhibited tumor-specific features, we compared their network density and enrichment of cancer-related genes to a cohort-level network from expression data of normal renal cortex tissues. We found that our ccRCC ssGCNs displayed a higher network density (1.6%) compared to the normal network (0.46%). In addition, 98.3% (304/309) of our ssGCNs had a higher enrichment of cancer-related genes among the top 1000 genes of the highest degree than the normal network (Supplementary Figure S3). These characteristics validate the relevance of our ccRCC sample-specific networks for further exploration using advanced network features.

To identify novel network-based markers for predicting immunotherapy response, we focused below on the pN and mN subcohorts of patients treated with nivolumab. The pE and mE subcohorts were used primarily to validate the biological relevance of network features.

### Adjusted network distance in Pearson Correlation-Based ssGCNs

Network distance, a measure of similarity, has been used to estimate gene regulation similarity between samples and accurately identify tumor subtypes [12]. To assess whether network distance could reflect clinical status similarity, we calculated pairwise network distances between ssGCNs of patients in the pN subcohort (Equation 3). Unsupervised clustering based on those network distance did not show significant association with survival data (Supplementary Figure S4; log-rank tests: p-value > 0.2 for both OS and PFS). The limited sensitivity of network distance may be explained by the lesser divergence between gene networks of patients responding and non-responding to immunotherapies than between the gene networks separating tumor subtypes.

To address this, we developed an adjusted version of network distance incorporating clinical information. This adjusted network distance was calculated using edges between each sample and an aggregated network constructed based on clinical outcomes. For the pN and mN subcohorts, clinical benefit (CB) aggregated networks were built from 44 and 13 samples, resulting in 2,450,498 and 720,447 edges, respectively. Non-clinical benefit (NCB) aggregated networks were built from 46 and 20 samples, resulting in 2,853,404 and 1,758,449 edges, respectively. Three versions of adjusted network distances were computed relative to the CB network, the NCB network, and the difference between the two.

Univariate Cox regression analysis revealed that adjusted network distances correlated with survival data (Figure 2A). For the pN subcohort, network distances adjusted to the CB aggregated network were favorable for OS and PFS, though not significantly. In contrast, distance adjusted with NCB aggregated networks were significantly unfavorable for OS and PFS (the Cox model: p-value < 0.05). Adjusted distance using both CB and NCB aggregated networks showed more pronounced favorable associations with OS and PFS. Comparison between CB and NCB patients revealed that CB Patients had significantly higher network distances when adjusted with CB or both CB and NCB networks, and lower distances when adjusted with NCB networks alone (Figure 2B; Wilcoxon sum-rank test; p-value < 0.01). Survival analysis based on the adjusted network distances showed that patients with higher distances were significantly associated with greater OS and PFS (Figure 2C; log rank tests, p-value < 0.01). Higher network distances also correlated with greater OS for the mN subcohort, and with both OS and PFS for the pE subcohort (Supplemental figure S5ABC). In conclusion, our results demonstrate that in the context of ssGCNs generated from Pearson correlations, network distance adjusted with prior clinical knowledge effectively groups patients based on their clinical response to nivolumab.

**Figure 2.**
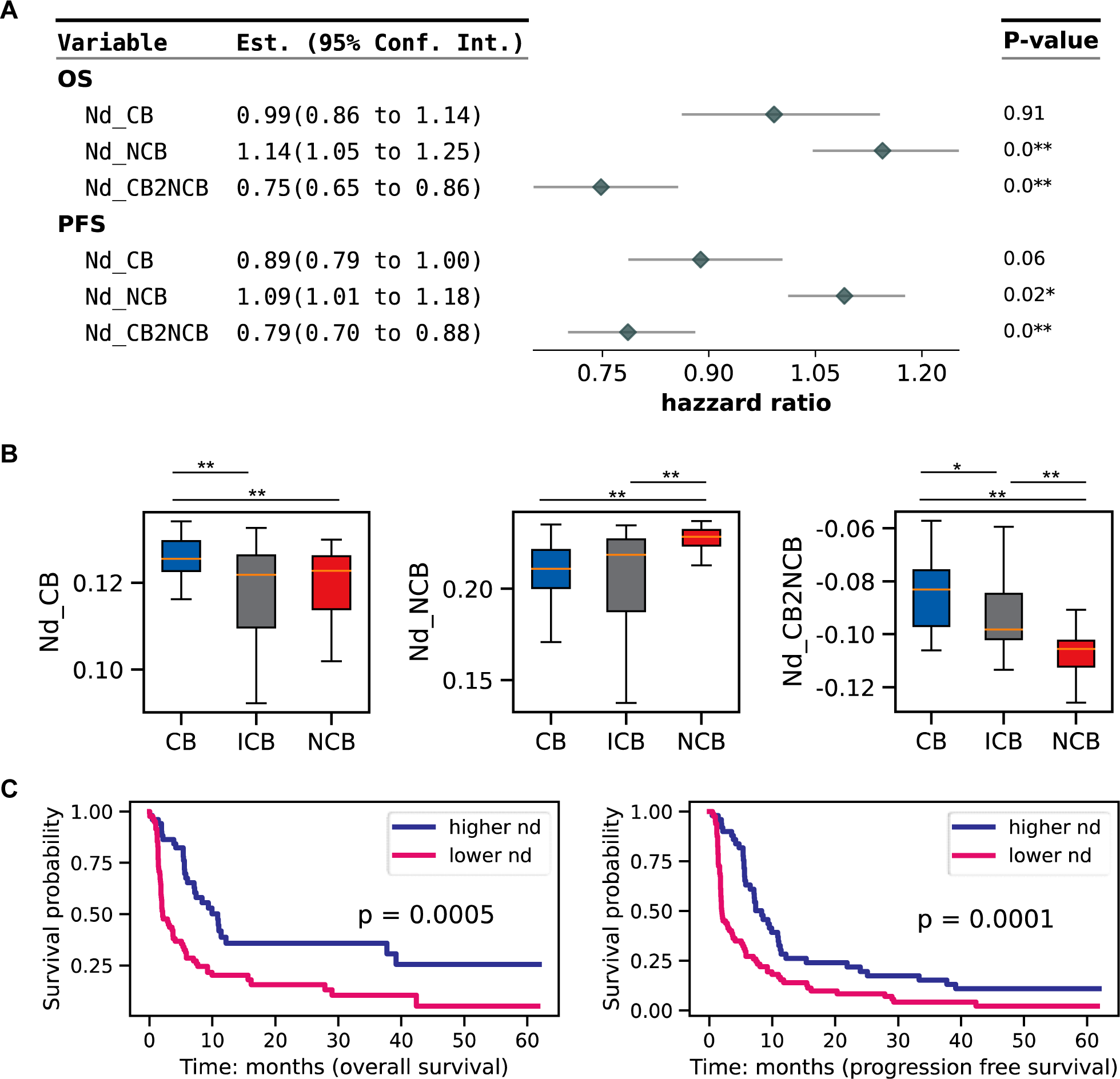
Survival analysis with the adjusted network distance calculated on the subcohort of tumor primary sites from patients followed after immunotherapy by Nivolumab (pN). (A). A forest plot depicting the univariate Cox regression results using adjusted network distances (Nd). Sample-specific network distances were adjusted with the cohort-level network associated with Clinical Benefit (CB), Non-Clinical Benefit (NCB) or the difference between them. (B). Comparison of the distributions of adjusted network distances between clinical benefits categories using Wilcoxon sum rank tests. (C). Survival analysis using network distances adjusted with the differences between CB and NCB cohort-level networks. Samples were divided into two groups based on the median value of the adjusted network distance (nd) (higher nd and lower nd groups). P values were calculated using the log-rank test. (**: p value < 0.01; *: p value < 0.05)

### Gene connectivity is associated with treatment response

Gene degree, or connectivity, has been linked to cancer subtypes and survival outcomes [13, 15]. Gene connectivity refers to the sum of total connections it has with other genes in a weighted network. We hypothesized that gene connectivity could distinguish patient treatment response. First, gene connectivity matrices were separately generated from positive and negative Pearson correlations of ssGCNs to avoid mixing positive and negative associations. From positive correlations matrix, we identified 21 genes whose connectivity significantly associated with OS and PFS in the pN subcohort (Figure 3A; the Cox model: p-value < 0.01; Table 2). Notably, only 8 were also linked to survival based on their expression values. Unsupervised clustering was then performed to split samples into two groups based on those 21 genes’ connectivity. We found that cluster 2 had higher gene connectivity on average (Figure 3B) and significantly lower survival probability for both PFS and OS compared to cluster 1 (Figure 3C; log rank tests: p-value < 0.01). Next, we investigated whether genomic mutation and clinical features differed between the two clusters. Cluster 2 showed that significantly higher frequencies of chromosomal losses at 11q12.3 and 11q23.1, as well as greater intratumor heterogeneity (ITH) (Figure 3D; Supplementary Figure S6; Fisher’s exact test: p-value < 0.05). Relevantly, 11q23 deletion has been linked with poor prognosis in several cancers [33, 34] and ITH has been associated with tumor progression and response to immunotherapy [35]. Gene ontology analysis revealed that the 21 highly connected genes were enriched in cancer and immune-related pathways such as MYC targets v1, oxidative phosphorylation, ribosome pathways, and natural killer T cell gene set (a cell population predictive of the response to anti-tumor treatments) (Figure 3E) [36, 37]. Also, genes like *NLRC5* and *PLCB3*, known for their involvement in ccRCC progression and tumor immunity, were also among these highly connected genes (Details in Supplementary Note 4).

**Figure 3.**
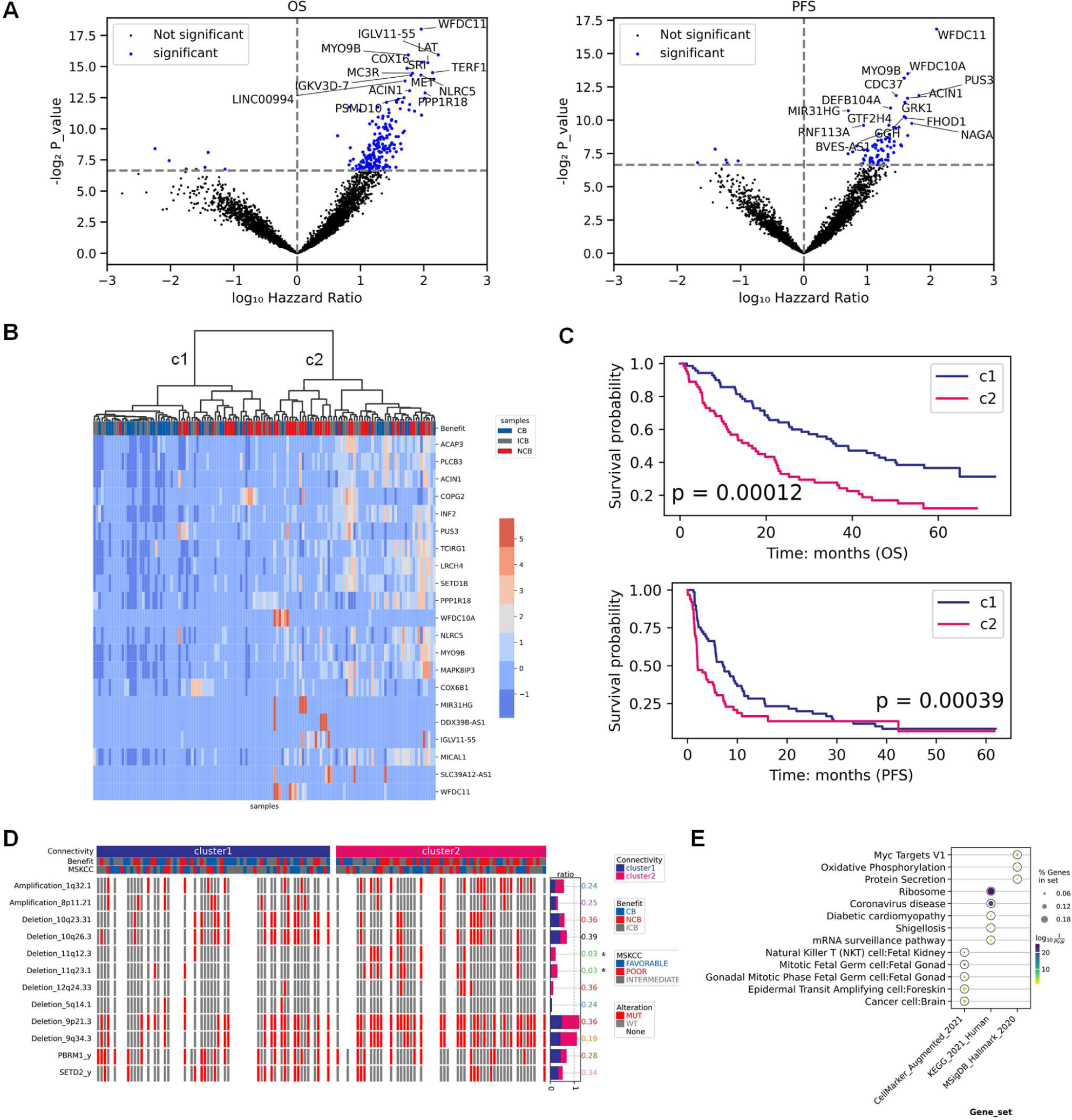
Survival and exploratory analysis using gene connectivity (positive correlation) calculated on the subcohort of tumor primary sites from patients followed after immunotherapy by Nivolumab (pN). (A). Volcano plots of gene connectivity association with patient overall survival (OS) and progression free survival (PFS) (Cox’s proportional hazards model, P value < 0.01). (B). Unsupervised hierarchical clustering of samples into two groups (c1 and c2) based on the connectivity of 21 genes significantly associated with both OS and PFS. Out of these 21 genes, expression values of 8 genes (’SLC39A12-AS1’, ‘WFDC10A’, ‘MYO9B’, ‘TCIRG1’, ‘WFDC11’, ‘MIR31HG’, ‘DDX39B-AS1’, ‘IGLV11-55’) were also associated with survival data. (C). Survival analysis between the clusters c1 (blue) and c2 (pink) of samples. P values were calculated using log rank test. (D). Distribution of chromosomal and gene mutations between the two clusters c1 and c2. Fisher test was conducted and p values less than 0.05 were considered as significant. (E). Gene ontology over-representation analysis. Genes were selected as the union of significant genes whose connectivity was associated with OS or PFS. Gene sets from KEGG, MSigDB hallmark and Cellmarker databases were used. For each pathway, the color of each circle represents the adjusted p-value and the size of circles indicates the percentage of selected genes in the gene sets.

A similar analysis of gene connectivity from negative correlations identified 48 significant genes, with greater connectivity associated with worse survival in the cluster 2 (Supplementary Figure S7 and Details in Supplementary Note 4). To study whether these findings were relevant for tumor metastases, we analyzed the mN subcohort and obtained 9 and 17 genes with higher connectivity (based on positive and negative associations, respectively) that were significantly associated with lower OS and PFS (Table 2; Supplementary Figure S8AB and Details in Supplementary Note 4). Interestingly, there was no overlap between these genes and those identified from the pN subcohort.

The same approach was applied to the pE and mE subcohorts. Consistently, patients with higher gene connectivity values displayed significantly lower survival probability (Supplementary Figure S9 and S10). Overall, our results indicate that the increase of gene connectivity—whether from positive or negative gene-gene correlations—is associated with poorer prognosis and shorter treatment recurrence. Furthermore, the genes associated with patient survival or cancer recurrence differed depending on the tumor primary or metastatic location.

### Highly negative gene-gene associations in patients without clinical benefits

Investigating gene pairwise associations in sample-specific networks has become a popular method for identifying cancer subtypes [12, 38]. To study perturbations in gene-gene associations, we first focused on edges shared by all samples. For ssGCNs of the pN subcohort, 238,804 edges were selected and filtered based on their variance (top 10,000 varied edges). From these, we identified 214 and 224 significant edges associated with OS and PFS values (the Cox model: p-value < 0.01), respectively, with 51 edges intersecting between two (Figure 4A; Table 3). A small proportion of genes from these edges overlapped with those previously identified by connectivity (Supplemental Figure S11). Unsupervised clustering of these edges revealed that cluster 2 had stronger negative gene associations (Figure 4B) and was significantly associated with worse OS and PFS (Figure 4C; log rank test: p-value < 0.01). However, these clusters were not associated with significant differences in genomic mutation or other clinical features (Supplementary Figure S12).

**Figure 4.**
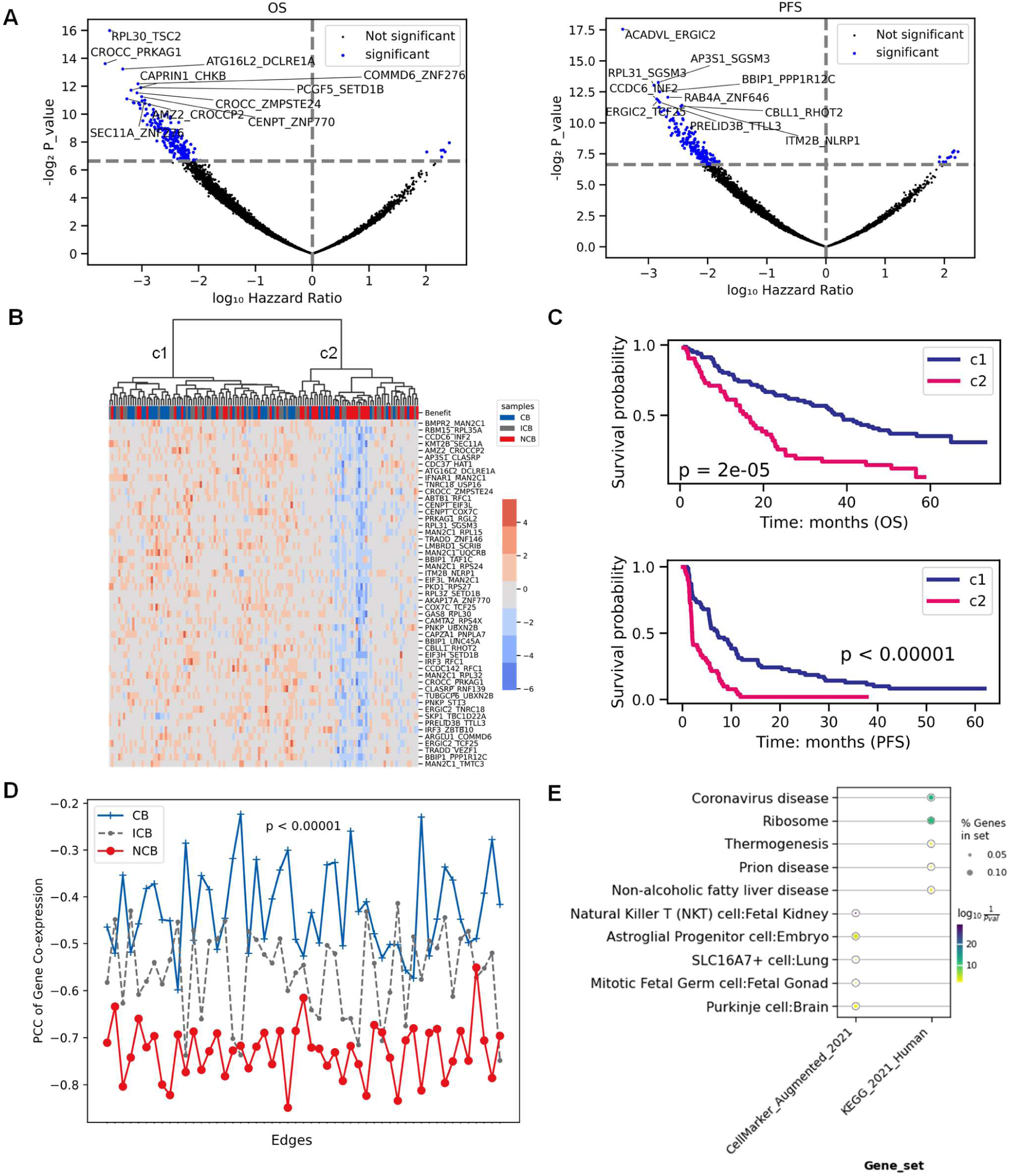
Survival analysis of gene-gene associations using the subcohort of tumor primary sites from patients followed after immunotherapy by Nivolumab (pN). (A). Volcano plots of gene-gene weight association with overall survival (OS) and progression free survival (PFS) (Cox’s proportional hazards model, P value < 0.01). (B). Unsupervised hierarchical clustering of samples into two groups (c1 and c2) using 51 edges weights significantly associated with both OS and PFS. (C). Survival analysis between the clusters c1 (blue) and c2 (pink) of samples. P values were calculated using the log rank test. (D). Distributions of Pearson correlation coefficients (PCC) for pN samples from the Clinical Benefit (CB, blue), Intermediate Clinical Benefit (ICB, grey) and Non-Clinical Benefit (NCB, red) groups. Wilcoxon rank sum test was conducted between CB and NCB patients. (E). Gene ontology over-representation analysis. Genes were selected as the union of significant genes whose edge weight was associated with OS and PFS. Gene sets from KEGG, MSigDB hallmark and Cellmarker databases were used. For each pathway, the color of each circle represents the adjusted p-value and the size of circles indicates the percentage of selected genes in the gene sets.

We further analyzed edge weights across patient categories, finding that non-clinical benefit (NCB) patients exhibited stronger negative correlations compared to clinical benefit (CB) patients (Figure 4D; Wilcoxon rank sum test: p-value < 0.01). It raised the question of whether transcription factors (TF) regulated genes from these edges. Enrichment analysis identified 3 significant TFs (*MZF1*, *ZNF692*, *RBCK1*) were associated with cancer progression and the prognosis in ccRCC (Table 4) [39–41]. Genes from these edges associated with both OS and PFS were over-represented in ribosome pathways and natural killer T cell gene sets (Figure 4E), with some known cancer-related genes (*PRELID3B*–*TTLL3, BMPR2*– *MAN2C1, etc*) among these edges [42–44].

For the mN subcohort, we identified 6 edges significantly associated with both OS and PFS values (Table 3; Details in Supplementary Note 5). Clustering of these edges revealed that cluster 2 was associated with significantly shorter OS and PFS (Supplemental Figure S13AB). Similarly, 5 of these edges were highly negatively co-expressed in NCB patients compared with CB patients (Supplemental Figure S13C).

In the pE and mE subcohorts, our approach identified 40 and 10 significant edges, respectively, which divided samples into two clusters with significantly different survival outcomes (Supplementary Figures S14AB). Visualization of the selected edges across all four subcohorts is shown (Supplementary Figure S15). Overall, highly negative gene-gene associations were linked with poor prognoses. Notably, genes from significant edges weakly overlapped (46.9% for pN) with significant genes based on their connectivity, indicating that edges and gene connectivity offer distinct insights into patient survival predictions. Furthermore, the differences in significant edges between primary and metastatic tumor sites underscore the importance of considering tumor site-specific analyses in cancer research.

### Pathway entropy and centrality scores

Given that both gene connectivity and gene-gene associations revealed differences in patient response to treatments, we next asked whether the complexities and topology features of biological pathways were also associated with patient survival. Previous studies found that survival was associated with the complexity of signaling pathways in pan-cancer molecular data [45]. To explore this, we developed a tool to calculate topology-based pathway scores at single-sample level, which first extracted pathway networks from our ssGCNs using KEGG gene sets. We then computed pathway entropy and centrality scores. While entropy measures the randomness or complexity of a network, eigenvector centrality reflects the transitive influence of genes, closeness centrality indicates the average shortest distance from one gene to the others, and edge betweenness centrality measures the influence of edges inside a network.

To evaluate whether these pathway scores captured specific biological significance, we selected pathways significantly associated with OS or PFS values using the Cox model with a p-value threshold of 0.05. Notably, these pathways rarely overlapped with those identified using the GSVA method based on gene expression values (Supplementary Figure S16). We then use the significant pathway scores to stratify patients into two clusters to assess their predictive power (Supplementary Figure S17). In the pN subcohort, all categories of pathway scores successfully clustered patients according to OS (log rank test: p-value < 0.05) (Figure 5, left). The most robust classifications were achieved with eigenvector and edge betweenness centrality scores. For treatment response, all pathway score categories except edge betweenness centrality were significantly associated with PFS (Figure 5, right; log rank test: p-value < 0.05), with eigenvector centrality showing the highest significance. Consistent with the patterns of gene connectivity and edges, patients exhibiting higher survival displayed lower entropy scores (Supplementary Figure S17B).

**Figure 5.**
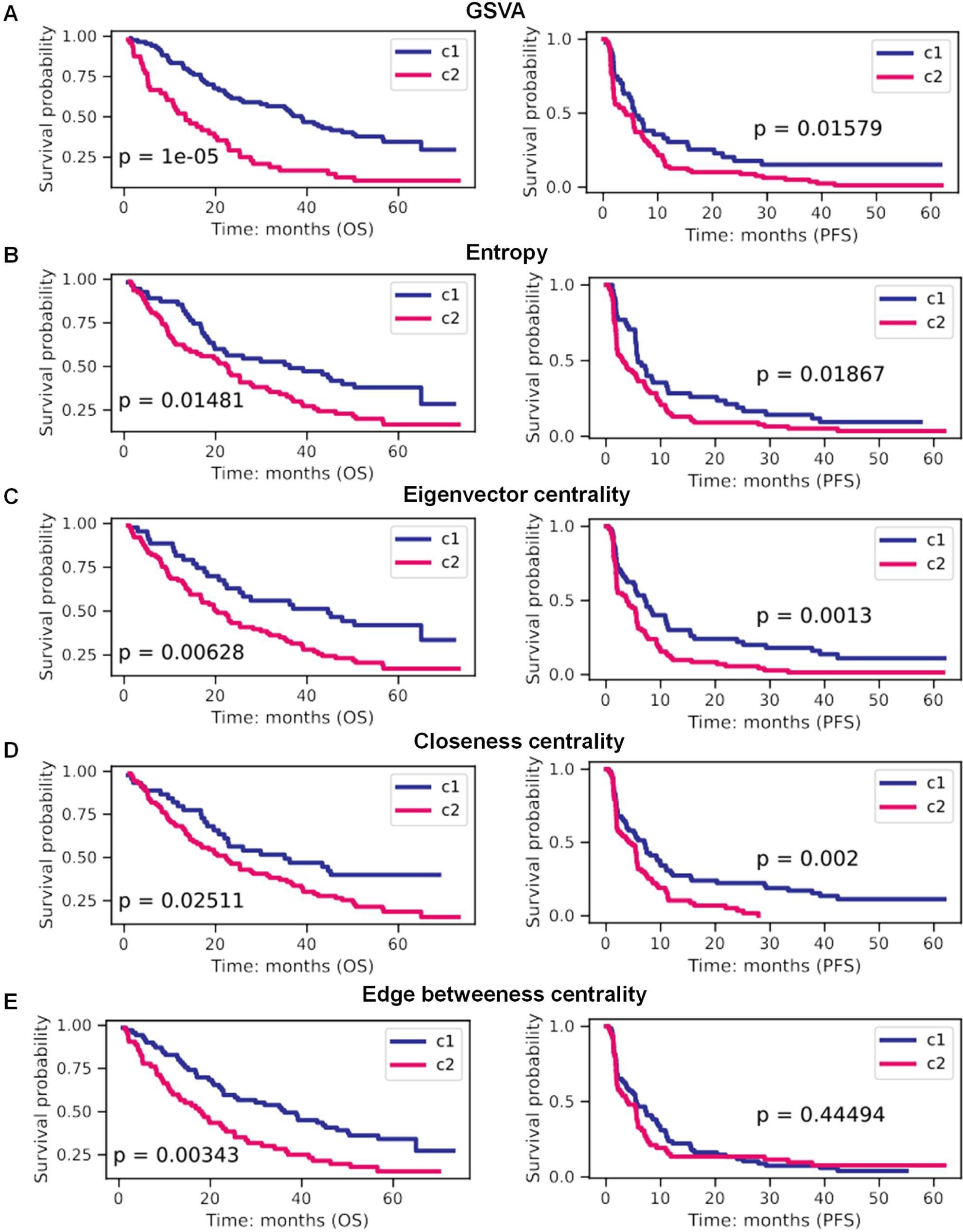
Survival analysis of biological pathway entropy and centrality scores using the subcohort of tumor primary sites from patients followed after immunotherapy by Nivolumab (pN). Several survival analyses were carried out with separation of the patient cohort according to topological pathway scores: (A). Use of 12 (OS) and 5 (PFS) significant pathways based on GSVA scores, (B). Use of 28 (OS) and 7 (PFS) significant pathways based on entropy scores, (C). Use of 8 (OS) and 10 (PFS) significant pathways based on gene eigenvector centrality scores, (D). Use of 16 (OS) and 3 (PFS) significant pathways based on gene closeness centrality scores, and (E). Use of 8 (OS) and 10 (PFS) significant pathways based on edge betweenness centrality scores. P values were calculated using log rank test.

Regarding the biological pathways involved, some were consistently associated with survival across multiple score categories, while others were identified by only one type of score. The phosphatidylinositol pathway, known to have a central role in ccRCC [46], was notably found by all scores, except GSVA. The ERBB signaling pathway, having a vital role in the initiation and progression of ccRCC [47], was identified by GSVA and the eigenvector centrality score. Regarding specific pathways, the eigenvector centrality score revealed some of the main pathways associated with the pathogenesis of kidney cancer such as mTOR, ascorbate and aldarate metabolism, unsaturated fatty acids, and mismatch repair [48–51]. Additionally, the circadian rhythm pathway linked to ccRCC prognosis was identified by the closeness centrality score [52].

The relevance of pathway entropy and centrality scores in predicting survival or recurrence was also confirmed in the mN, pE, and mE subcohorts (Supplementary Figure S19-20 and Details in Supplementary Note 6). Overall, these analyses of pathways deregulation, considering the perturbation of gene network topology, provide a highly complementary approach to the traditional methods based on direct gene expression analysis. Indeed, pathway entropy and centrality scores demonstrated strong predictive power for the prognosis and treatment response of ccRCC patients.

### Combining network features and gene expression values in machine learning models better predict immunotherapy response

Gene network information from existing databases can enhance survival predictions in cancer patients [53]. We set out to investigate whether sample-specific network features could improve the performance of gene expression-based machine learning (ML) models in predicting drug responses among immunotherapy-treated patients (Figure 6A). For this analysis, we focused on 90 samples from the pN subcohort, which were categorized as either CB or NCB. Using expression values of 64 genes selected by the Cox model, we trained the logistic regression ML model with the training set of 70% samples allocated, and evaluated its prediction performance with the test set. The area under the curve (AUC) of the receiver operating characteristics curve was used as a performance metric. The gene expression-based reference model yielded an AUC of 0.73 (Figure 6B, blue curve). In contrast, network feature-based ML showed comparable or better prediction performance (Figure 6C). Notably, combining gene expression data with network features enhanced the performance of gene expression-based model. For example, using 64 genes combined with previously identified 51 edges improved the AUC to 0.83. Additionally, the ML model using sample-level pathway entropy exhibited a higher AUC (0.73) compared to the gene set variation analysis (GSVA) pathway-based ML model, which had an AUC of 0.69. The combination of pathway entropy with GSVA scores also resulted in a better AUC (0.72). Besides, we validated the prediction performance of ML models with different algorithms using Leave One Out Cross Validation and found that network features combined with gene expression ML models had higher AUC than gene expression-based ML models (Supplementary Figure S21). These results suggest that integrating sample-level network features with gene or pathway markers can enhance the performance of ML models in predicting ccRCC patient response to immunotherapy.

**Figure 6.**
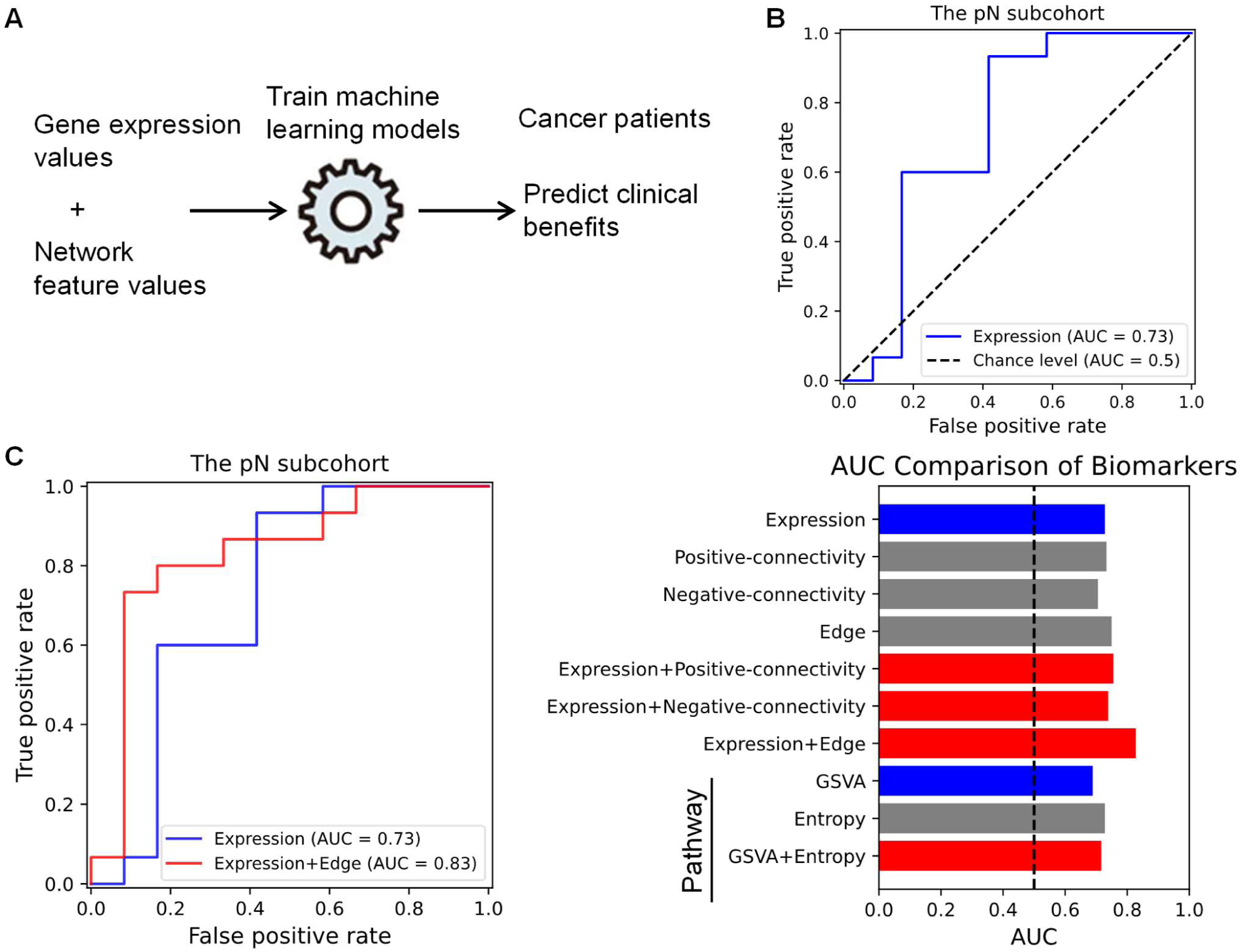
Prediction of drug response for immunotherapy-treated patients using gene expression values and network features. (A). Overall scheme of ML models using gene expression and network features to predict treatment response for cancer patients in the pN subcohort. (B). The area under the receiver operating characteristic curve (AUC) of a logistic regression ML model based on gene expression value as the input. (C). The ML performance: AUC using gene expression (blue), or network features (grey), or pathway scores (blue), or the combination between gene expression and network features (red). The pN subcohort (n=90) used here is composed of 44 patients with clinical benefits and 46 patients without clinical benefits. To calculate AUC, the dataset is split into a train set and a test set with 30% of sample size. The ML models were trained with the train set and evaluated with the test set.

### Validation of pathway entropy and centrality scores in another cohort

Our sample-specific pathway scores having shown their ability to reveal the susceptibility of patients to nivolumab, we extended this study to an independent cohort of 354 ccRCC patients followed after treatment with the therapeutic combination avelumab and axitinib (immune checkpoint inhibitor and anti-angiogenic) [19]. We therefore inferred sample-specific gene networks, as performed with the Braun et al data, and calculated pathway scores for clustering samples into two groups (Supplementary Figure S21). Respectively with eigenvector, closeness, and edge betweenness scores, the response for this treatment was found to be significantly different between two clusters generated using 10, 5, or 6 significant pathways (Figure 7, Supplementary Figure S22). These pathways were already known to be deregulated in ccRCC such as those associated with the metabolism of amino acids and fatty acids or those involved with DNA repair [54–57]. While no significance of survival was found with pathway entropy, the consistent divergence of metabolism pathways was found between clusters suggesting their role in the progression of ccRCC (Supplementary Figure S22).

**Figure 7.**
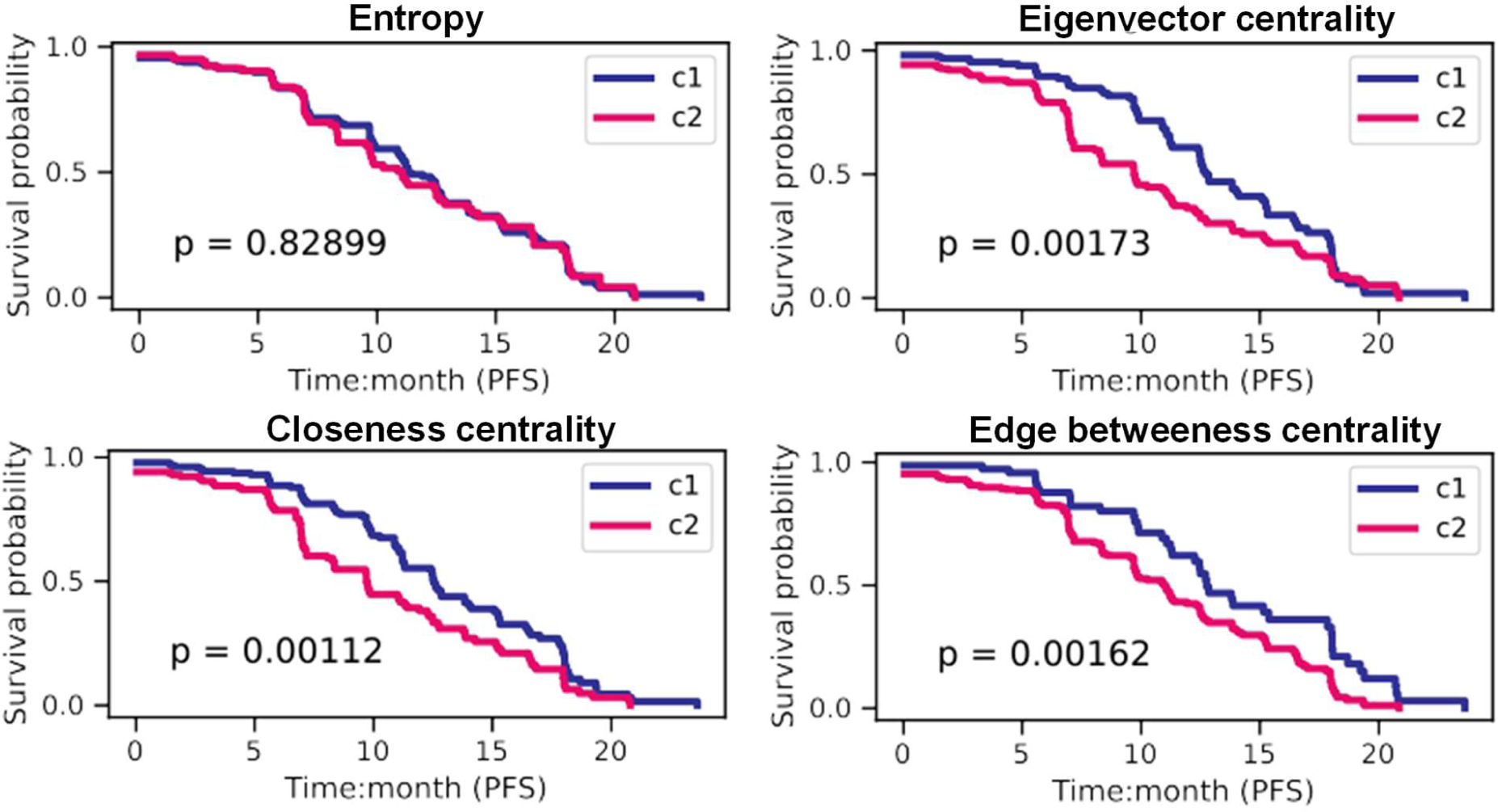
Survival analysis using biological pathway entropy and centrality scores calculated using an independent patient cohort from Mozter et al. Clusters c1 (blue) and c2 (pink) of samples were defined using an unsupervised hierarchical clustering of pathway scores significantly associated with patient PFS values. P values were calculated using the log rank test.

To conclude, we confirmed on an independent cohort of ccRCC patients followed post-treatment by a combination involving immunotherapy, the relevance of pathway scores based on the topology of ssGCNs to classify patients according to response to treatment.

## Discussion

Detection of regulatory perturbations in sample-specific gene networks inferred from expression data has contributed to progress in personalized medicine by refining the stratification of patients into cancer subtypes [12, 14, 15, 38, 58]. Our study introduced an innovative computational framework for sample-specific network analysis, facilitating the use of multiscale network features in cancer research. Our approach enabled a more refined characterization of gene networks for ccRCC patients, and revealed distinct gene coexpression patterns linked to immunotherapy response.

Network similarity facilitates the comparison of gene networks, allowing for the clustering of networks that group patients with similar gene regulation patterns. While network distance has been effective in identifying tumor types in Partial correlation-based networks [12], we detected a lack of sensitivity when applying it to Pearson correlation-based networks to classify patients based on treatment responses. To enhance its efficacy, we improved network distance by comparing one sample to a selected group of patients with known medical outcomes. Our adjusted network distance revealed a high-resolution way to associate each sample-specific co-expression network with the patient’s clinical outcome.

Specific network markers such as node degree and edge weight further showed that higher gene connectivity and stronger negative gene pairwise associations were prevalent in patients with poorer survival probability and worse treatment response. The biological significance of these network markers was substantiated by previous published works, which reported that many genes revealed by our study were involved in tumor progression, metastasis, and anti-tumor treatment response. Furthermore, the change of these network features was associated with deletion_11q12.3 and deletion 11q23.1, and intratumor heterogeneity for the subcohort pN, which were previously linked with cancer progression and prognosis [33–35]. The enrichment of identified genes related to natural killer (NK) T cells suggests the role of this cell type in the response of patients to nivolumab, in line with that gene markers and the population of NK cells were reported to be associated with prognosis and clinical outcomes in ccRCC [36, 37]. These signatures of network features were different from frequently used differentially expressed genes, suggesting novel and complementary predictors of treatment response in ccRCC patients.

The topology of gene coexpression networks revealed by varied gene connectivity and edges could be also observed through the lens of biological pathway networks. Entropy measures of signaling pathways have been already proposed to assess their activation state for survival analysis in pan-cancer studies [59]. We calculated our entropy and topology-based pathways scores as well as a classic pathway score GSVA, at a single-sample level. This approach allowed us to uncover biological pathways whose coexpression network perturbations were significantly associated with patient survival or cancer recurrence after treatment. While some pathways were revealed in common by several scores, others were specific to a score, illustrating the strong complementary of these approaches. Concerning the identification of deregulated pathways associated with response to nivolumab, we revealed the relevance of the eigenvector centrality score for samples from primary and metastatic sites of ccRCC, supplemented by the closeness and the edge betweenness centrality scores for metastases. Our scores provide various options to detect significant pathways with entropy or topological changes to decipher the divergence of patients’ clinical benefits, serving as an alternative in sample-wise gene set enrichment analysis.

Also, these network features demonstrated their power in predicting patient’s clinical benefits and further improved the prediction performance of gene expression-based ML models. Specifically, edges increased the performance of gene-expression based ML model from 0.73 to 0.83. Although our pathway entropy increased slightly the performance of GSVA-based ML model due to the small number of selected pathways, the predictive ability of our pathway scores were confirmed using another independent dataset. Consistent with other network-based prediction models [60, 61], our results suggest that combing sample specific network features with gene expression can better predict patient response to immunotherapy.

In terms of potential novel predictors of treatment response in ccRCC patients, we detected a large discrepancy in these lists of genes carrying co-expression patterns between primary and metastatic sites which was consistent with previous studies that expression profiles changed significantly between primary tumors site and metastases [62]. Furthermore, through pathway network analysis, we revealed that the deregulations of the ccRCC primary site associated with PFS values were rather captured by the phosphatidyl inositol, ERBB and mTOR signaling pathways, and by the ascorbate and aldarate metabolism and the mismatch repair system, while those of metastases were linked to cytokine inflammation, sphingolipid metabolism, amino-acyl-Trna biosynthesis and Citrate-TCA cycle pathways, as well as the nucleotide excision process. These findings underscore the distinct co-expression patterns and biological pathways that are differentially associated with patient survival depending on whether the sample is from a primary or metastatic site. This specificity suggests that the biological mechanisms driving patient outcomes in ccRCC can vary significantly depending on the tumor site location, highlighting the importance of considering the biopsie or surgery site in the development of predictive models and therapeutic strategies.

Our approach has several limitations and perspectives. The sample size of clinical cohorts is limited and gene coexpression patterns observed from our analysis need to be validated in other independent cohorts for different cancers. Moreover, our approach may require the control of the heterogeneity of samples in a group to maintain common edges for identifying specific edges. Lastly, the biological meaning of coexpression patterns needs to be further defined. The change in co-expression networks from bulk RNA-seq data may be influenced by the cellular content of the tumor immune microenvironment, tumor heterogeneity, or intercellular communication. The advent of single-cell RNA-seq datasets from patient tumors may provide an opportunity to explore network change at the scale of cellular subtypes to unravel intracellular factors from those in the microenvironment sources driving changes in coexpression.

## Conclusion

In conclusion, sample-specific gene coexpression network features demonstrate coexpression patterns as alternative markers to predict survival and treatment response of patients with advanced kidney cancer. Our computational framework for investigating gene network features linked to patient treatment responses is useful to support the personalization of therapies in the clinic.

## Key Points

- We developed a sample-specific gene network approach to predict the immunotherapy response of kidney cancer patients.
- Higher gene connectivity and stronger negative gene-gene associations are pivotal factors in immunotherapy-treated patients with poor prognoses.
- Network features, predictive of patients’ treatment response, were different between primary tumor site and metastases.
- Sample-specific pathway entropy and topology scores were complementary to conventional sample-wise pathway enrichment scores.

## List of abbreviations

AUC: The area under the curve of the receiver operating characteristics curve
CB: Clinical benefits
ccRCC: Clear cell renal cell carcinoma
GCN: Gene coexpression network
GSVA: Gene set variation analysis
ICI: Immune checkpoint inhibitors
KEGG: Kyoto Encyclopedia of Genes and Genomes
MSigDB: Molecular Signatures Database
mE: Metastases patients treated with everolimus
mN: Metastases of patients treated with nivolumab
NCB: Non-clinical benefits NCB
Nd: Network distance
pE: Primary tumor site of patients treated with everolimus
PCC: Pearson correlation coefficient
pN: Primary tumor site of patients treated with nivolumab
ssGCN: Sample-specific gene coexpression network
TF: Transcription factor
TPM: Transcript per million

## Declarations

### Ethics approval and consent to participate

Not applicable.

### Consent for publication

Not applicable.

### Availability of data and material

The code for these analyses in this manuscript is available in the GitHub repository (https://github.com/liangwei01/ssGCA). Data are incorporated into the article and its online supplementary material. Supplementary filea are available Online.

### Authors’ contributions

LY and CB conceived the study. LY developed and performed the analysis. LY and CB wrote the manuscript. PT contributes to the methodologies. NMT, ME, and YM contributed to the revision of the manuscript.

## Acknowledgments

Part of the computations were carried out using the GRICAD infrastructure (https://gricad.univ-grenoble-alpes.fr), which is supported by Grenoble research communities. The authors also appreciate the valuable suggestions of Dr. Guilherme Ferraz de Arruda and Dr. Alberto Aleta on the method.

