## Supplemental Note for "Sample-specific network analysis identifies gene coexpression patterns of immunotherapy response in advanced kidney cancer"

**Supplementary Information**

Liangwei Yin^1^, Pietro Traversa^2,3,4^, Mohamed Elati^5^, Yamir Moreno^2,3,4^, Natalia Marek-Trzonkowska^6,7^, Christophe Battail^1, *^

^1^ Université Grenoble Alpes, IRIG, Laboratoire Biosciences et Bioingénierie pour la Santé, UA 13 INSERM-CEA-UGA, 38000 Grenoble, France.

^2^ Institute for Biocomputation and Physics of Complex Systems (BIFI), University of Zaragoza, 50018 Zaragoza, Spain

^3^ Department of Theoretical Physics, University of Zaragoza, 50018 Zaragoza, Spain

^4^ CENTAI Institute, 10138 Turin, Italy

^5^ Univ. Lille, CNRS, Inserm, CHU Lille, UMR9020-U1277 – CANTHER – Cancer Heterogeneity Plasticity and Resistance to Therapies, Lille F-59000, France

^6^ International Centre for Cancer Vaccine Science, University of Gdansk, Kladki 24, 80-822, Gdansk, Poland

^7^ Laboratory of Immunoregulation and Cellular Therapies, Department of Family Medicine, Medical University of Gdańsk, ul. Dębinki 2, 80-811 Gdańsk, Poland

**Supplementary Note 1. Sample specific gene network inference**

Several methods have been developed to infer sample-specific GCNs (ssGCN), such as sample-specific networks using one sample against a group of given control samples (SSNs) [1], single networks using linear interpolation of single sample against its corresponding group (LIONESS) [2], sample networks inferred with the genome wide sample-to-sample correlation (SWEET) [3], single networks based on the partial correlations between genes (P-SSN) [4], and single networks based on gene rank (Edge perturbation) [5]. Recently reported, the SWEET method may overcome the effect of sample size and achieve the best performance on the construction of sample networks [3].

In this study, we used the recently developed method (SWEET) to construct single-sample weighted GCN. An aggregated network (N_ij_^G^) was first constructed using gene expression of all samples within a category and a perturbed network (N_ij_^G-S^) was then constructed after popping out one specific sample (Figure 1A). Pearson correlation coefficient (PCC) was used as gene-gene association (i.e. edge weight). Sample-specific networks were estimated by the difference in edge weight between the aggregated network and each perturbed network, and they were added to the perturbed network to mimic the cohort network. To eliminate the noise inside networks, the significance level of confidence scores of edges was assessed by the z-score normalization method using a z score at 2.58 which is 0.01 of the two-sided p-value. If the significance level was less than 2.58, the edges would be considered as noise and eliminated in our networks. Cohort level GCNs were also constructed using the z-score method above for the group of samples with clinical benefits (CB) or non-clinical benefits (NCB).

**Supplementary Note 2. Selection of network features and its clustering**

To reduce the computation time, network features were selected based on their variance across all samples in their subcohort. Furthermore, genes, edges and pathway scores were filtered by applying the univariate Cox regression model (p-values < 0.01). Hence the matrix of network features was composed of samples and their network features (gene connectivity or edge weight). For networks from the negative association between genes, they were transformed to their absolute values for further analysis.

Unsupervised hierarchical clustering was performed using the ward method and cosine metric with gene connectivity, edge weights or pathway scores as inputs. To facilitate comparisons of treatment responses and of other clinical features, we chose to separate the samples into two clusters. Note that irrelevant pathways were filtered out for the clustering step in pathway analysis.

**Supplementary Note 3. Pathway centrality.**

Sample-specific pathway networks were subtracted from sample networks using gene sets of pathways from the KEGG database [6]. To enable the calculation of pathway scores, both positive and negative values (converted to their absolute values) of edges were considered. Pathway topological scores were calculated based on the average of gene eigenvector centrality, gene closeness centrality, and edge betweenness centrality. Gene eigenvector centrality refers to the influence of nodes within a network. Its high score indicates that one node is connected with many nodes with high scores. Gene closeness centrality was used to detect nodes that had the shortest distance to other nodes and its high score means one node is closest to the others. Edge betweenness centrality indicates the importance of an edge on the paths in a network, and its high score indicates that an edge lies on many shortest paths. Centrality scores were calculated using the functions from the Python igraph package.

**Supplementary Note 4. Gene connectivity is associated with treatment response**

Based on gene connectivity from positive correlations, 21 genes were selected because of the significant association of their connectivity with OS and PFS survival data in the pN subcohort (Figure 3A, p-value < 0.01; Table 2). Among these genes, some were already known to be involved in the progression of ccRCC or other cancers or in tumor immunity. NLRC5 mediates cell proliferation, migration and invasion in ccRCC through the Wnt/beta-catenin pathway and plays a key role in cancer immune surveillance [7, 8]. PLCB3 is involved in ccRCC proliferation with its association with SHP-1 and being a prognostic marker for lung cancer [9, 10]. Finally, COPG2 is considered to be an oncogene, and LRCH4 is involved in leukocyte biology [11, 12].

A similar analysis of gene connectivity from negative correlations identified 48 genes significantly associated with both PFS and OS. In terms of expression values, only 5 of them were associated with survival data (Supplementary Figure S7A, p-value < 0.01). Unsupervised clustering was also carried out from these 48 genes to classify the samples into two groups, with cluster 2 associated with greater gene connectivity (Supplementary Figure S7B). We then studied the relevance of the separation of primary site tumor tissues according to patient survival and found that cluster 2, with stronger gene connectivity, was associated with significantly lower PFS and OS values than cluster 1 (Supplementary Figure S7C). We finally found that the chromosomal loss 11q23.1 was also significantly enriched in cluster 2 (Supplementary Figure S7D). These genes were enriched in similar pathways and gene sets as showed from positive association and a bibliographic study of these genes notably revealed that some of them were already associated with metastasis and poor prognosis for various cancers. ACIN1 was found to be up-regulated in pro-metastatic B cells in ccRCC and correlated to a poor prognosis [13]. The over-expression of IFN2 was associated with tumor progression and metastasis in glioblastoma, triple-negative breast cancer and gastric cancer [14–16]. SNED1 promotes breast cancer metastasis [17]. MYO9B, SYPL1 and BICDL1 were already linked to poor prognosis in several cancers [18–20].

From the results of the gene connectivity of primary tumor sites, we then studied whether they were preserved for tumor metastases by exploring the mN subcohort. We therefore selected 9 and 17 genes whose connectivity, based on positive and negative associations, were significantly associated with OS and PFS survival values (Table 2). Out of them, the expression values of only 1, and 6 genes were associated with survival data. Interestingly, no overlap was identified between these genes and those previously identified from the pN subcohort based on gene connectivity. Unsupervised clustering split the samples into two groups, with cluster 2 associated with higher gene connectivity, and significantly lower PFS and OS values than cluster 1 (Supplementary Figure S8AB.1&2). Cluster 2 was also associated with significantly higher ITH (Supplementary Figure S8B.4). Although no pathway enrichment was found, several of these genes (*GMNN*, *STON1*, *SUSP1*) were already known to be associated with the prognosis or treatment response of ccRCC patients [21–23].

**Supplementary Note 5. Highly negative gene-gene associations in patients without clinical benefits**

For the pN subcohort, a non-exhaustive bibliographic study of these gene from edges has shown the selection of known cancer genes. Concerning PRELID3B–TTLL3 edge, PRELID3B was proposed as a prognostic and immunological marker in pan-human cancer [24], while TTLL3 was shown to carry prognostic value in ccRCC [25]. For BMPR2–MAN2C1 edge, BMPR2 was identified as a driver gene in ccRCC and MAN2C1 inhibited proliferation and invasion in ccRCC [26–28]. Regarding IRF3–ZBTB10 edge, IRF3 was linked with poor prognosis in ccRCC and ZBTB10 was implicated in tumor growth and metastasis in breast cancer [29, 30].

We then studied edges from ssGCNs of the mN subcohort and identified 6 edges significantly associated with both OS and PFS values (Table 3). An unsupervised classification based on the 6 edges highlighted two clusters, with cluster 2 enriched in patients associated with significantly shorter OS and PFS for nivolumab treatment (Supplemental Figure S13AB). Similarly, 5 of these edges were highly co-expressed in NCB patients compared with CB patients (Supplemental Figure S13C). Next, a gene ontology analysis revealed the enrichment in molecular processes (Myc targets, ribosome) and the NK T cell population (Supplemental Figure S13D). Interestingly, almost all of the genes of edges were already known to be involved in cancer progression and metastasis [31–38].

**Supplementary Note 6. Pathway entropy and centrality scores**

A similar analysis for the mN subcohort clustered patients based on pathway scores. We found greater abilities to differentiate patients according to OS values for the entropy and topology scores of pathways, compared to GSVA (Supplementary Figure S18). Concerning the association of pathway scores with PFS values, the best performances were obtained using the eigenvector, closeness and edge betweenness centrality scores. Among the pathways jointly found by several scores, the nucleotide excision process, linked to the ccRCC therapeutic response [39], was identified by both GSVA and edge betweenness centrality score. As for specific pathways, the GSVA method identified the cytokine inflammation, involved in the metastatic progression of ccRCC [40]. The pathways of sphingolipid metabolism, associated with the prognosis of ccRCC metastasis, and of amino-acyl-Trna biosynthesis, activated in high-grade ccRCC [41] were found by the entropy score. Additionally, the eigenvector centrality score identified the Citrate-TCA cycle pathway suggested to be activated in metastatic ccRCC [42].
