## Supplemental Figure for "Sample-specific network analysis identifies gene coexpression patterns of immunotherapy response in advanced kidney cancer"

Tables:

| Cancer | Localization | Treatment | No. of tumor samples |
|---|---|---|---|
| Advanced ccRCC | primary | Nivolumab | 133 |
| Advanced ccRCC | primary | Everolimus | 92 |
| Advanced ccRCC | metastasis | Nivolumab | 47 |
| Advanced ccRCC | metastasis | Everolimus | 37 |
| ccRCC | unknow | Avelumab and Axitinib | 354 |
| Normal kidney tissue | Cortex | NA | 85 |

Table 1. We collected RNA-seq data from several studies,
Including the Braun 2020 paper, the Motzer 2020 paper and the GTEx portal .

| | | Number of prognostic genes based on gene connectivity | | | | | | |
|---|---|---|---|---|---|---|---|---|
| subcohort | | OS | PFS | Overlapped | | OS | PFS | Overlapped |
| pN | Positive correlation | 218 | 85 | 21 | Negative correlation | 209 | 108 | 48 |
| pE | | 74 | 97 | 29 | | 114 | 131 | 39 |
| mN | | 72 | 114 | 9 | | 100 | 165 | 17 |
| mE | | 69 | 17 | 1 | | 67 | 11 | 0 |

Table 2.Genes were selected by cox regression model, and p value at 0.01 was applied.
To reduce computation time, only top 5000 variable genes were explored.

| subcohort | | Number of prognostic genes based on edges | | |
|---|---|---|---|---|
| | | OS | PFS | Overlapped |
| pN | Edges | 214 | 224 | 51 |
| pE | | 228 | 165 | 40 |
| mN | | 85 | 100 | 6 |
| mE | | 70 | 84 | 10 |

Table 3. Genes were selected by cox regression model, and p value at 0.01 was applied.

| Query Name | Rank | TF | Score | Library | Overlapping_Genes |
|---|---|---|---|---|---|
| gene_set_query | 1 | MZF1 | 6 | ssion,1;Enrichr Queries,9;GTE | 6L2,TTLL3,TAF1C,CROCC,SGSM3,TUBGCP6,TNRC18,CAMTA2,MA |
| gene_set_query | 2 | ZNF692 | 9.667 | ssion,6;Enrichr Queries,1;GTE | 2,ATG16L2,TAF1C,CROCC,SGSM3,TUBGCP6,TNRC18,CLASRP,MA |
| gene_set_query | 3 | ZNF76 | 15.67 | sion,19;Enrichr Queries,25;GT | LL3,TAF1C,CROCC,SGSM3,TUBGCP6,TNRC18,CAMTA2,MAN2C1, |
| gene_set_query | 4 | CXXC1 | 18.67 | sion,29;Enrichr Queries,17;GT | NF2,IRF3,TAF1C,CROCC,SGSM3,CDC37,TUBGCP6,CLASRP,MAN2 |
| gene_set_query | 5 | ZNF316 | 20.33 | sion,24;Enrichr Queries,12;GT | RF3,ATG16L2,TAF1C,CROCC,TUBGCP6,TNRC18,CLASRP,MAN2C1 |
| gene_set_query | 6 | RBCK1 | 21 | Coexpression,35;GTEx Coexp | ,RHOT2,INF2,IRF3,ATG16L2,SGSM3,TUBGCP6,TNRC18,CLASRP, |
| gene_set_query | 7 | PRR12 | 23.33 | sion,27;Enrichr Queries,11;GT | 3,PKD1,RHOT2,INF2,CROCC,TUBGCP6,TNRC18,CAMTA2,CLASRP |
| gene_set_query | 8 | SCX | 25 | ARCHS4 Coexpression,25 | F1C,KMT2B,SCRIB,TNRC18,CLASRP,PPP1R12C,RGL2 |
| gene_set_query | 9 | MBD6 | 29.5 | Coexpression,38;GTEx Coexp | ABTB1,RHOT2,ATG16L2,TUBGCP6,TNRC18,MAN2C1,CLASRP,RGL |
| gene_set_query | 10 | ZNF783 | 31 | sion,46;Enrichr Queries,29;GT | F3,TTLL3,ATG16L2,TAF1C,CROCC,TUBGCP6,TNRC18,MAN2C1,CL |
| gene_set_query | 11 | SAFB2 | 33.5 | Coexpression,4;GTEx Coexpr | TTLL3,TAF1C,CROCC,TUBGCP6,TNRC18,CLASRP,MAN2C1,NLRP1 |
| gene_set_query | 12 | ANKZF1 | 35.67 | ssion,8;Enrichr Queries,95;GT | G16L2,TTLL3,TAF1C,CROCC,SGSM3,TUBGCP6,CLASRP,MAN2C1, |
| gene_set_query | 13 | FLYWCH1 | 43 | Coexpression,3;GTEx Coexpr | ROCC,TUBGCP6,TNRC18,CAMTA2,MAN2C1,NLRP1,CLASRP,ARG |
| gene_set_query | 14 | E4F1 | 51.5 | S4 Coexpression,21;Enrichr Qu | C,CROCC,TUBGCP6,TNRC18,CAMTA2,CLASRP,MAN2C1,ERGIC2, |
| gene_set_query | 15 | IRF3 | 52.6 | P-seq,106;Enrichr Queries,71;R | 15,BBIP1,COMMD6,ABTB1,RPS27,ATG16L2,TTLL3,IRF3,EIF3L,SG |
| gene_set_query | 16 | CC2D1A | 54 | Coexpression,9;GTEx Coexpr | AF1C,CROCC,TUBGCP6,TNRC18,CAMTA2,MAN2C1,CLASRP,ERG |
| gene_set_query | 17 | ATF6B | 54 | Coexpression,31;GTEx Coexpr | TB1,IRF3,ATG16L2,TUBGCP6,TNRC18,MAN2C1,CLASRP,RGL2,PP |
| gene_set_query | 18 | HSF4 | 61.67 | sion,2;Enrichr Queries,171;GT | NF2,TAF1C,CROCC,SGSM3,TUBGCP6,TNRC18,CAMTA2,MAN2C1, |
| gene_set_query | 19 | KMT2B | 71.5 | Coexpression,18;ReMap ChIP | 2B,SCRIB,RHOT2,RPS27,INF2,TAF1C,TUBGCP6,TNRC18,MAN2C1, |

Table 4. The enrichment of transcription factor were obtained from online query on the website: https://maayanlab.cloud/chea3/#top.

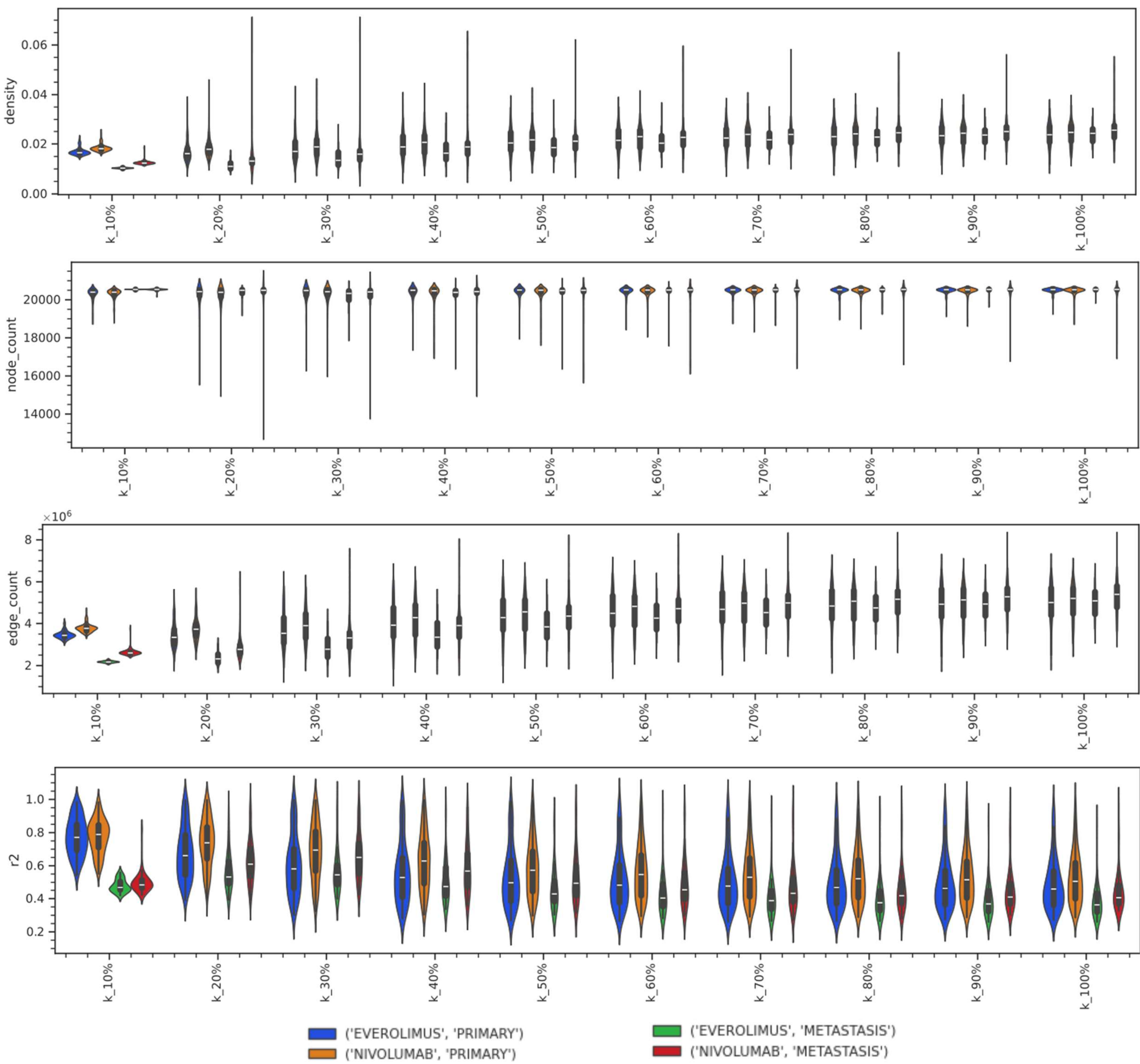


**Supplementary Figure S1. Network characteristics of ssGCNs constructed with different balance parameter k (from 0.1 to 1) in our four subcohorts**. When k was at 10%, ssGCNs achieved the best network density and determination coefficients $R^2$ of scale free topology, which appears to be more similar to realistic biological network. Specifically when k was set to 10%, ssGCNs of pE, pN, mE, nE achieved 1.68%, 1.83%, 1.03%, 1.25% network density, 20,273, 20,302, 20,538, 20302 node count, 3,458,841, 3,782,518, 2,176,807, 2,640,410 edge count and 0.772, 0.776, 0.477, 0.496 $R^2$ coefficients. The 10% k was used from all the downstream analysis.

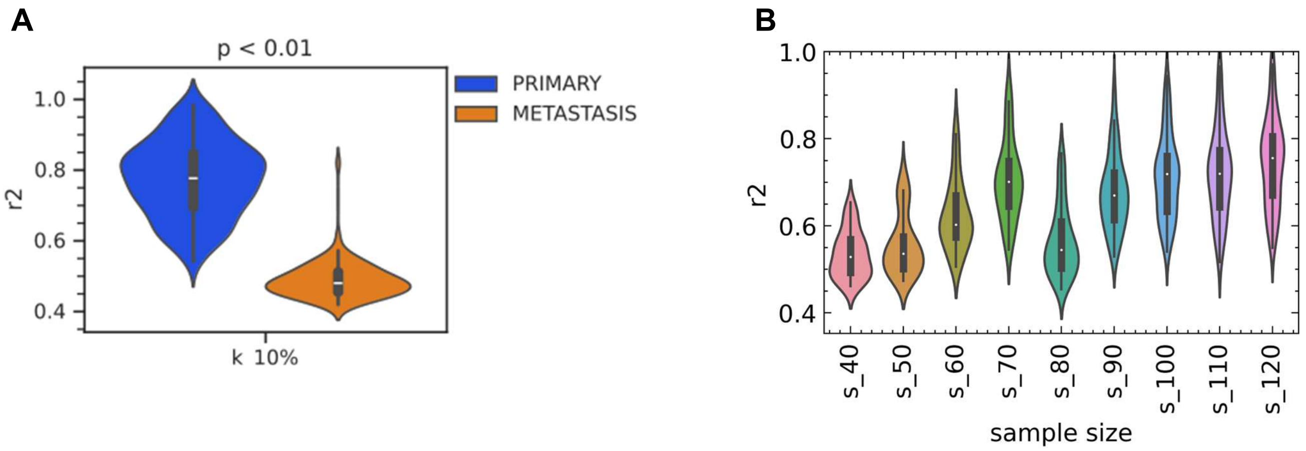


**Supplementary Figure S2. Analysis of the influence of cohort sample size on the scale free topology nature of networks using the pN group.** (A) The comparison of $R^2$ of ssGCNs between metastasis tumor in primary sites and metastasis. (B) The distribution of $R^2$ for ssGCNs constructed from different group size. The nature of scale free topology for networks was accessed by determination coefficient $R^2$. The closer $R^2$ is to 1, the degree distribution of a network follows a power law. Here, we used the biggest subcohort pN as the stimulation cohort to do the test. For different sample size, ssGCNs were constructed with a set number of randomly picked samples from pN and then $R^2$ of these ssGCNs was calculated. This figure provided that $R^2$ of network achieved higher values with larger group sample size.

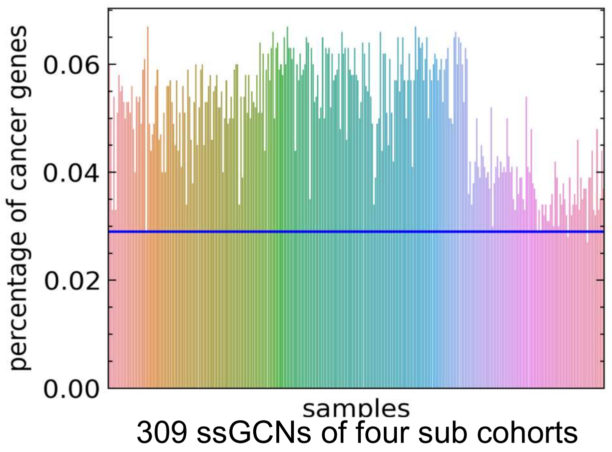


**Supplementary Figure S3. Enrichment analysis of cancer related gene in top 1000 high connectivity genes between normal samples and our tumor samples.** Cancer related genes were extracted from the Cancer Gene Census database. The blue line indicates the percentage (2.9%) of cancer related gene in the top 1000 nodes of kidney cortex network. Out of our 309 ssGCNs, 304 ssGCNs were found to have a higher enrichment of cancer related genes inside their networks.

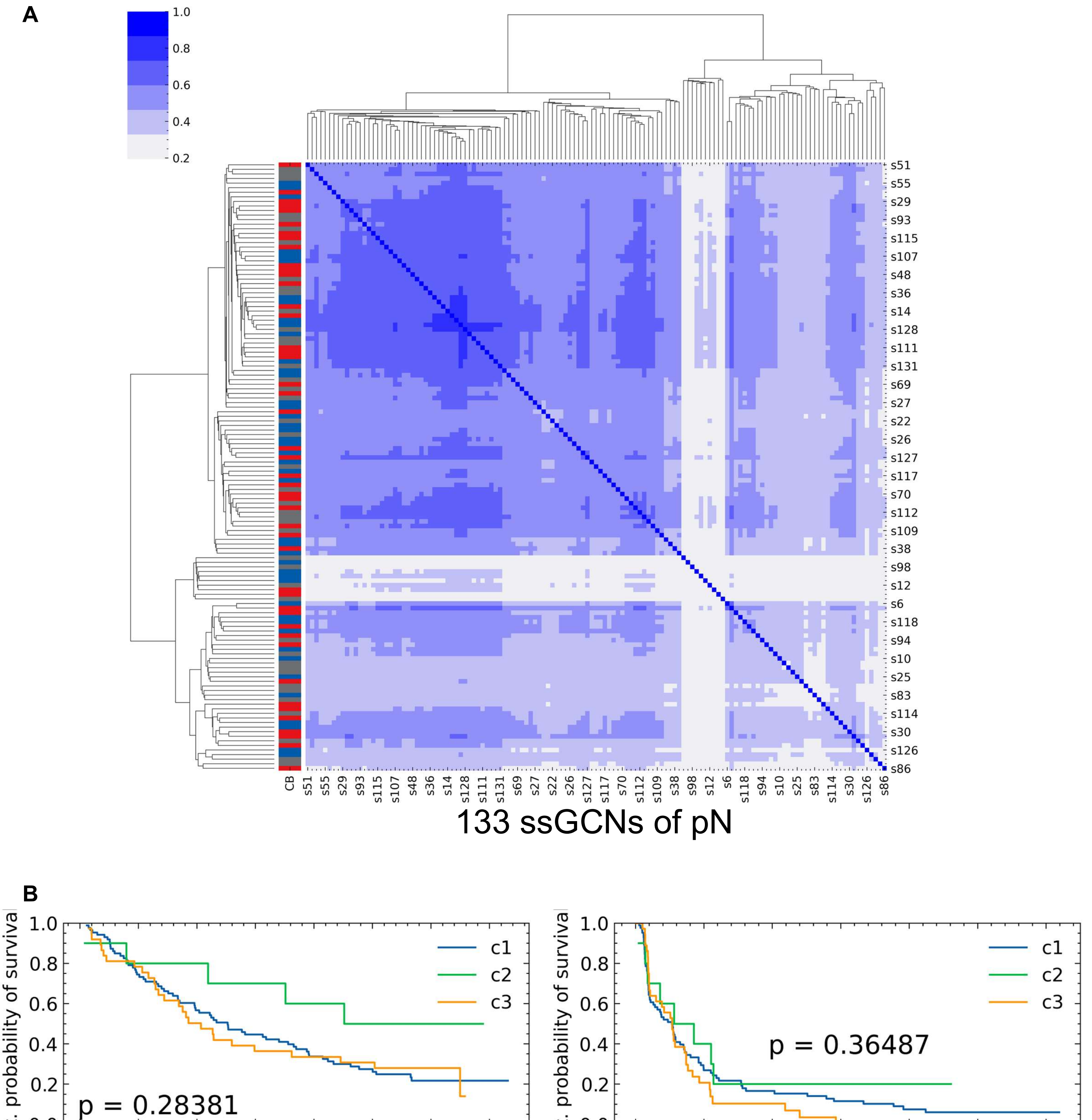


**Supplementary Figure S4. Patient clustering using network distance between samples of pN and survival analysis**. (A) Patient clustering of pN samples. Similarity matrix, calculated by network distance, was used as the basis of hierarchical clustering. (B) Survival analysis. Three clusters were inferred from the clustering result above and their survival probability were measured using the log rank test (p values were obtained here).

A

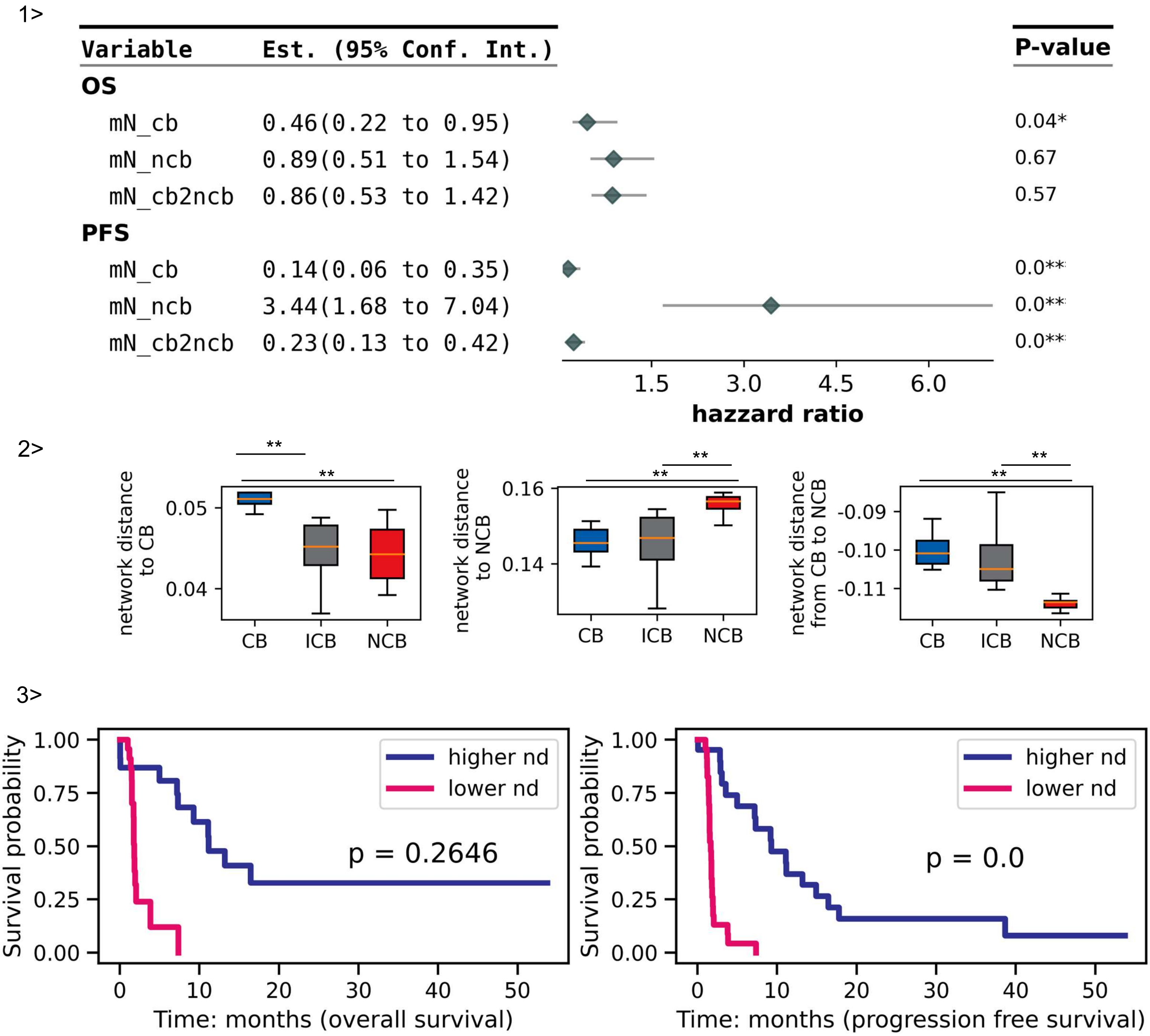
1>
Variable
Est. (95% Conf. Int.)
P-value
OS
mN_cb 0.46(0.22 to 0.95) 0.04*
mN_ncb 0.89(0.51 to 1.54) 0.67
mN_cb2ncb 0.86(0.53 to 1.42) 0.57
PFS
mN_cb 0.14(0.06 to 0.35) 0.0**
mN_ncb 3.44(1.68 to 7.04) 0.0**
mN_cb2ncb 0.23(0.13 to 0.42) 0.0**
1.5
3.0
4.5
6.0
hazzard ratio
2>
network distance to CB
0.05
0.04
network distance to NCB
0.16
0.14
network distance from CB to NCB
-0.09
-0.10
-0.11
**
CB
ICB
NCB
3>
Survival probability
1.00
0.75
0.50
0.25
0.00
higher nd
lower nd
p = 0.2646
p = 0.0
0
10
20
30
40
50
Time: months (overall survival)
Time: months (progression free survival)

B

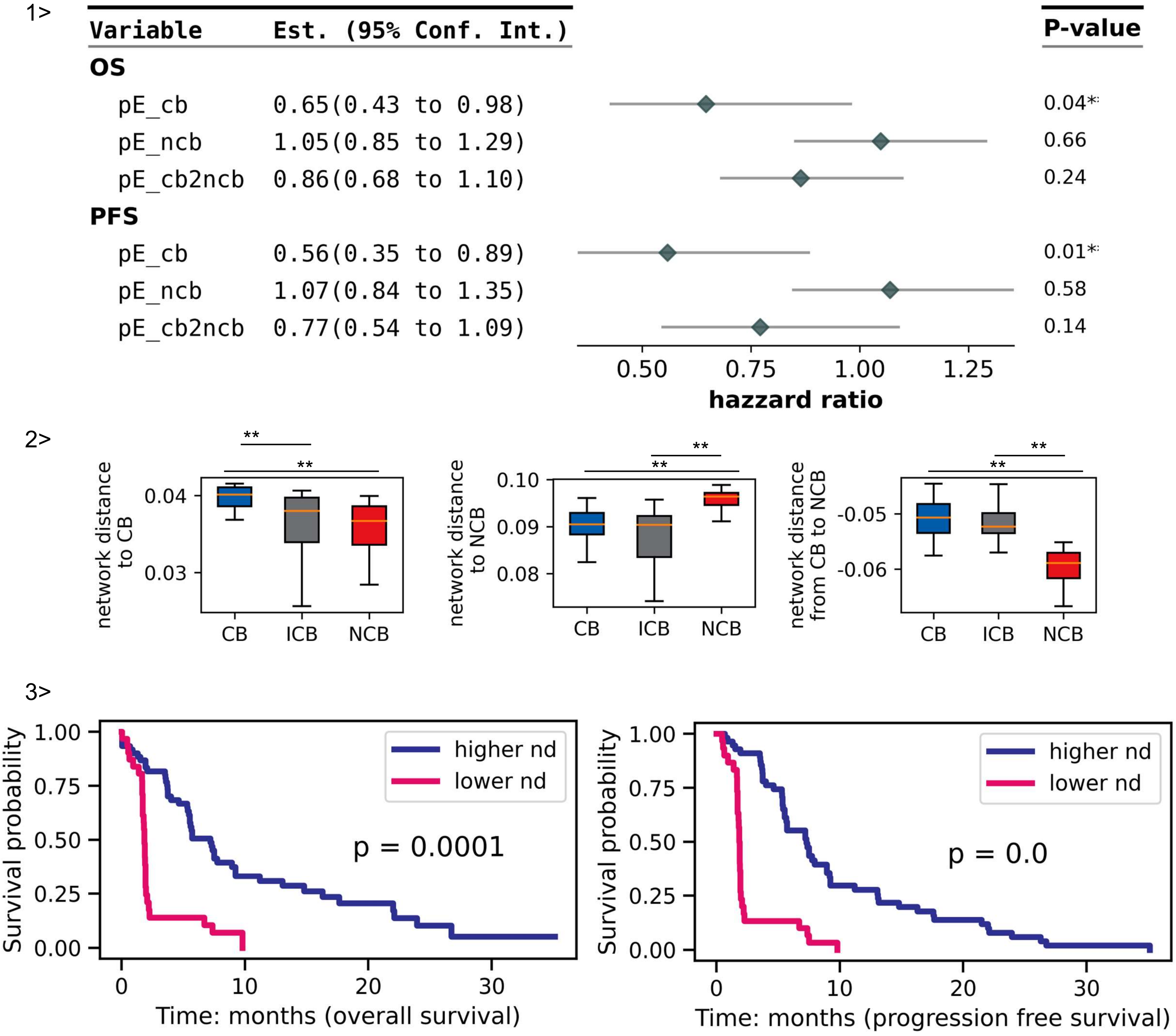
1>
Variable
Est. (95% Conf. Int.)
P-value
OS
pE_cb 0.65(0.43 to 0.98) 0.04*
pE_ncb 1.05(0.85 to 1.29) 0.66
pE_cb2ncb 0.86(0.68 to 1.10) 0.24
PFS
pE_cb 0.56(0.35 to 0.89) 0.01*
pE_ncb 1.07(0.84 to 1.35) 0.58
pE_cb2ncb 0.77(0.54 to 1.09) 0.14
0.50
0.75
1.00
1.25
hazzard ratio
2>
network distance to CB
0.04
0.03
**
**
CB
ICB
NCB
network distance to NCB
0.10
0.09
0.08
**
**
CB
ICB
NCB
network distance from CB to NCB
-0.05
-0.06
**
**
CB
ICB
NCB
3>
Survival probability
1.00
0.75
0.50
0.25
0.00
higher nd
lower nd
p = 0.0001
0
10
20
30
Time: months (overall survival)
Survival probability
1.00
0.75
0.50
0.25
0.00
higher nd
lower nd
p = 0.0
0
10
20
30
Time: months (progression free survival)

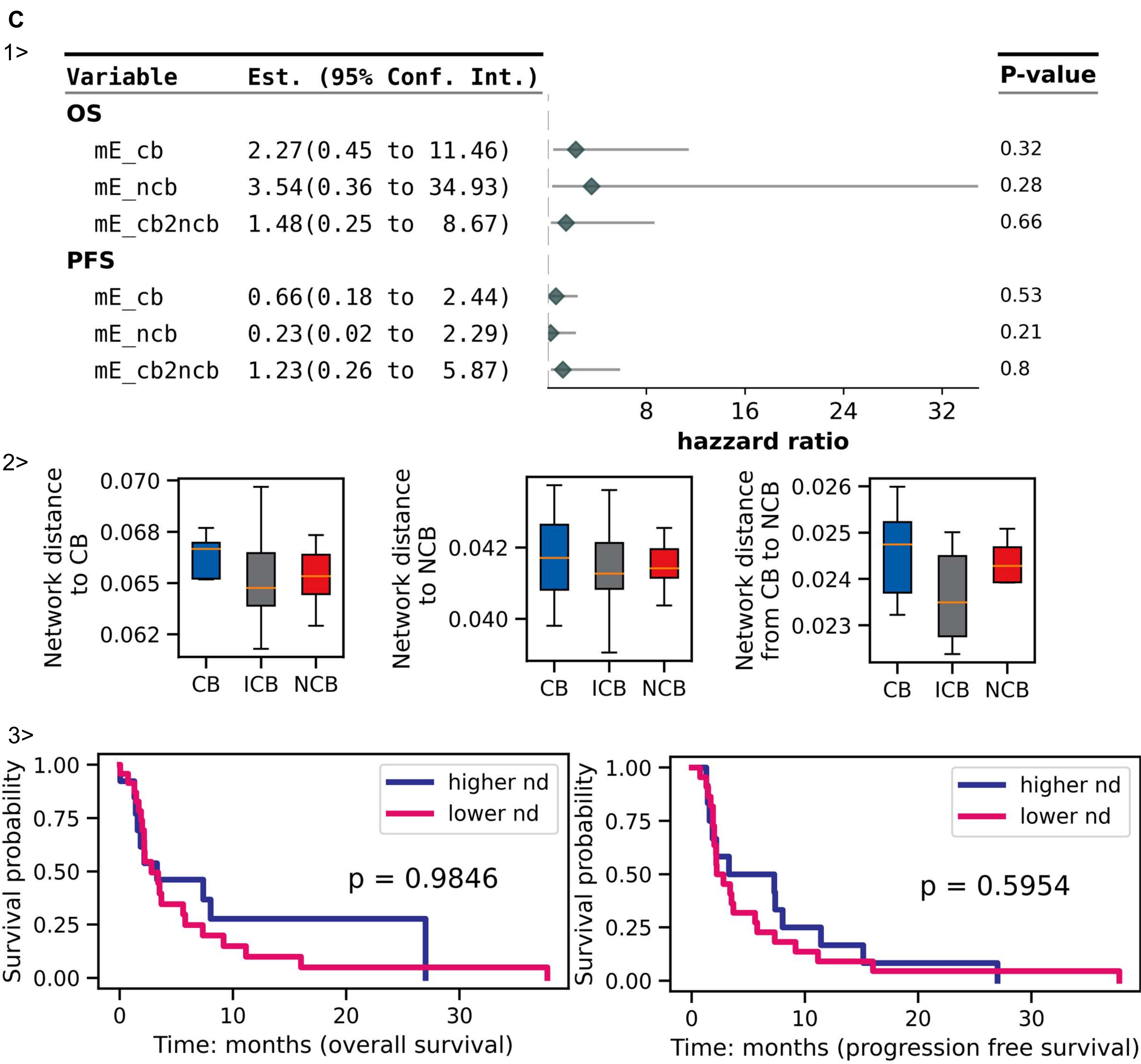


**Supplementary Figure S5. Adjusted network distance in mN(A), pE(B), mE(C)**.

1>. A forest plot of the univariate cox regression result using adjusted network distance. 2>. Comparison of adjusted network distance between samples with or without clinical benefits. Wilcoxon sum rank test was conducted. 3>. Survival analysis using network distance adjusted with CB to NCB. P value is from the log-rank test. Survival analysis using network distance adjusted with CB to NCB. Samples were divided into two groups (higher nd and lower nd groups) based on the median value of network distance adjusted with CB to NCB. P value is from the log-rank test. ( **: p value < 0.01; *: p value < 0.05)

Noted that mN, mE subcohorts have smaller sample size (47, 37), which may lead to that adjusted network distance was not effective here.

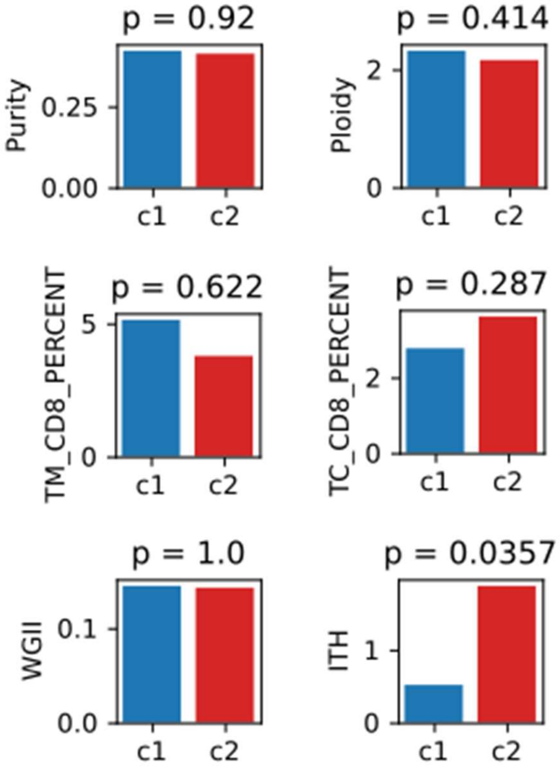


**Supplementary Figure S6. Comparison between two clusters generated from their positive correlation based gene connectivity of pN.** Wilcoxon rank sum test was conducted and p values below 0.05 were taken as significant. Cluster c1 has lower gene connectivity on average and significantly higher survival probability.

Clinical information of pN samples were retrieved from supplementary data of Braun 2020 paper.

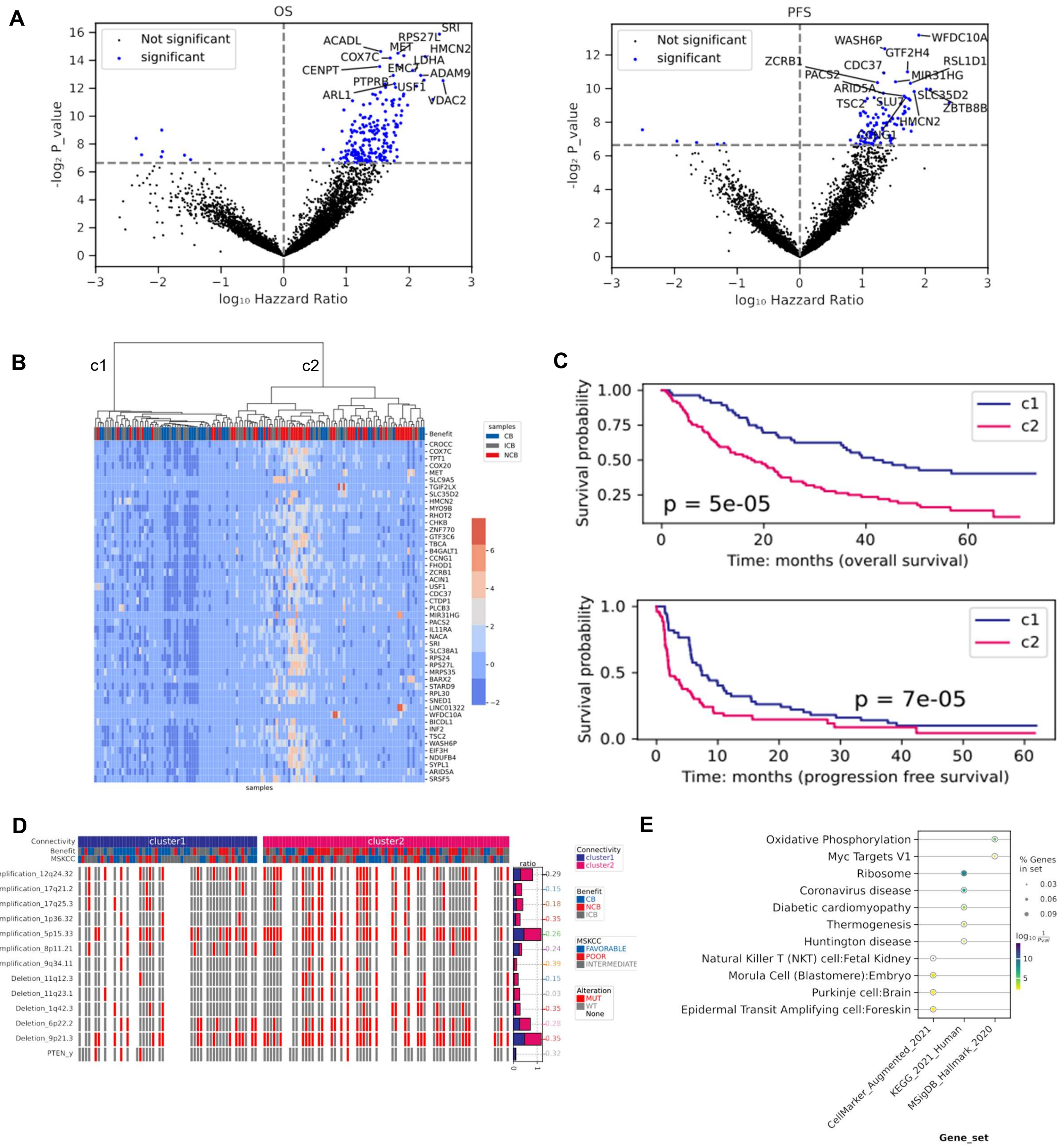

A
OS
Not significant
significant
SRI
RPS27L
ACADL
MET
HMCN2
COX7C
LDHA
CENPT
EMC7
ADAM9
PTPRB
USF1
ARL1
VDAC2
-log₂ P_value
log₁₀ Hazzard Ratio
PFS
WASH6P
WFDC10A
GTF2H4
ZCRB1
CDC37
RSL1D1
PACS2
MIR31HG
ARID5A
SLC35D2
TSC2
SLU7
ZBTB8B
HMCN2
CCNG1
B
c1
c2
Benefit
samples
CB
ICB
NCB
CROCC
COX7C
TPT1
COX20
MET
SLC9A5
TGIF2LX
SLC35D2
HMCN2
MYO9B
RHOT2
CHKB
ZNF770
GTF3C6
TBCA
B4GALT1
CCNG1
FHOD1
ZCRB1
ACIN1
USF1
CDC37
CTDP1
PLCB3
MIR31HG
PACS2
IL11RA
NACA
SRI
SLC38A1
RPS24
RPS27L
MRPS35
BARX2
STARD9
RPL30
SNED1
LINC01322
WFDC10A
BICDL1
INF2
TSC2
WASH6P
EIF3H
NDUFB4
SYPL1
ARID5A
SRSF5
C
Survival probability
c1
c2
p = 5e-05
Time: months (overall survival)
p = 7e-05
Time: months (progression free survival)
D
Connectivity
Benefit
MSKCC
cluster1
cluster2
ratio
Amplification_12q24.32
Amplification_17q21.2
Amplification_17q25.3
Amplification_1p36.32
Amplification_5p15.33
Amplification_8p11.21
Amplification_9q34.11
Deletion_11q12.3
Deletion_11q23.1
Deletion_1q42.3
Deletion_6p22.2
Deletion_9p21.3
PTEN_y
0.29
0.15
0.18
0.35
0.26
0.24
0.39
0.15
0.03
0.35
0.28
0.35
0.32
CB
NCB
ICB
FAVORABLE
POOR
INTERMEDIATE
Alteration
MUT
WT
None
E
Oxidative Phosphorylation
Myc Targets V1
Ribosome
Coronavirus disease
Diabetic cardiomyopathy
Thermogenesis
Huntington disease
Natural Killer T (NKT) cell:Fetal Kidney
Morula Cell (Blastomere):Embryo
Purkinje cell:Brain
Epidermal Transit Amplifying cell:Foreskin
% Genes in set
CellMarker_Augmented_2021
KEGG_2021_Human
MSigDB_Hallmark_2020
Gene_set

F

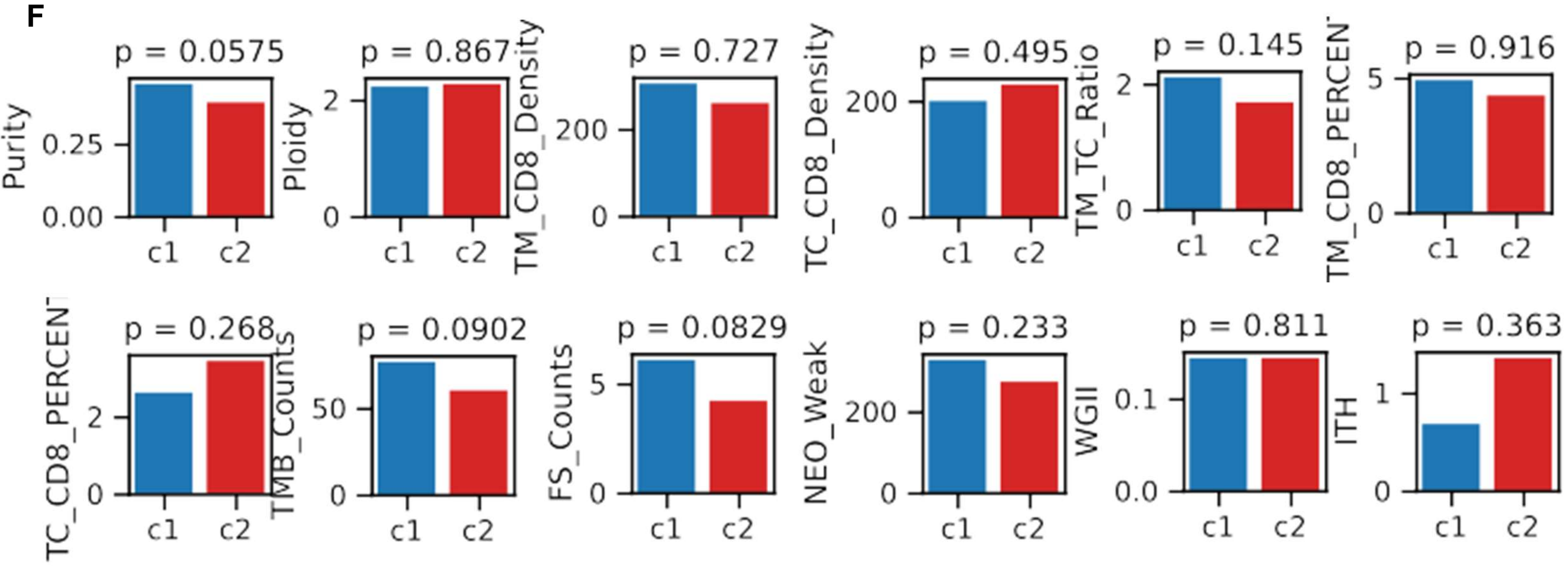


**Supplementary Figure S7. Comparison between two clusters generated from their negative correlation based gene connectivity of pN.** (A). Volcano plot of significant genes related to overall survival and progression free survival. P value below 0.01 was taken as significant. (B). Hierarchy clustering of samples based on selected genes. Genes were used if they were significantly related to both OS and PFS. Two clusters (c1 and c2) were preferred. Out of 48 selected genes, 5 of them ('MIR31HG', 'BARX2', 'LINC01322', 'WFDC10A', 'MYO9B') in terms of their expression values were associated with survival data. (C). Survival analysis between cluster c1 (blue) and c2 (pink). P values were from the log rank test. (D). Distribution of chromosomal and gene mutation between clusters. Fisher test was conducted and p value less than 0.05 was considered as significant. (E). Overall representation analysis. Genes were selected as the union of significant genes related to OS and PFS. (F). Comparison of clinical features between two clusters.

Six genes ('ACIN1', 'INF2', 'MIR31HG', 'MYO9B', 'PLCB3', 'WFDC10A') were found in both positive and negative associations.

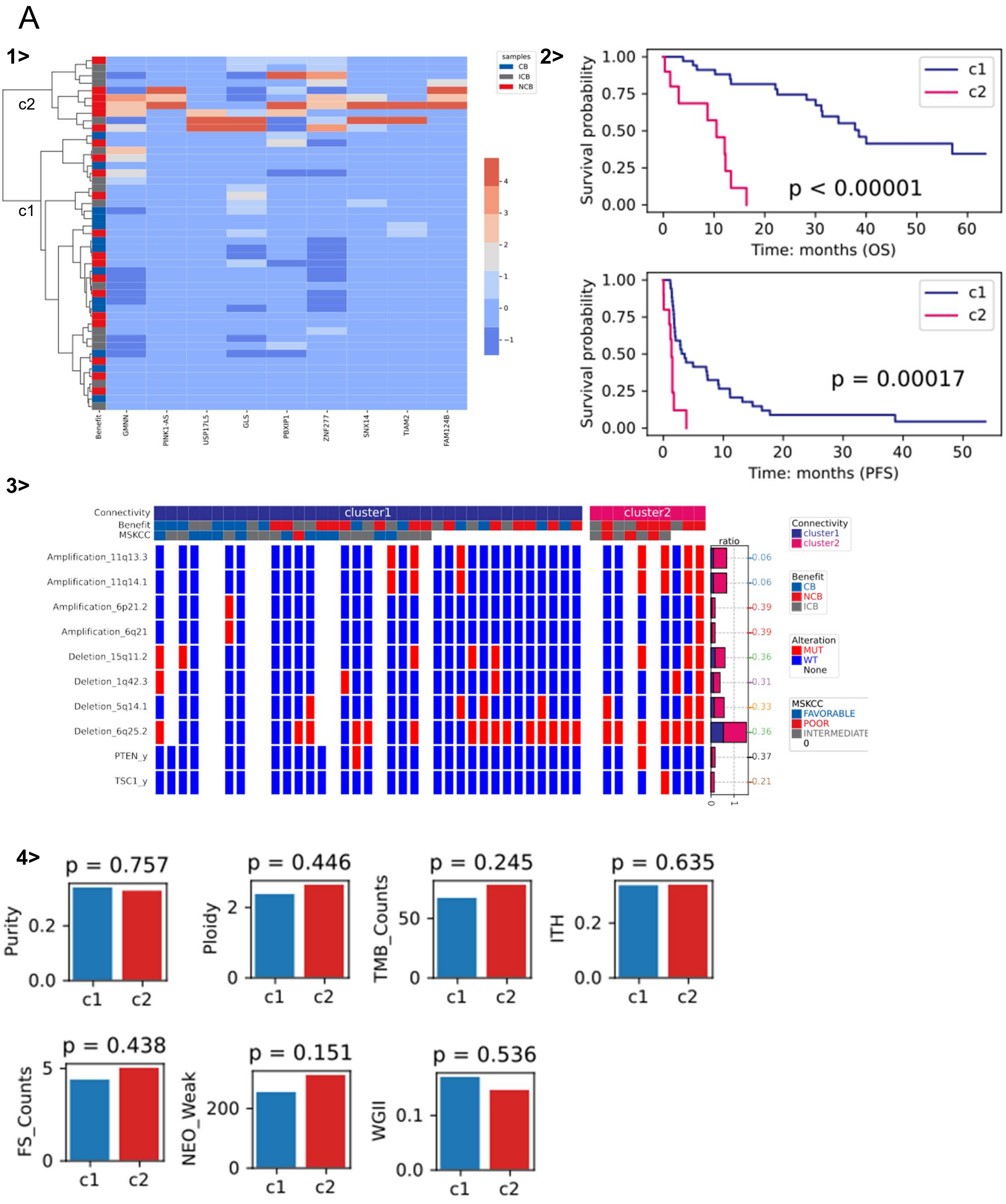

A
1>
c2
c1
samples
CB
ICB
NCB
Benefit
GMNN
PINK1-AS
USP17L5
GLS
PBXIP1
ZNF277
SNX14
TIAM2
FAM124B
2>
Survival probability
c1
c2
p < 0.00001
Time: months (OS)
p = 0.00017
Time: months (PFS)
3>
Connectivity
Benefit
MSKCC
cluster1
cluster2
Amplification_11q13.3
Amplification_11q14.1
Amplification_6p21.2
Amplification_6q21
Deletion_15q11.2
Deletion_1q42.3
Deletion_5q14.1
Deletion_6q25.2
PTEN_y
TSC1_y
ratio
0.06
0.06
0.39
0.39
0.36
0.31
0.33
0.36
0.37
0.21
Connectivity
cluster1
cluster2
Benefit
CB
NCB
ICB
Alteration
MUT
WT
None
MSKCC
FAVORABLE
POOR
INTERMEDIATE
4>
p = 0.757
Purity
p = 0.446
Ploidy
p = 0.245
TMB_Counts
p = 0.635
ITH
p = 0.438
FS_Counts
p = 0.151
NEO_Weak
p = 0.536
WGII
c1
c2

B

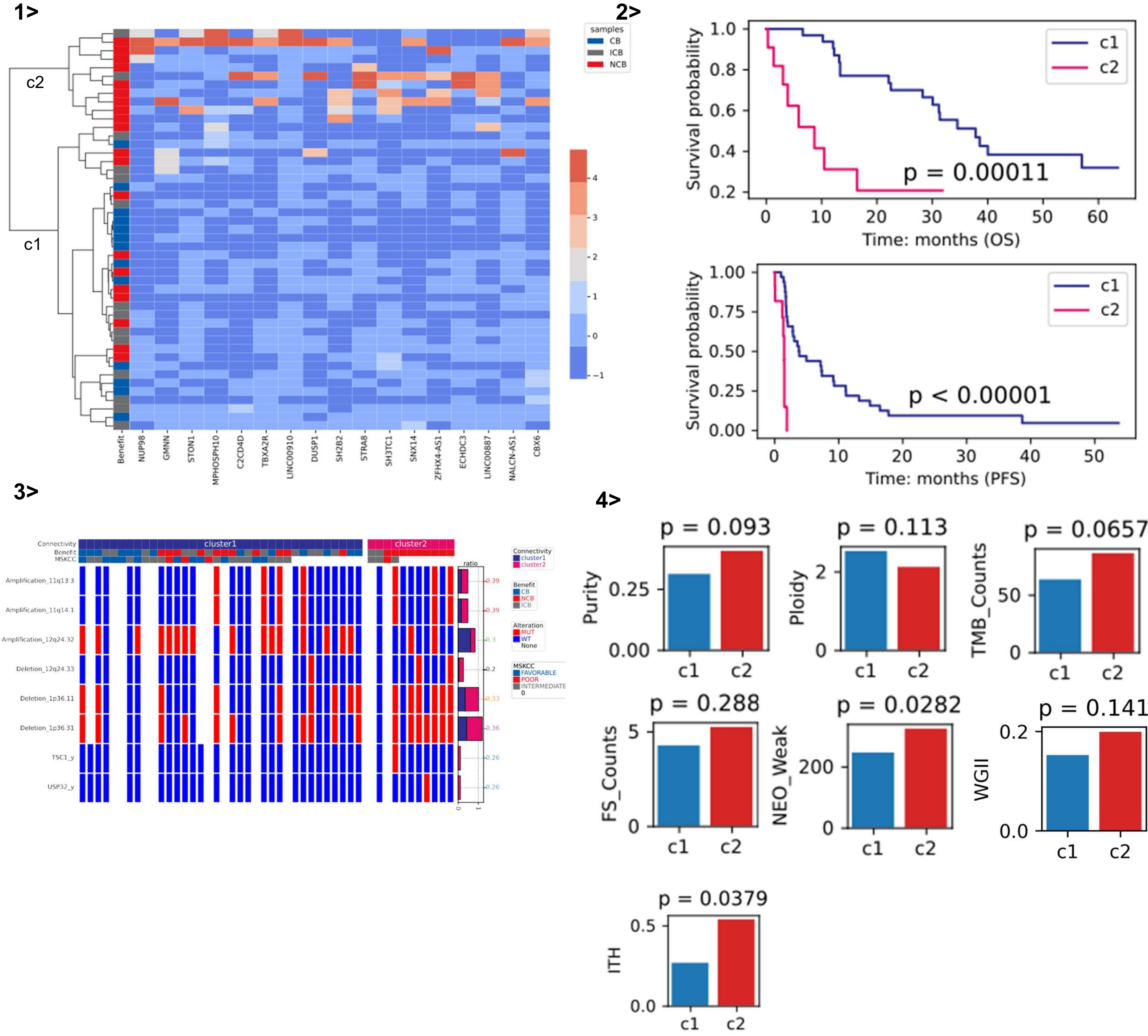


**Supplementary Figure S8. Comparison between two clusters generated from their positive correlation (A) and negative correlation (B) based gene connectivity of mN.** 1>. Hierarchy clustering of samples based on selected genes. Genes were selected if they were significantly related to both OS and PFS (9, and 17 genes were obtained from positive and negative based gene connectivity). Two clusters (c1 and c2) were preferred. Out of these genes, expression values of 1, and 6 of them ( 'USP17L5'; 'LINC00887', 'ZFHX4-AS1', 'TBXA2R', 'ECHDC3', 'NALCN-AS1', 'LINC00910') were associated with survival data. 2>. Survival analysis between cluster c1 (blue) and c2 (pink). P values were from the log rank test. 3>. Distribution of chromosomal and gene mutation between clusters. Fisher test was conducted and p value less than 0.05 was considered as significant. 4>. Comparison of clinical features between the two clusters.

Noted that we did not enrich any pathway for over representation analysis and two genes ('GMNN', 'SNX14') were found in both positive and negative associations.

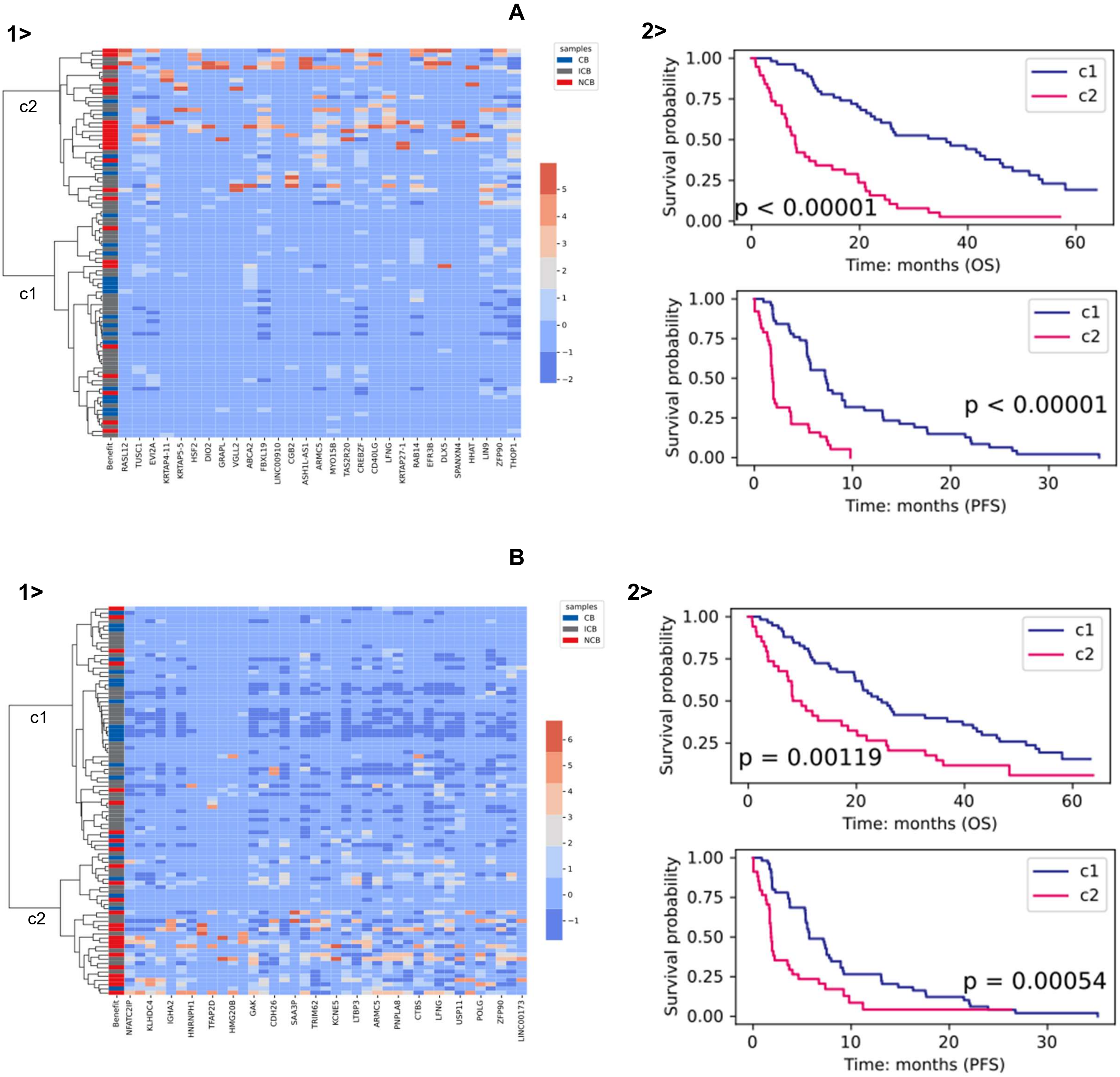


**Supplementary Figure S9. Comparison between two clusters generated from their positive correlation (A) and negative correlation (B) based gene connectivity of pE.** 1>. Hierarchy clustering of samples based on selected genes. Genes were selected if they were significantly related to both OS and PFS (29, and 39 genes were obtained from positive and negative based gene connectivity). Two clusters (c1 and c2) were preferred. 2>. Survival analysis between cluster c1 (blue) and c2 (pink). P values were from the log rank test.

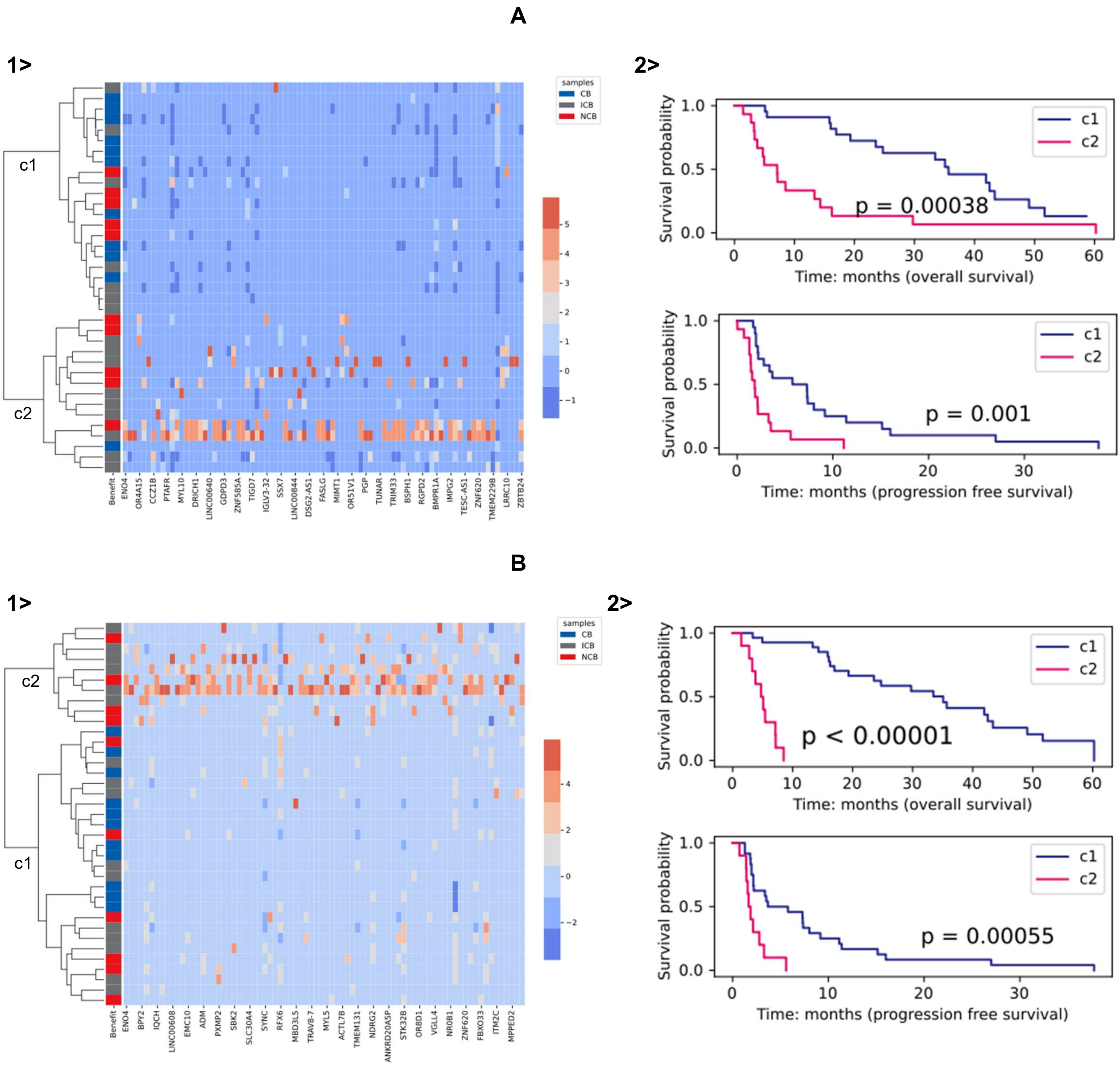


**Supplementary Figure S10. Comparison between two clusters generated from their positive correlation (A) and negative correlation (B) based gene connectivity of mE.** 1>. Hierarchy clustering of samples based on selected genes. Genes were selected if they were significantly related to either OS or PFS (78, and 85 genes were obtained from positive and negative based gene connectivity). No common gene was found between OS significant genes and PFS significant genes. Two clusters (c1 and c2) were preferred. 2>. Survival analysis between cluster c1 (blue) and c2 (pink). P values were from the log rank test.

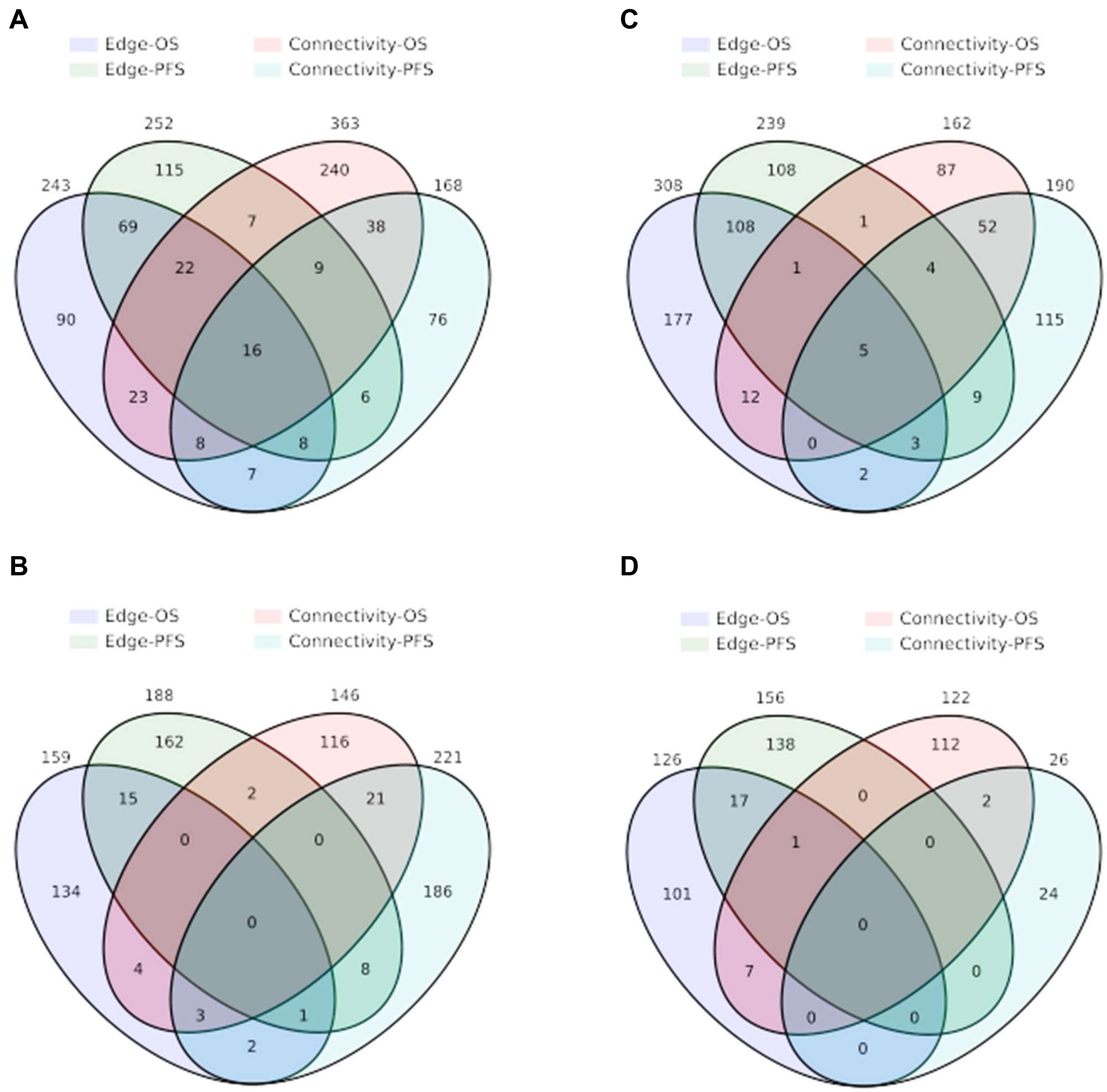


**Supplementary Figure S11. Venn plots for genes from gene connectivity and edges.** (A). pN. (B). mN. (C). pE. (D). mE.

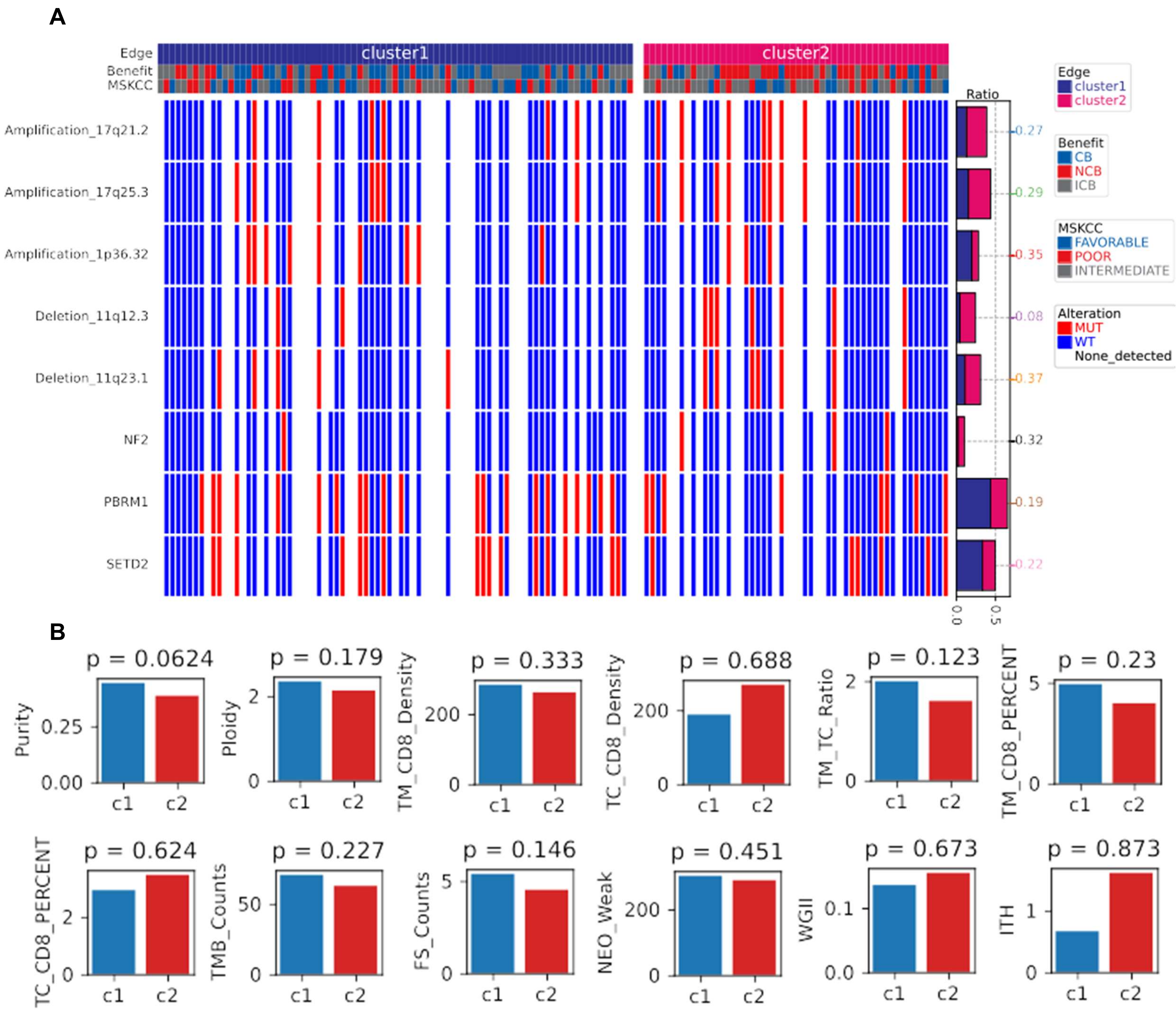


**Supplementary Figure S12. Comparison of genomic mutation and clinical features between two clusters generated from selected edges of pN ssGCNs. A. Mutation plot. B. Bar plots for clinical features.** Wilcoxon rank sum tests were conducted and p values below 0.05 were taken as significant. Cluster c1 has high PCC on average and were associated with higher survival probability.

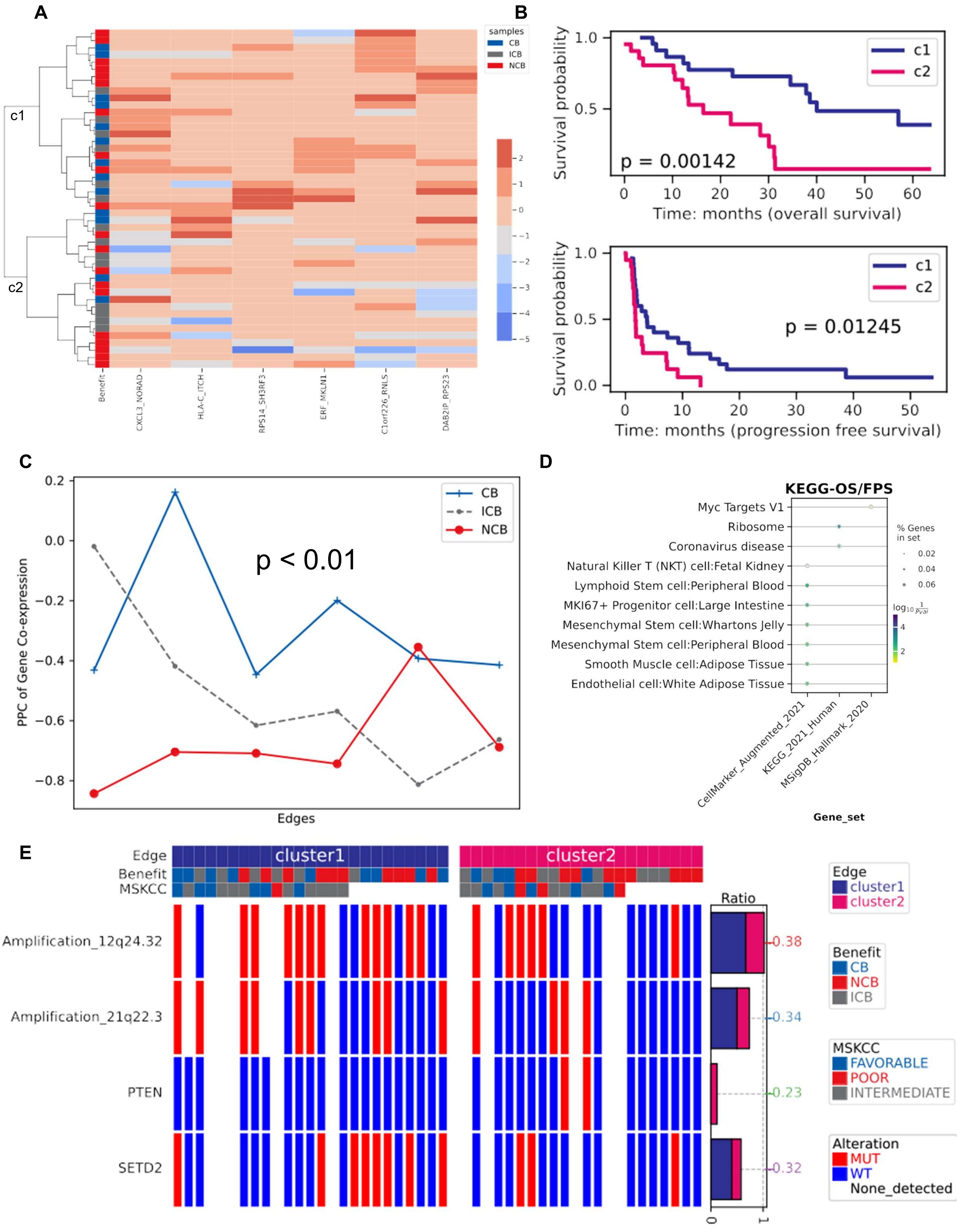
A
B
C
D
E
samples
CB
ICB
NCB
c1
c2
Benefit
CXCL3_NORAD
HLA-C_ITCH
RPS14_SH3RF3
ERF_MKLN1
C1orf226_RNLS
DAB2IP_RPS23
Survival probability
p = 0.00142
Time: months (overall survival)
p = 0.01245
Time: months (progression free survival)
PPC of Gene Co-expression
p < 0.01
Edges
KEGG-OS/FPS
Myc Targets V1
Ribosome
Coronavirus disease
Natural Killer T (NKT) cell:Fetal Kidney
Lymphoid Stem cell:Peripheral Blood
MKI67+ Progenitor cell:Large Intestine
Mesenchymal Stem cell:Whartons Jelly
Mesenchymal Stem cell:Peripheral Blood
Smooth Muscle cell:Adipose Tissue
Endothelial cell:White Adipose Tissue
% Genes in set
CellMarker_Augmented_2021
KEGG_2021_Human
MSigDB_Hallmark_2020
Gene_set
Edge
Benefit
MSKCC
cluster1
cluster2
Amplification_12q24.32
Amplification_21q22.3
PTEN
SETD2
Ratio
0.38
0.34
0.23
0.32
FAVORABLE
POOR
INTERMEDIATE
Alteration
MUT
WT
None_detected

F

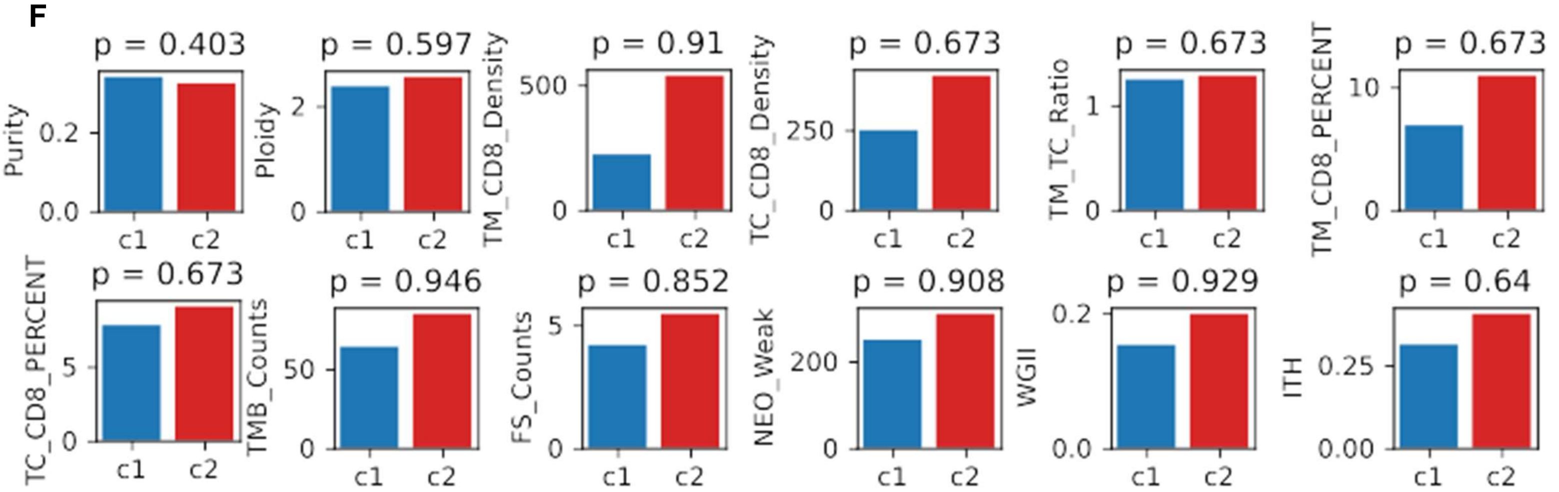


**Supplementary Figure S13. Edges classified samples of mN.** (A). Hierarchy clustering of samples using 6 edges significantly associated with both OS and PFS. (B). Survival analysis between two clusters c1 (blue) and c2 (pink). P values were from the log rank test. (C). Distributions of pearson correlation coefficients (PCC) for mN samples of CB (blue), ICB (grey) and NCB (red). Wilcoxon rank sum tests were conducted between CB and NCB patients. (D). Overall representation analysis. Genes were selected as the union of genes from significant edges significantly related to OS and PFS. (E). Mutation plot. Fisher test was conducted. (F). Bar plots for clinical features. Wilcoxon rank sum tests were conducted. p values below 0.05 were taken as significant.

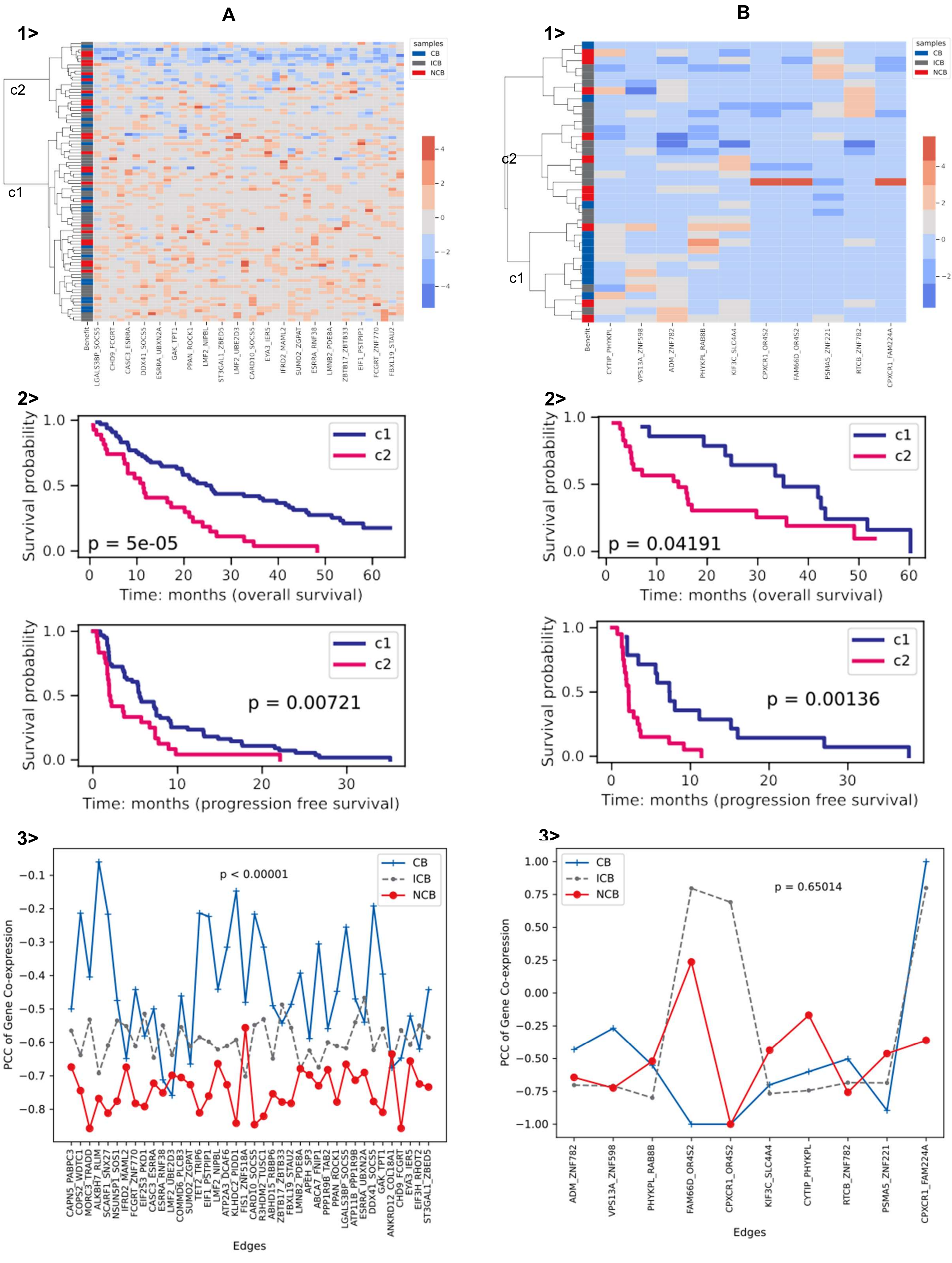

A
B
1>
c1
c2
samples
CB
ICB
NCB
Benefit
LGALS3BP_SOCS5
CHD9_FCGRT
CASC3_ESRRA
DDX41_SOCS5
ESRRA_UBXN2A
GAK_TPT1
PPAN_ROCK1
LMF2_NIPBL
ST3GAL1_ZBED5
LMF2_UBE2D3
CARD10_SOCS5
EYA3_IER5
IFRD2_MAML2
SUMO2_ZGPAT
ESRRA_RNF38
LMNB2_PDE8A
ZBTB17_ZBTB33
EIF1_PSTPIP1
FCGRT_ZNF770
FBXL19_STAU2
CYTIP_PHYKPL
VPS13A_ZNF598
ADM_ZNF782
PHYKPL_RAB8B
KIF3C_SLC4A4
CPXCR1_OR4S2
FAM66D_OR4S2
PSMA5_ZNF221
RTCB_ZNF782
CPXCR1_FAM224A
2>
Survival probability
Time: months (overall survival)
Time: months (progression free survival)
p = 5e-05
p = 0.00721
p = 0.04191
p = 0.00136
3>
PCC of Gene Co-expression
Edges
p < 0.00001
p = 0.65014
CAPN5_PABPC3
COPS2_WDTC1
MORC3_TRADD
ALKBH7_RLIM
SCARF1_SNX27
NSUN5P1_SOS1
IFRD2_MAML2
FCGRT_ZNF770
EIF2S3_PKD1
CASC3_ESRRA
ESRRA_RNF38
LMF2_UBE2D3
COMMD6_PLCB3
SUMO2_ZGPAT
TET2_PSTPIP1
EIF1_PSTPIP1
LMF2_NIPBL
ATP2A3_DCAF6
KLHDC2_PIDD1
FIS1_ZNF518A
CARD10_SOCS5
R3HDM2_TUSC1
ABHD15_RBBP6
ZBTB17_ZBTB33
FBXL19_STAU2
LMNB2_PDE8A
APEH_SP3
ABCA7_FNIP1
PPP1R9B_TAB2
PPAN_ROCK1
LGALS3BP_SOCS5
ATP11B_PPP1R9B
ESRRA_UBXN2A
DDX41_SOCS5
GAK_TPT1
ANKRD12_COL18A1
CHD9_FCGRT
EYA3_IER5
EIF3H_RHOT2
ST3GAL1_ZBED5

**Supplementary Figure S14. Edges classified samples of pE (A) and mE (B).** 1>. Hierarchy clustering of samples using 40/ 10 edges significantly associated with both OS and PFS. 2>. Survival analysis between two clusters c1 (blue) and c2 (pink). P values were from the log rank test. 3>. Distributions of pearson correlation coefficients (PCC) for mN samples of CB (blue), ICB (grey) and NCB (red). Wilcoxon rank sum tests were conducted between CB and NCB patients.

**A**

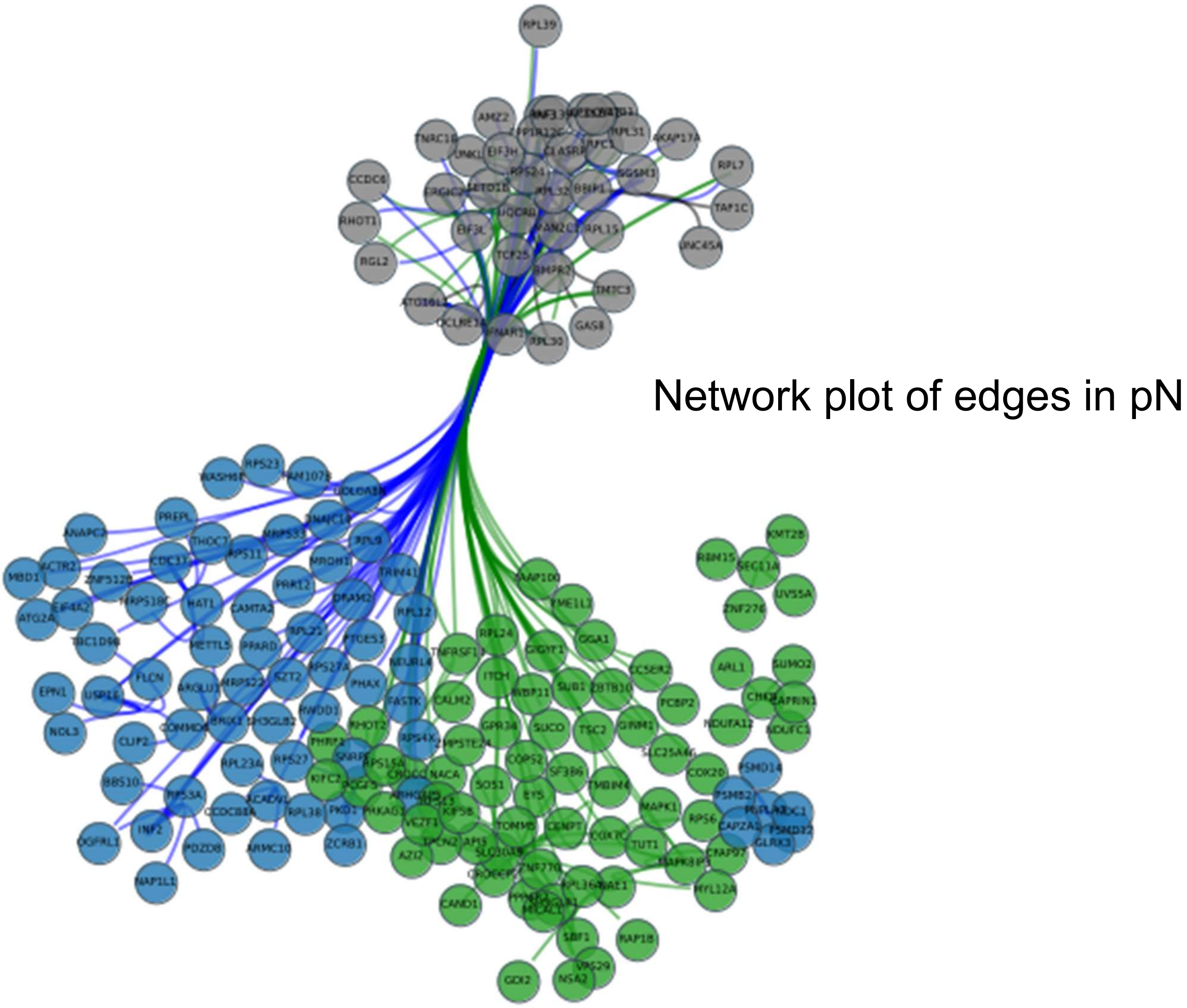


**B**

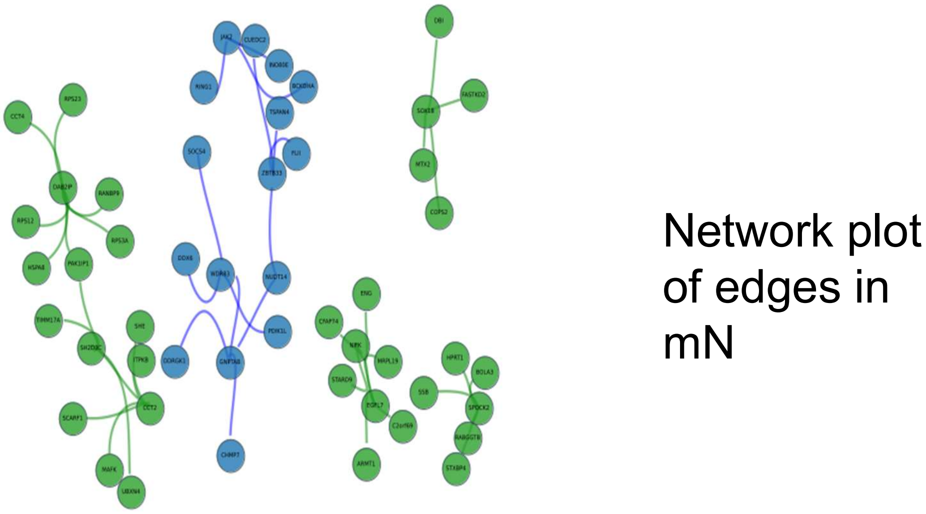

C

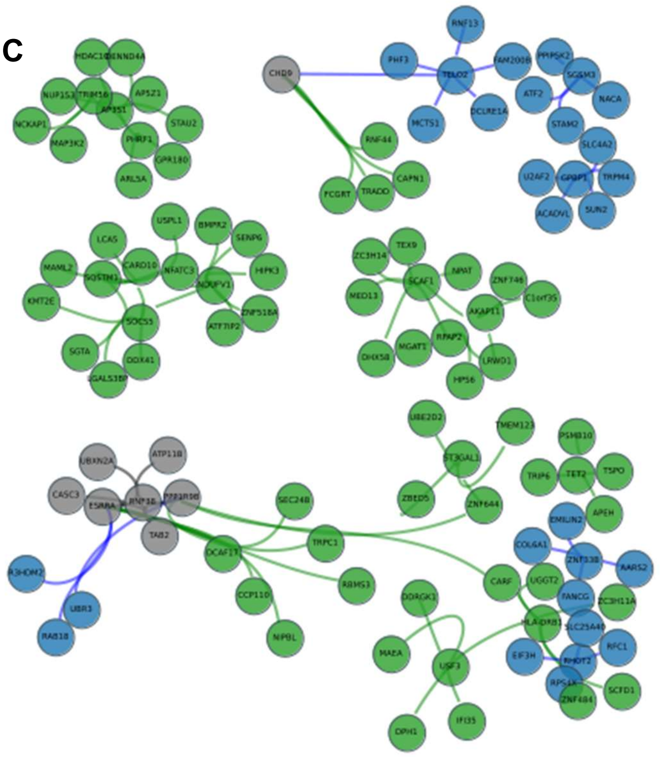

Network plot of edges in pE

D

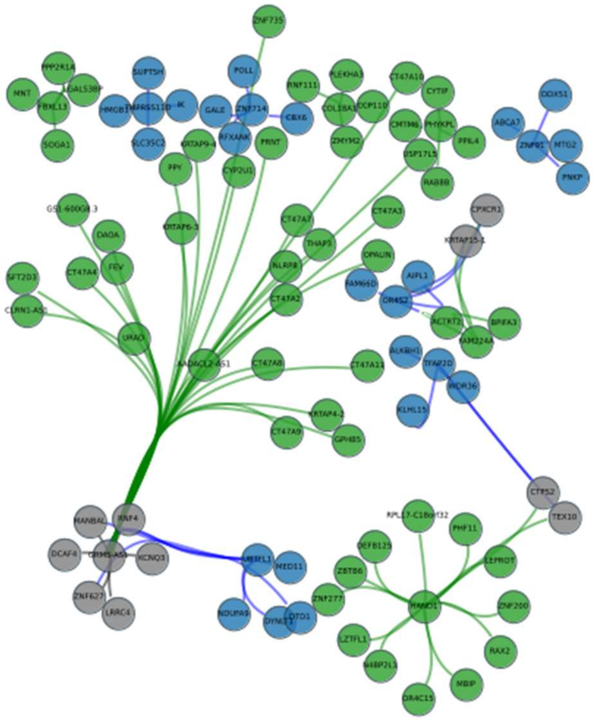

Network plot of edges in mE

**Supplementary Figure S15. Network plots of selected edges significantly associated with either OS or PFS.** Genes, showing more than 3 times across edges, were displayed here. Black color indicated genes from both OS and PFS, and green / blue color refers that genes related with only OS or PFS. (A). pN. (B). mN. (C). pE. (D). mE. Noted that for mN and Me, significant edges were selected by p value 0f 0.05.

OS significant pathways (left) and PFS significant pathways (right)

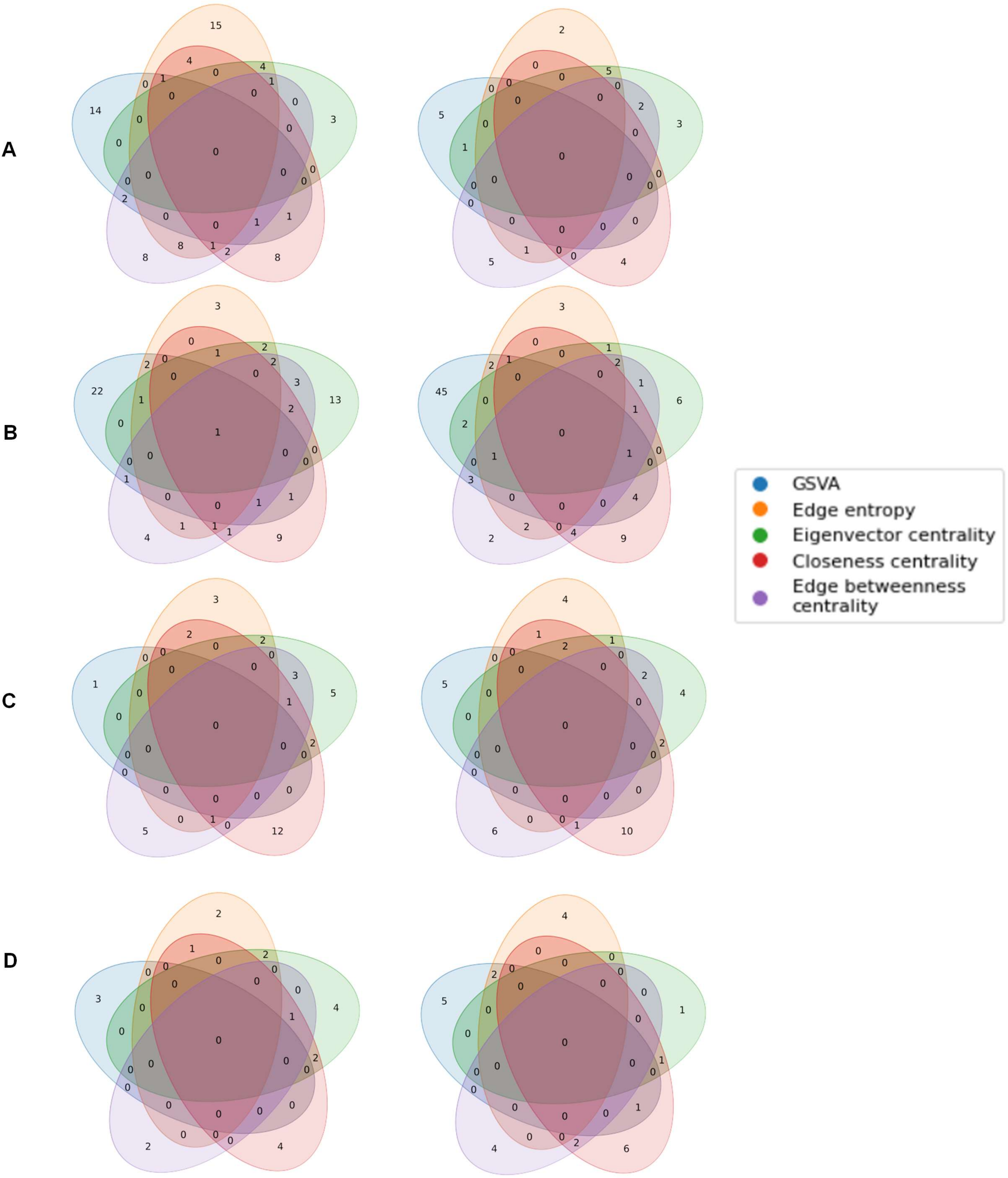


**Supplementary Figure S16. Venn plot of signficant pathways in the subcohorts: pN (A), mN (B), pE (C), mE (D).** The significant pathways were selected if their p values were less than 0.05 from the cox regression models. These significant pathways were barely overlapped with the others and it may be explained that they were calculated based on gene expression, network complexity and the influence of genes or edges.

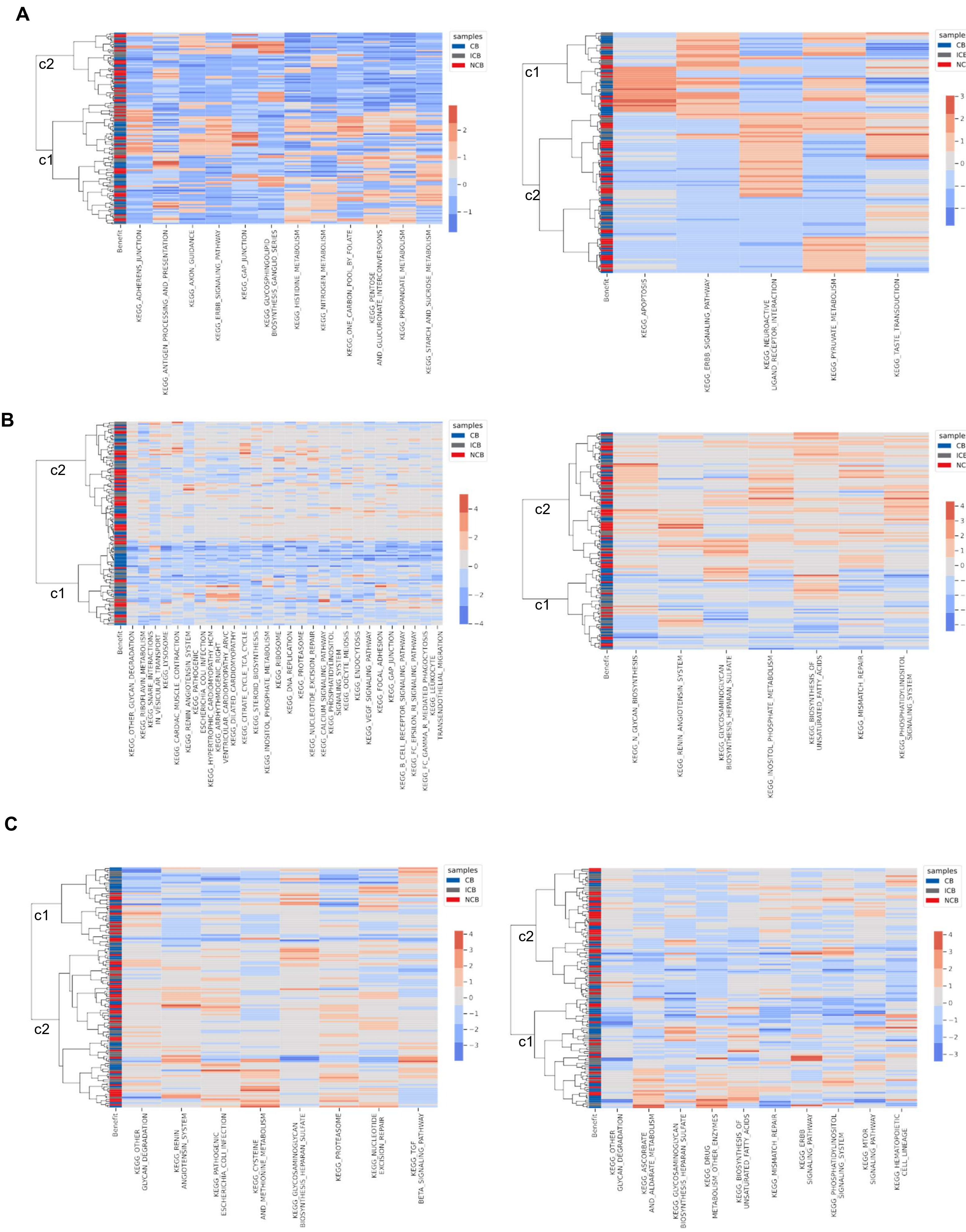
A
B
C
c1
c2
samples
CB
ICB
NCB
Benefit
KEGG_ADHERENS_JUNCTION
KEGG_ANTIGEN_PROCESSING_AND_PRESENTATION
KEGG_AXON_GUIDANCE
KEGG_ERBB_SIGNALING_PATHWAY
KEGG_GAP_JUNCTION
KEGG_GLYCOSPHINGOLIPID BIOSYNTHESIS_GANGLIO_SERIES
KEGG_HISTIDINE_METABOLISM
KEGG_NITROGEN_METABOLISM
KEGG_ONE_CARBON_POOL_BY_FOLATE
KEGG_PENTOSE AND_GLUCURONATE_INTERCONVERSIONS
KEGG_PROPANOATE_METABOLISM
KEGG_STARCH_AND_SUCROSE_METABOLISM
KEGG_APOPTOSIS
KEGG_NEUROACTIVE LIGAND_RECEPTOR_INTERACTION
KEGG_PYRUVATE_METABOLISM
KEGG_TASTE_TRANSDUCTION
KEGG_N_GLYCAN_BIOSYNTHESIS
KEGG_RENIN_ANGIOTENSIN_SYSTEM
KEGG_GLYCOSAMINOGLYCAN BIOSYNTHESIS_HEPARAN_SULFATE
KEGG_INOSITOL_PHOSPHATE_METABOLISM
KEGG_BIOSYNTHESIS_OF UNSATURATED_FATTY_ACIDS
KEGG_MISMATCH_REPAIR
KEGG_PHOSPHATIDYLINOSITOL SIGNALING_SYSTEM
KEGG_OTHER GLYCAN_DEGRADATION
KEGG_PATHOGENIC ESCHERICHIA_COLI_INFECTION
KEGG_CYSTEINE AND_METHIONINE_METABOLISM
KEGG_PROTEASOME
KEGG_NUCLEOTIDE EXCISION_REPAIR
KEGG_TGF BETA_SIGNALING_PATHWAY
KEGG_ASCORBATE AND_ALDARATE_METABOLISM
KEGG_DRUG METABOLISM_OTHER_ENZYMES
KEGG_ERBB SIGNALING_PATHWAY
KEGG_MTOR SIGNALING_PATHWAY
KEGG_HEMATOPOIETIC CELL_LINEAGE

D

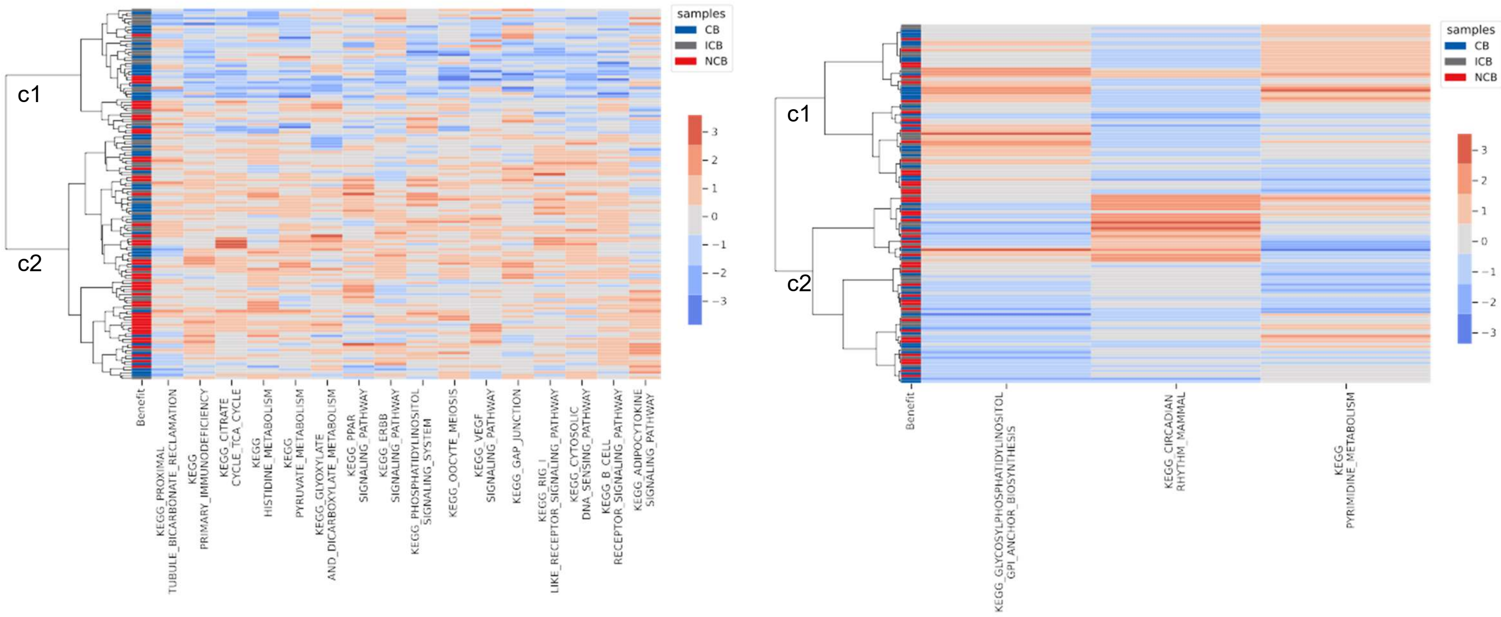


E

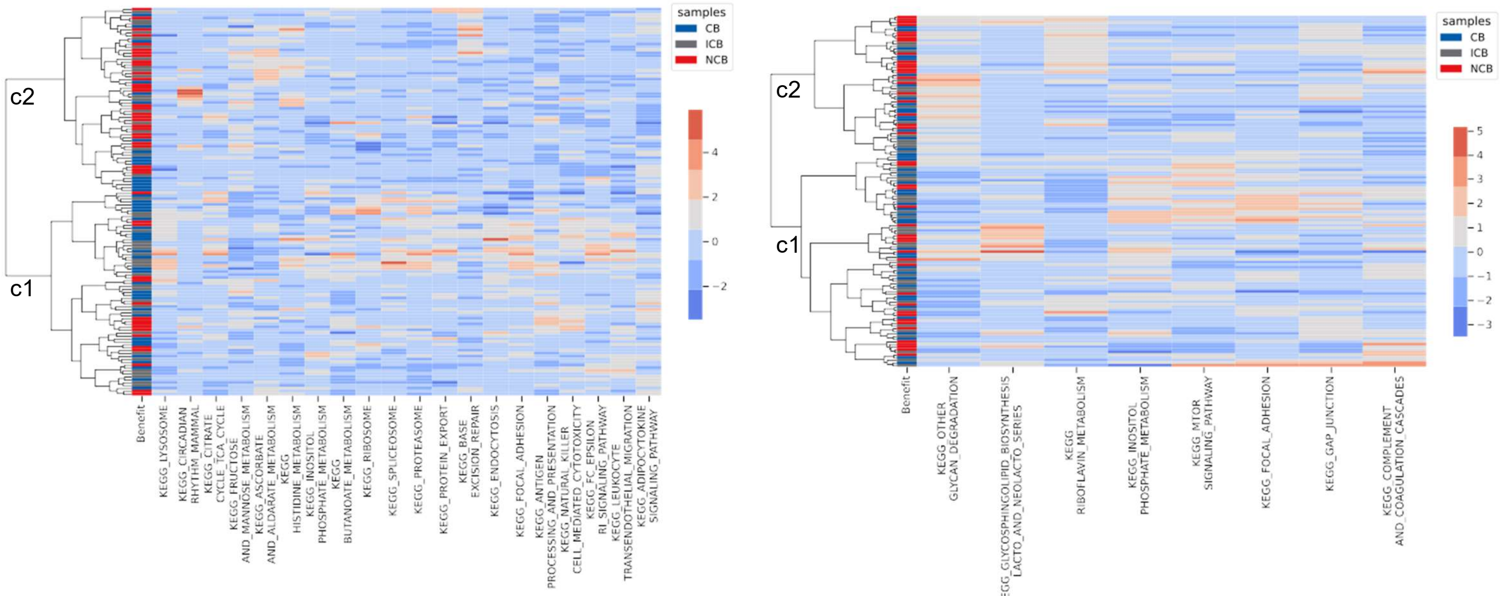


**Supplementary Figure S17. Clustermap of pN samples using pathway scores.** (A). 12 (OS)/ 5 (PFS) significant pathways based on GSVA. (B). 28 (OS)/ 7 (PFS) significant pathways based on entropy. (C). 8 (OS)/ 10 (PFS) significant pathways based on gene eigenvector centrality scores. (D). 16 (OS)/ 3 (PFS) significant pathways based on gene closeness centrality scores. (E).20 (OS)/ 8 (PFS) significant pathways based on edge betweenness centrality score. Note that irrelevant pathways were filtered out for the clustering step in pathway analysis.

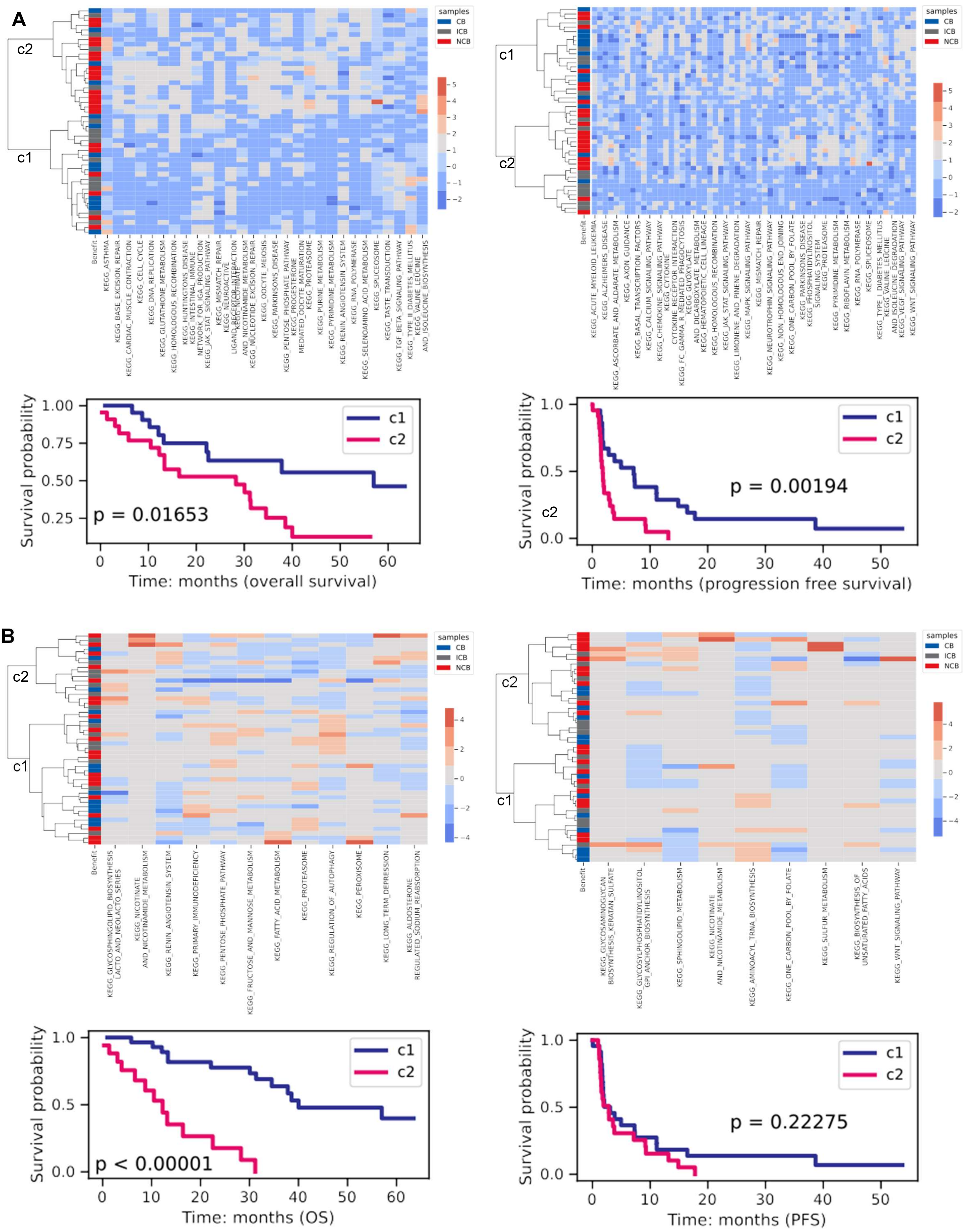
A
B
samples
CB
ICB
NCB
Benefit
c1
c2
Survival probability
p = 0.01653
Time: months (overall survival)
p = 0.00194
Time: months (progression free survival)
p < 0.00001
Time: months (OS)
p = 0.22275
Time: months (PFS)

C

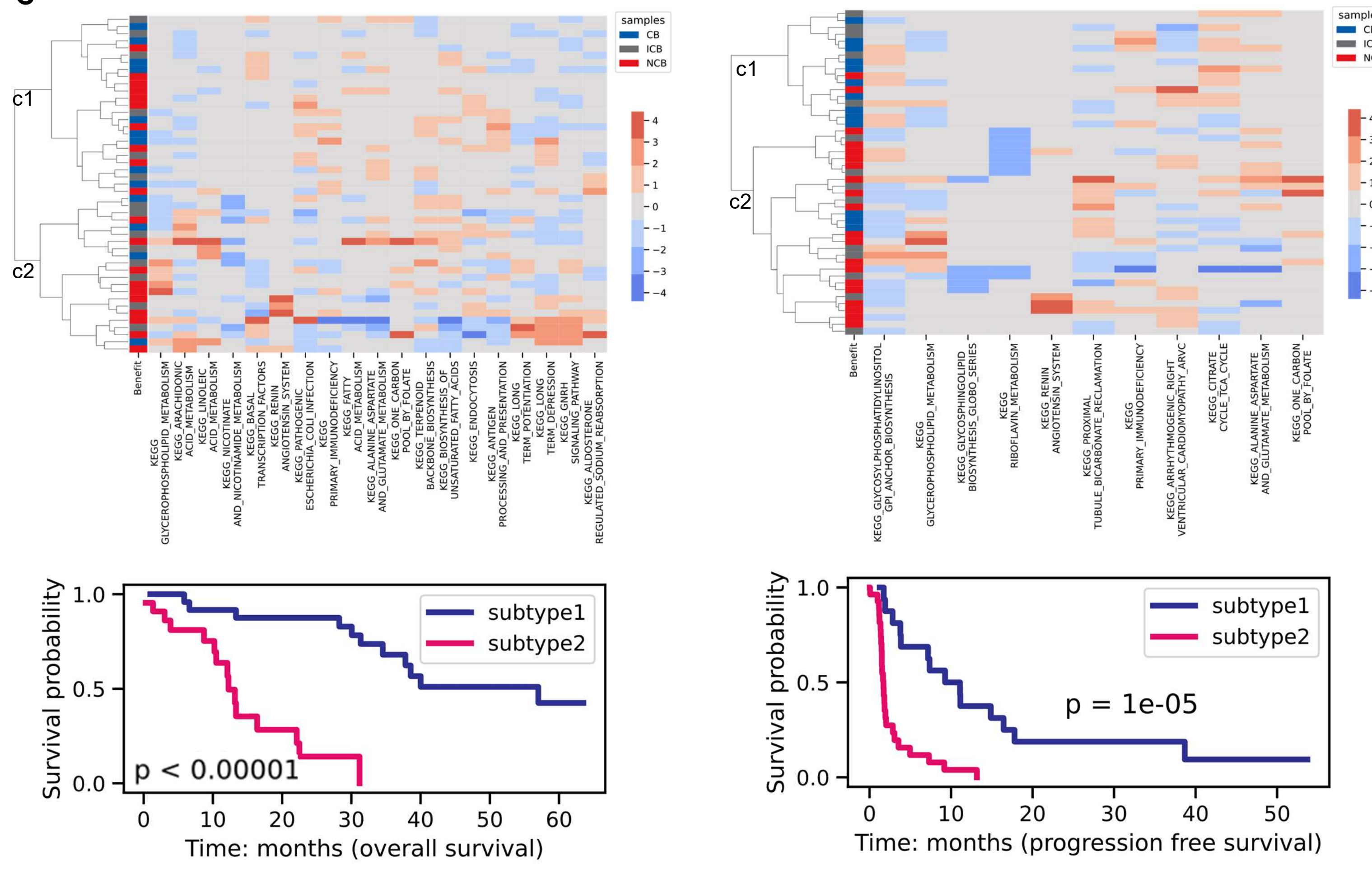

samples
CB
ICB
NCB
c1
c2
Benefit
subtype1
subtype2
p < 0.00001
Survival probability
Time: months (overall survival)
p = 1e-05
Time: months (progression free survival)


D

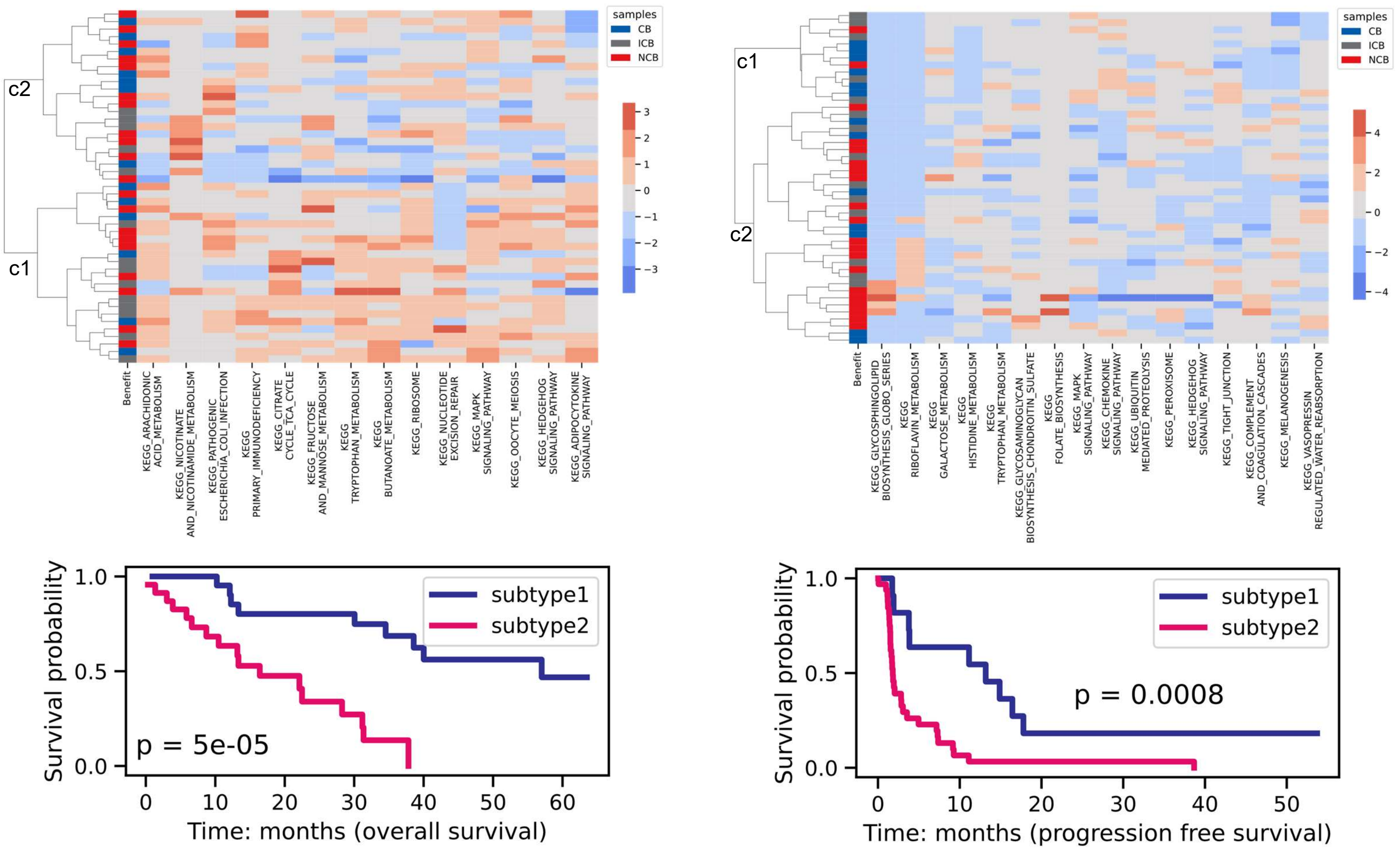

samples
CB
ICB
NCB
c1
c2
Benefit
subtype1
subtype2
p = 5e-05
Survival probability
Time: months (overall survival)
p = 0.0008
Time: months (progression free survival)

**E**

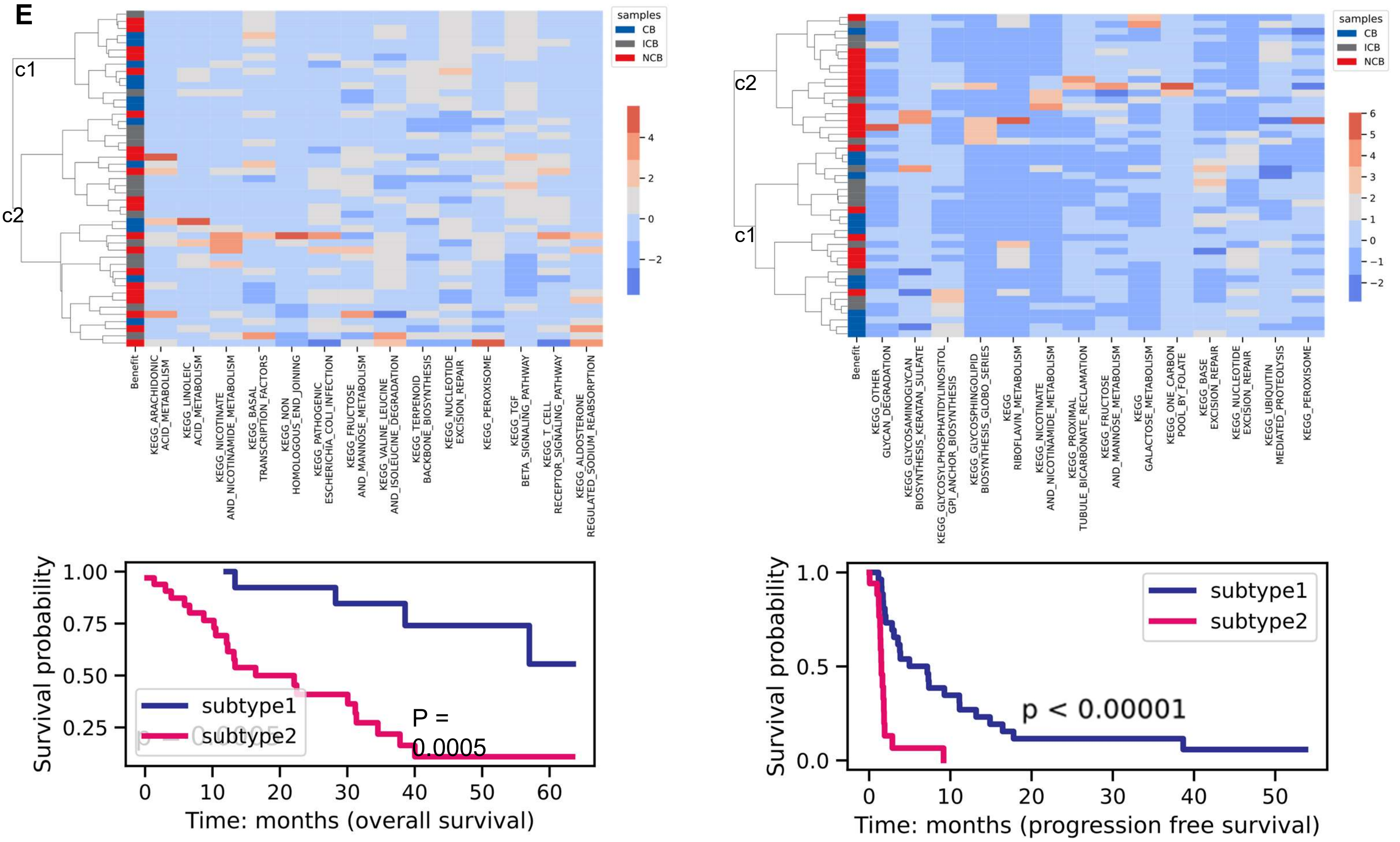


**Supplementary Figure S18. Clustermap and survival analysis of mN samples using pathway scores.** (A). 29 (OS)/ 59 (PFS) significant pathways based on GSVA. (B). 12 (OS)/ 10 (PFS) significant pathways based on entropy. (C). 19 (OS)/ 11 (PFS) significant pathways based on gene eigenvector centrality scores. (D). 14 (OS)/ 16 (PFS) significant pathways based on gene closeness centrality scores. (E). 14 (OS)/ 14 (PFS) significant pathways based on edge betweenness centrality score.

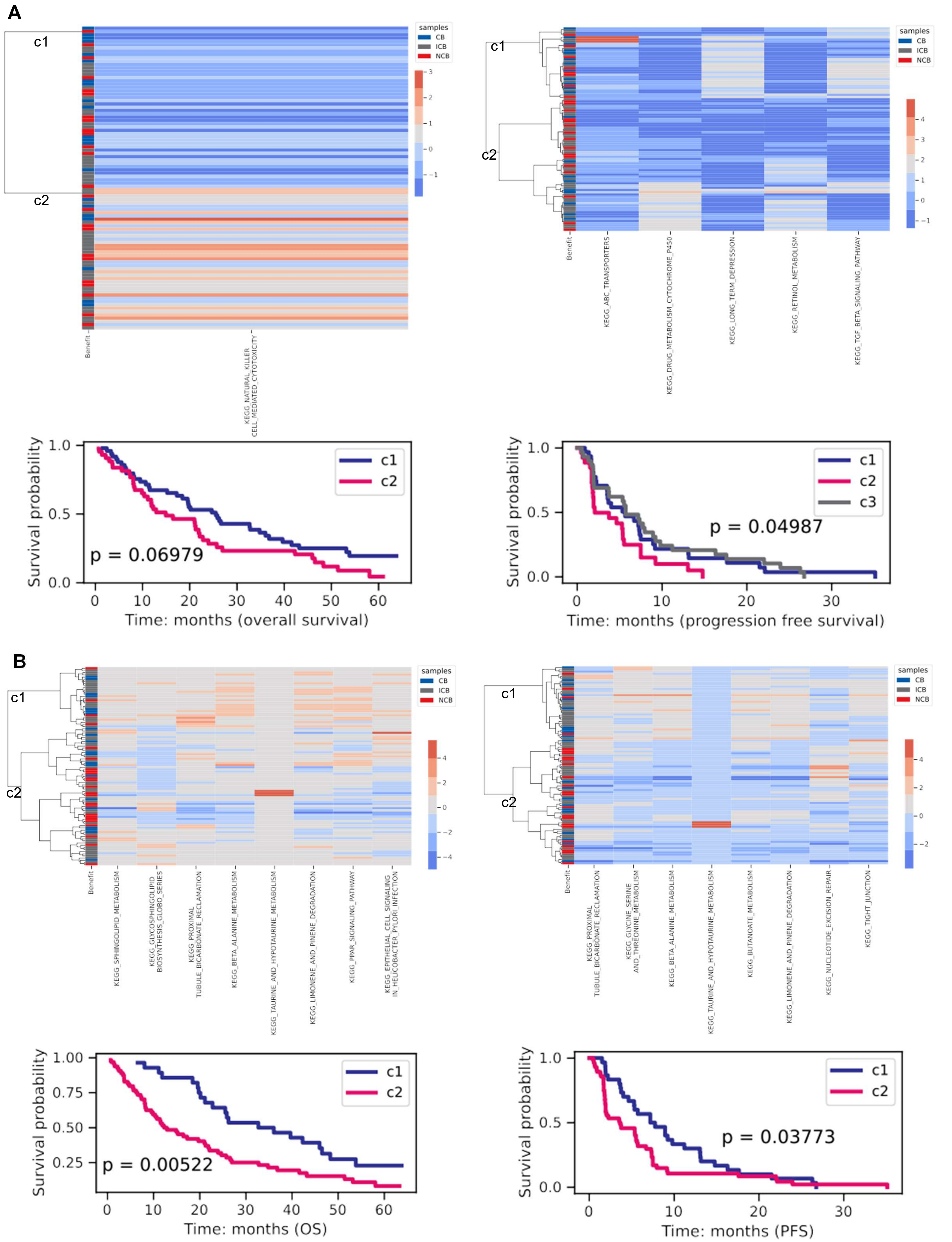
A
c1
c2
samples
CB
ICB
NCB
Benefit
KEGG_NATURAL_KILLER_CELL_MEDIATED_CYTOTOXICITY
KEGG_ABC_TRANSPORTERS
KEGG_DRUG_METABOLISM_CYTOCHROME_P450
KEGG_LONG_TERM_DEPRESSION
KEGG_RETINOL_METABOLISM
KEGG_TGF_BETA_SIGNALING_PATHWAY
Survival probability
p = 0.06979
Time: months (overall survival)
c3
p = 0.04987
Time: months (progression free survival)
B
KEGG_SPHINGOLIPID_METABOLISM
KEGG_GLYCOSPHINGOLIPID_BIOSYNTHESIS_GLOBO_SERIES
KEGG_PROXIMAL_TUBULE_BICARBONATE_RECLAMATION
KEGG_BETA_ALANINE_METABOLISM
KEGG_TAURINE_AND_HYPOTAURINE_METABOLISM
KEGG_LIMONENE_AND_PINENE_DEGRADATION
KEGG_PPAR_SIGNALING_PATHWAY
KEGG_EPITHELIAL_CELL_SIGNALING_IN_HELICOBACTER_PYLORI_INFECTION
KEGG_GLYCINE_SERINE_AND_THREONINE_METABOLISM
KEGG_BUTANOATE_METABOLISM
KEGG_NUCLEOTIDE_EXCISION_REPAIR
KEGG_TIGHT_JUNCTION
p = 0.00522
Time: months (OS)
p = 0.03773
Time: months (PFS)

C

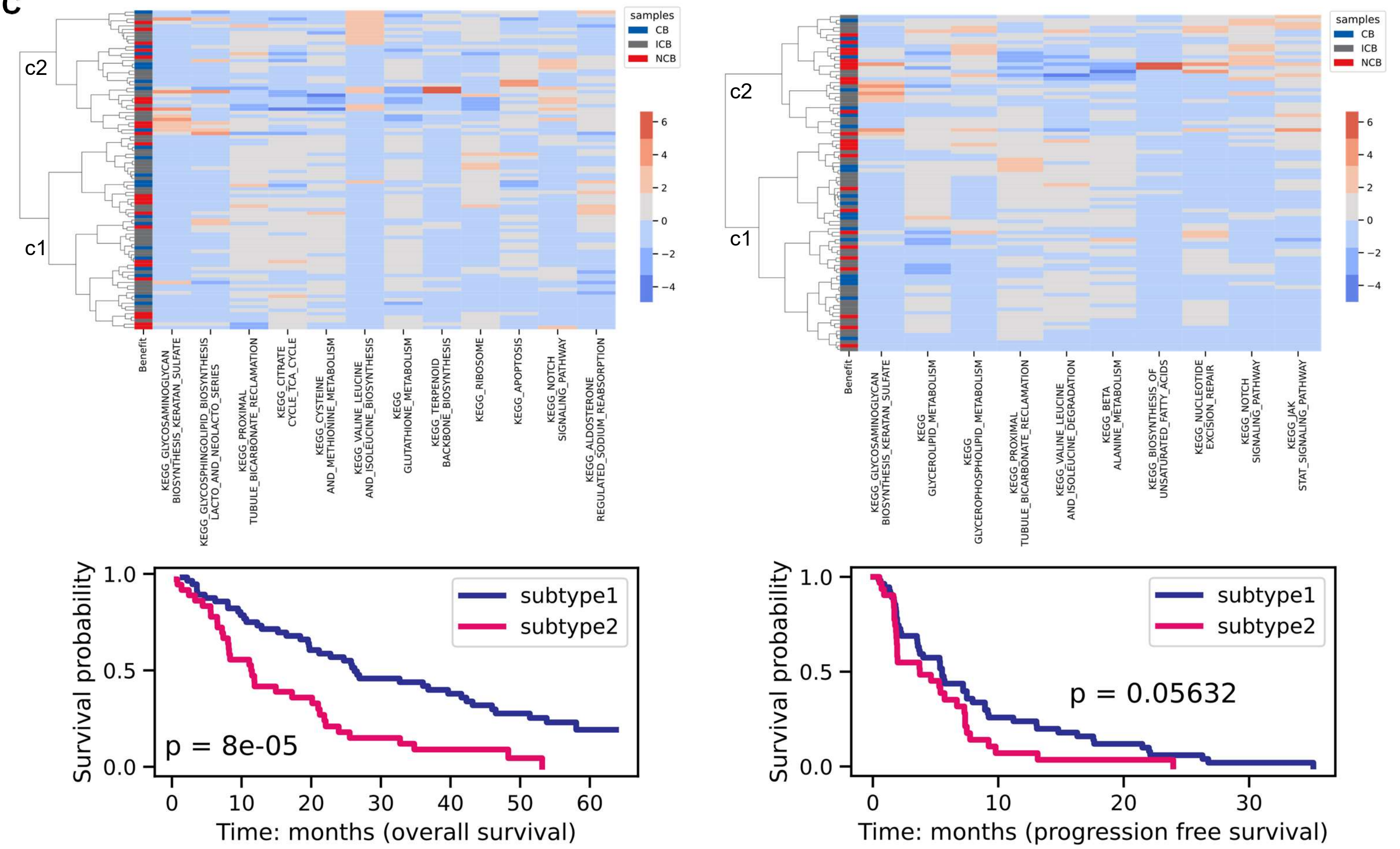
c2
c1
samples
CB
ICB
NCB
Benefit
KEGG_GLYCOSAMINOGLYCAN BIOSYNTHESIS_KERATAN_SULFATE
KEGG_GLYCOSPHINGOLIPID_BIOSYNTHESIS LACTO_AND_NEOLACTO_SERIES
KEGG_PROXIMAL TUBULE_BICARBONATE_RECLAMATION
KEGG_CITRATE CYCLE_TCA_CYCLE
KEGG_CYSTEINE AND_METHIONINE_METABOLISM
KEGG_VALINE_LEUCINE AND_ISOLEUCINE_BIOSYNTHESIS
KEGG GLUTATHIONE_METABOLISM
KEGG_TERPENOID BACKBONE_BIOSYNTHESIS
KEGG_RIBOSOME
KEGG_APOPTOSIS
KEGG_NOTCH SIGNALING_PATHWAY
KEGG_ALDOSTERONE REGULATED_SODIUM_REABSORPTION
KEGG_GLYCOSAMINOGLYCAN BIOSYNTHESIS_KERATAN_SULFATE
KEGG GLYCEROLIPID_METABOLISM
KEGG GLYCEROPHOSPHOLIPID_METABOLISM
KEGG_PROXIMAL TUBULE_BICARBONATE_RECLAMATION
KEGG_VALINE_LEUCINE AND_ISOLEUCINE_DEGRADATION
KEGG_BETA ALANINE_METABOLISM
KEGG_BIOSYNTHESIS_OF UNSATURATED_FATTY_ACIDS
KEGG_NUCLEOTIDE EXCISION_REPAIR
KEGG_NOTCH SIGNALING_PATHWAY
KEGG_JAK STAT_SIGNALING_PATHWAY
Survival probability
subtype1
subtype2
p = 8e-05
Time: months (overall survival)
p = 0.05632
Time: months (progression free survival)

D

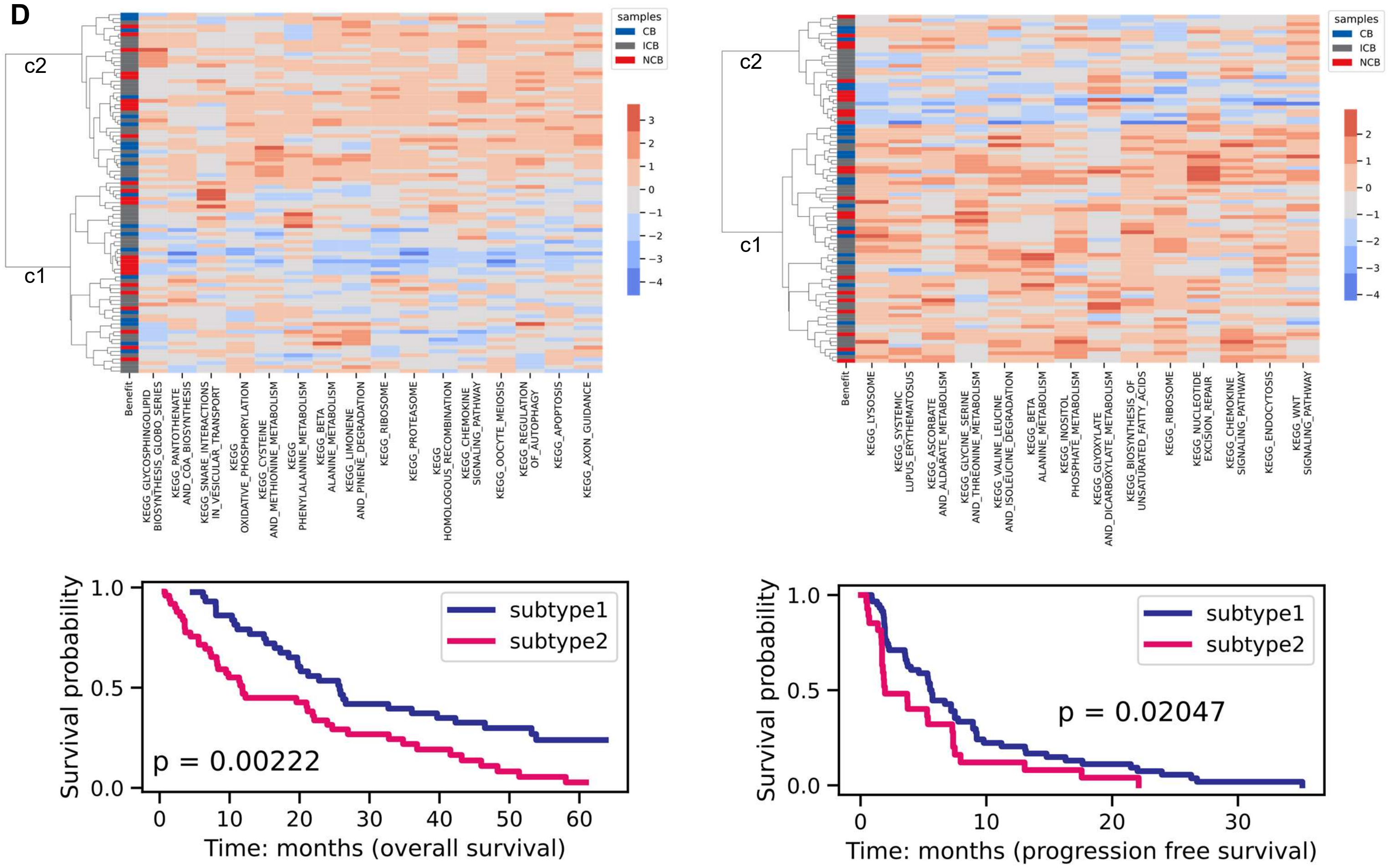
c2
c1
samples
CB
ICB
NCB
Benefit
KEGG_GLYCOSPHINGOLIPID BIOSYNTHESIS_GLOBO_SERIES
KEGG_PANTOTHENATE AND_COA_BIOSYNTHESIS
KEGG_SNARE_INTERACTIONS IN_VESICULAR_TRANSPORT
KEGG OXIDATIVE_PHOSPHORYLATION
KEGG_CYSTEINE AND_METHIONINE_METABOLISM
KEGG PHENYLALANINE_METABOLISM
KEGG_BETA ALANINE_METABOLISM
KEGG_LIMONENE AND_PINENE_DEGRADATION
KEGG_RIBOSOME
KEGG_PROTEASOME
KEGG HOMOLOGOUS_RECOMBINATION
KEGG_CHEMOKINE SIGNALING_PATHWAY
KEGG_OOCYTE_MEIOSIS
KEGG_REGULATION OF_AUTOPHAGY
KEGG_APOPTOSIS
KEGG_AXON_GUIDANCE
KEGG_LYSOSOME
KEGG_SYSTEMIC LUPUS_ERYTHEMATOSUS
KEGG_ASCORBATE AND_ALDARATE_METABOLISM
KEGG_GLYCINE_SERINE AND_THREONINE_METABOLISM
KEGG_VALINE_LEUCINE AND_ISOLEUCINE_DEGRADATION
KEGG_BETA ALANINE_METABOLISM
KEGG_INOSITOL PHOSPHATE_METABOLISM
KEGG_GLYOXYLATE AND_DICARBOXYLATE_METABOLISM
KEGG_BIOSYNTHESIS_OF UNSATURATED_FATTY_ACIDS
KEGG_RIBOSOME
KEGG_NUCLEOTIDE EXCISION_REPAIR
KEGG_CHEMOKINE SIGNALING_PATHWAY
KEGG_ENDOCYTOSIS
KEGG_WNT SIGNALING_PATHWAY
Survival probability
subtype1
subtype2
p = 0.00222
Time: months (overall survival)
p = 0.02047
Time: months (progression free survival)

E

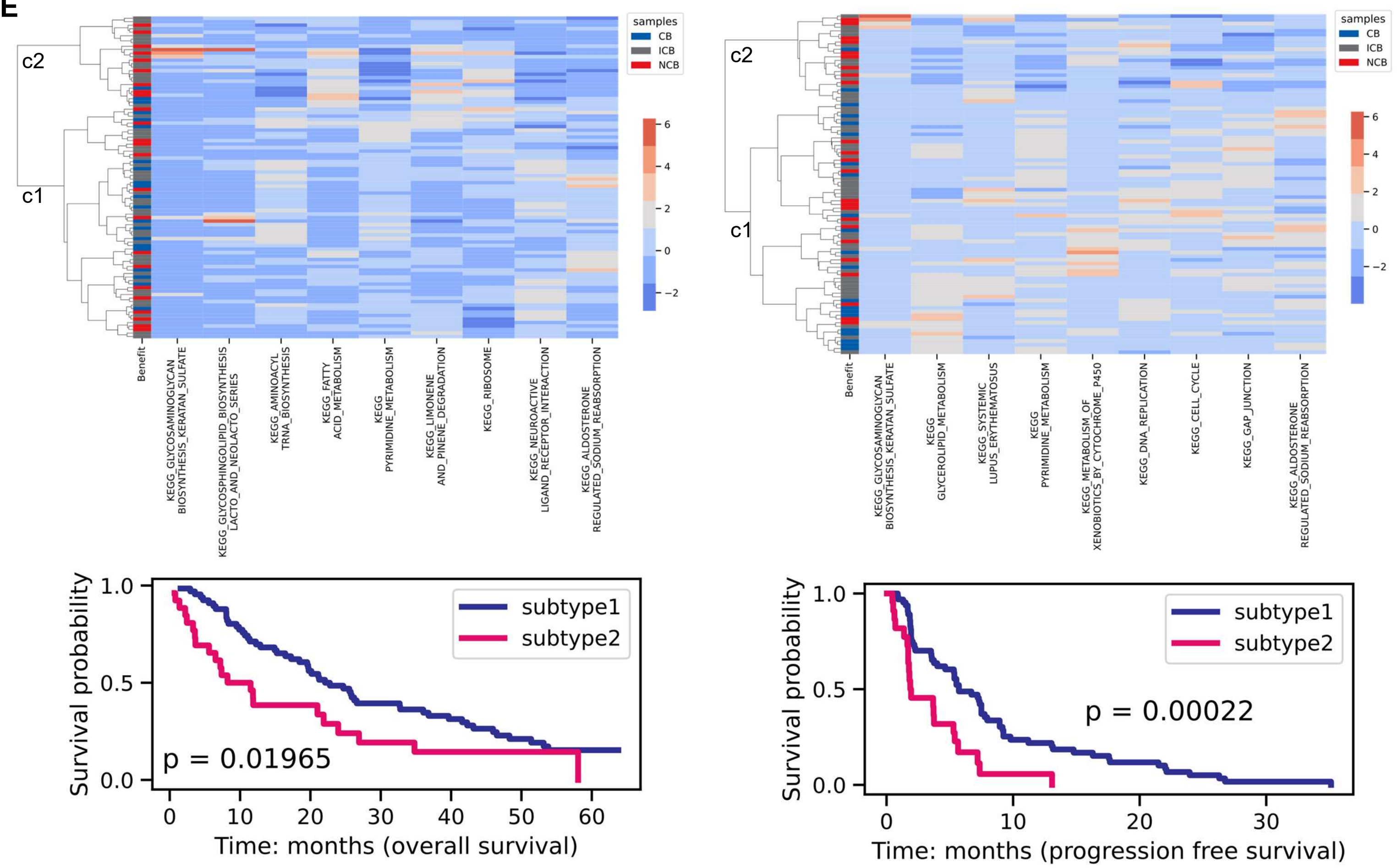


**Supplementary Figure S19. Clustermap and survival analysis of pE samples using pathway scores.** (A). 1 (OS)/ 5 (PFS) significant pathways based on GSVA. (B). 8 (OS)/ 8 (PFS) significant pathways based on entropy. (C). 12 (OS)/ 10 (PFS) significant pathways based on gene eigenvector centrality scores. (D). 16 (OS)/ 14 (PFS) significant pathways based on gene closeness centrality scores. (E). 9 (OS)/ 9 (PFS) significant pathways based on edge betweenness centrality score.

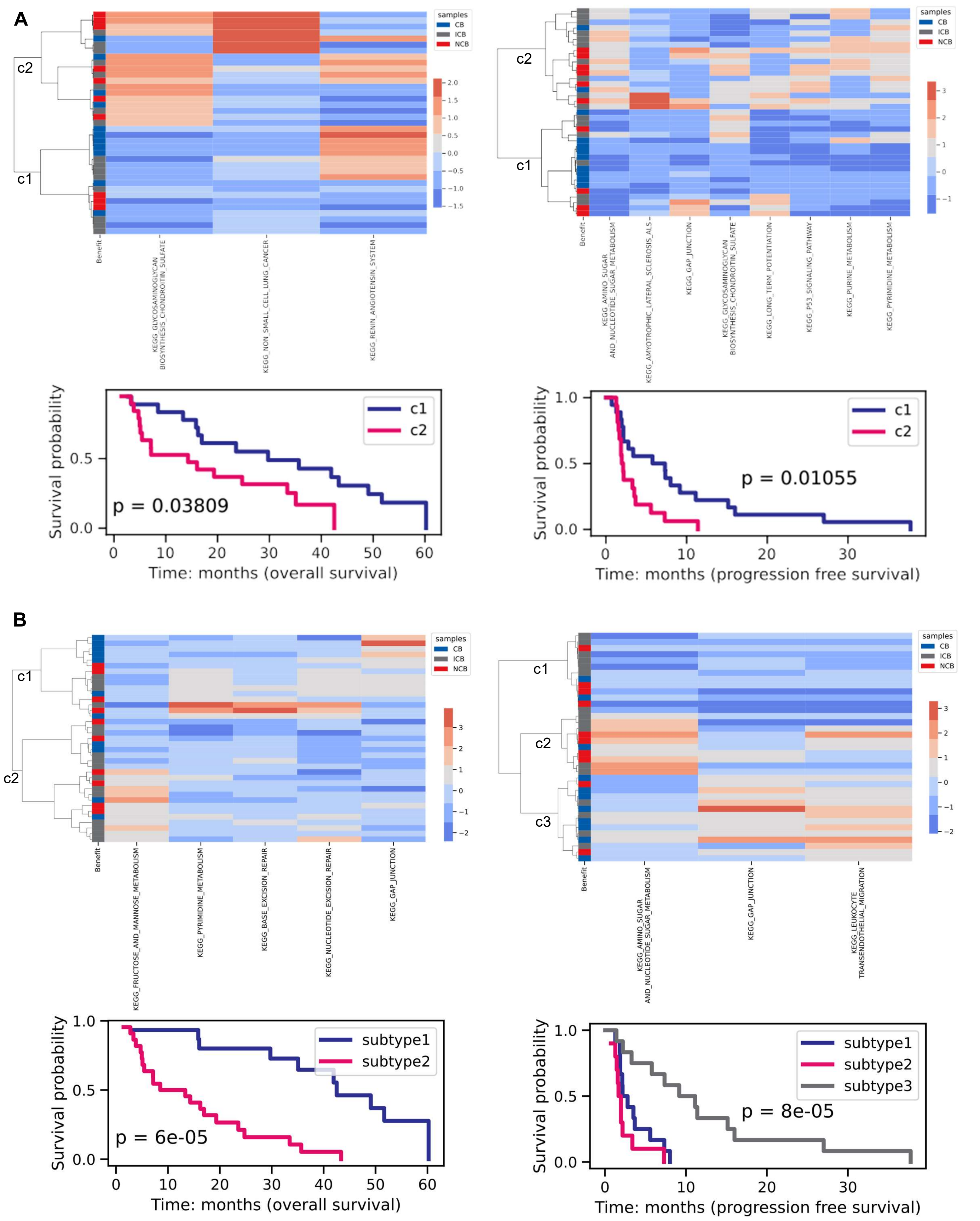

A
c2
c1
samples
CB
ICB
NCB
Benefit
KEGG_GLYCOSAMINOGLYCAN_BIOSYNTHESIS_CHONDROITIN_SULFATE
KEGG_NON_SMALL_CELL_LUNG_CANCER
KEGG_RENIN_ANGIOTENSIN_SYSTEM
KEGG_AMINO_SUGAR_AND_NUCLEOTIDE_SUGAR_METABOLISM
KEGG_AMYOTROPHIC_LATERAL_SCLEROSIS_ALS
KEGG_GAP_JUNCTION
KEGG_GLYCOSAMINOGLYCAN_BIOSYNTHESIS_CHONDROITIN_SULFATE
KEGG_LONG_TERM_POTENTIATION
KEGG_P53_SIGNALING_PATHWAY
KEGG_PURINE_METABOLISM
KEGG_PYRIMIDINE_METABOLISM
Survival probability
p = 0.03809
Time: months (overall survival)
p = 0.01055
Time: months (progression free survival)
B
c1
c2
c3
KEGG_FRUCTOSE_AND_MANNOSE_METABOLISM
KEGG_PYRIMIDINE_METABOLISM
KEGG_BASE_EXCISION_REPAIR
KEGG_NUCLEOTIDE_EXCISION_REPAIR
KEGG_GAP_JUNCTION
KEGG_AMINO_SUGAR_AND_NUCLEOTIDE_SUGAR_METABOLISM
KEGG_GAP_JUNCTION
KEGG_LEUKOCYTE_TRANSENDOTHELIAL_MIGRATION
subtype1
subtype2
subtype3
p = 6e-05
Time: months (overall survival)
p = 8e-05
Time: months (progression free survival)

C

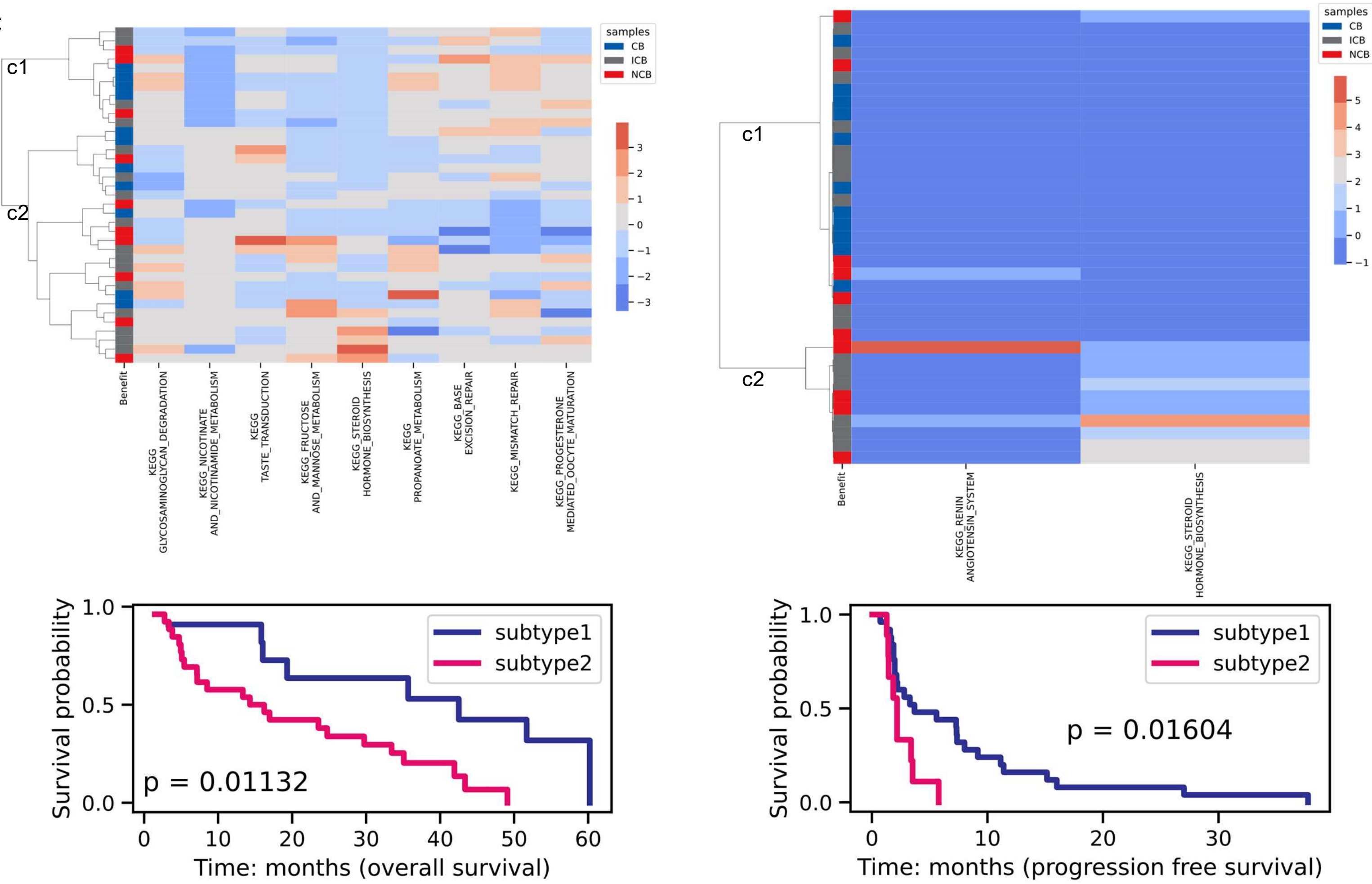

c1
c2
samples
CB
ICB
NCB
Benefit
KEGG_GLYCOSAMINOGLYCAN_DEGRADATION
KEGG_NICOTINATE_AND_NICOTINAMIDE_METABOLISM
KEGG_TASTE_TRANSDUCTION
KEGG_FRUCTOSE_AND_MANNOSE_METABOLISM
KEGG_STEROID_HORMONE_BIOSYNTHESIS
KEGG_PROPANOATE_METABOLISM
KEGG_BASE_EXCISION_REPAIR
KEGG_MISMATCH_REPAIR
KEGG_PROGESTERONE_MEDIATED_OOCYTE_MATURATION
KEGG_RENIN_ANGIOTENSIN_SYSTEM
KEGG_STEROID_HORMONE_BIOSYNTHESIS
Survival probability
subtype1
subtype2
p = 0.01132
Time: months (overall survival)
p = 0.01604
Time: months (progression free survival)


D

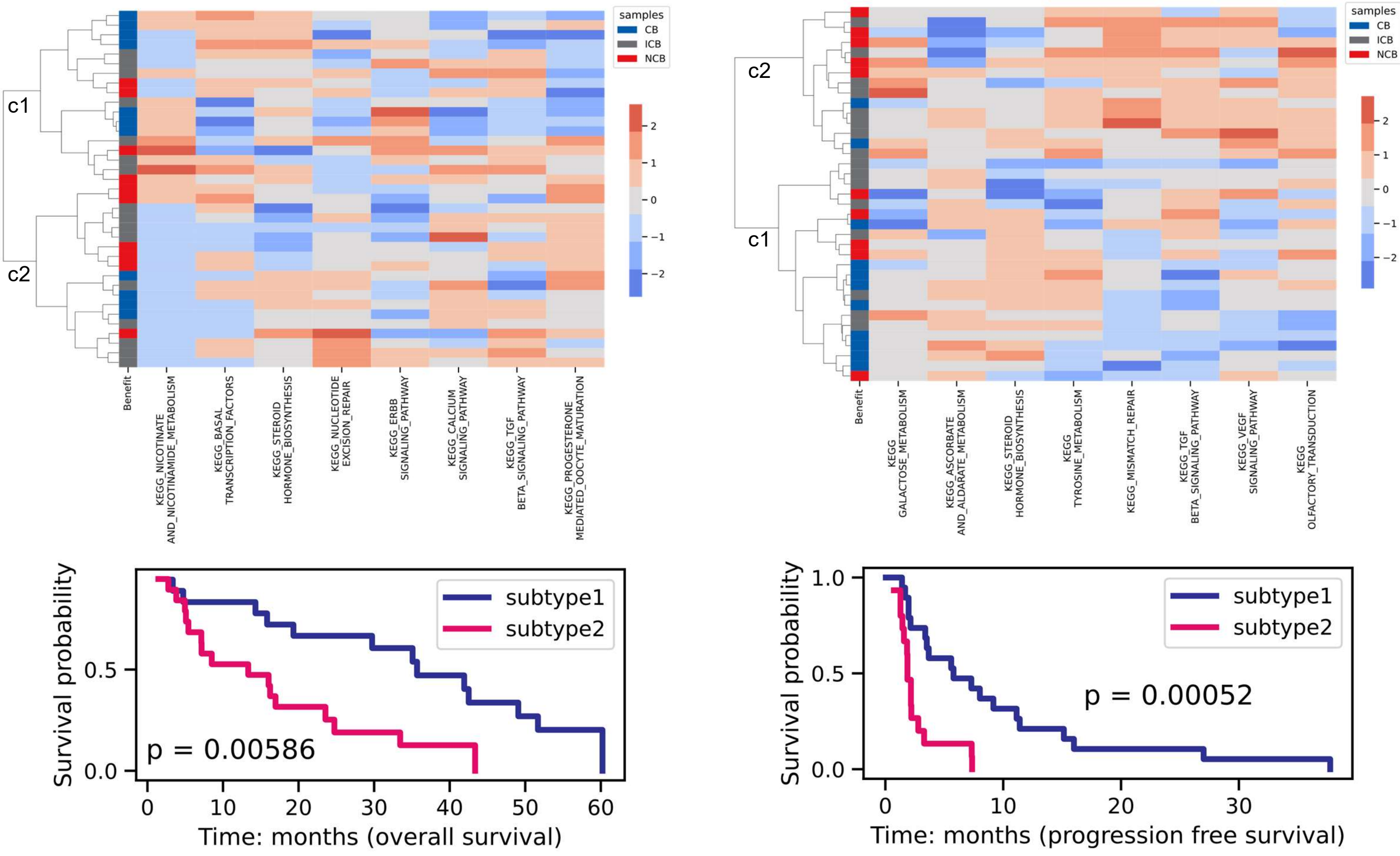

c1
c2
samples
CB
ICB
NCB
Benefit
KEGG_NICOTINATE_AND_NICOTINAMIDE_METABOLISM
KEGG_BASAL_TRANSCRIPTION_FACTORS
KEGG_STEROID_HORMONE_BIOSYNTHESIS
KEGG_NUCLEOTIDE_EXCISION_REPAIR
KEGG_ERBB_SIGNALING_PATHWAY
KEGG_CALCIUM_SIGNALING_PATHWAY
KEGG_TGF_BETA_SIGNALING_PATHWAY
KEGG_PROGESTERONE_MEDIATED_OOCYTE_MATURATION
KEGG_GALACTOSE_METABOLISM
KEGG_ASCORBATE_AND_ALDARATE_METABOLISM
KEGG_STEROID_HORMONE_BIOSYNTHESIS
KEGG_TYROSINE_METABOLISM
KEGG_MISMATCH_REPAIR
KEGG_TGF_BETA_SIGNALING_PATHWAY
KEGG_VEGF_SIGNALING_PATHWAY
KEGG_OLFACTORY_TRANSDUCTION
Survival probability
subtype1
subtype2
p = 0.00586
Time: months (overall survival)
p = 0.00052
Time: months (progression free survival)

**E**

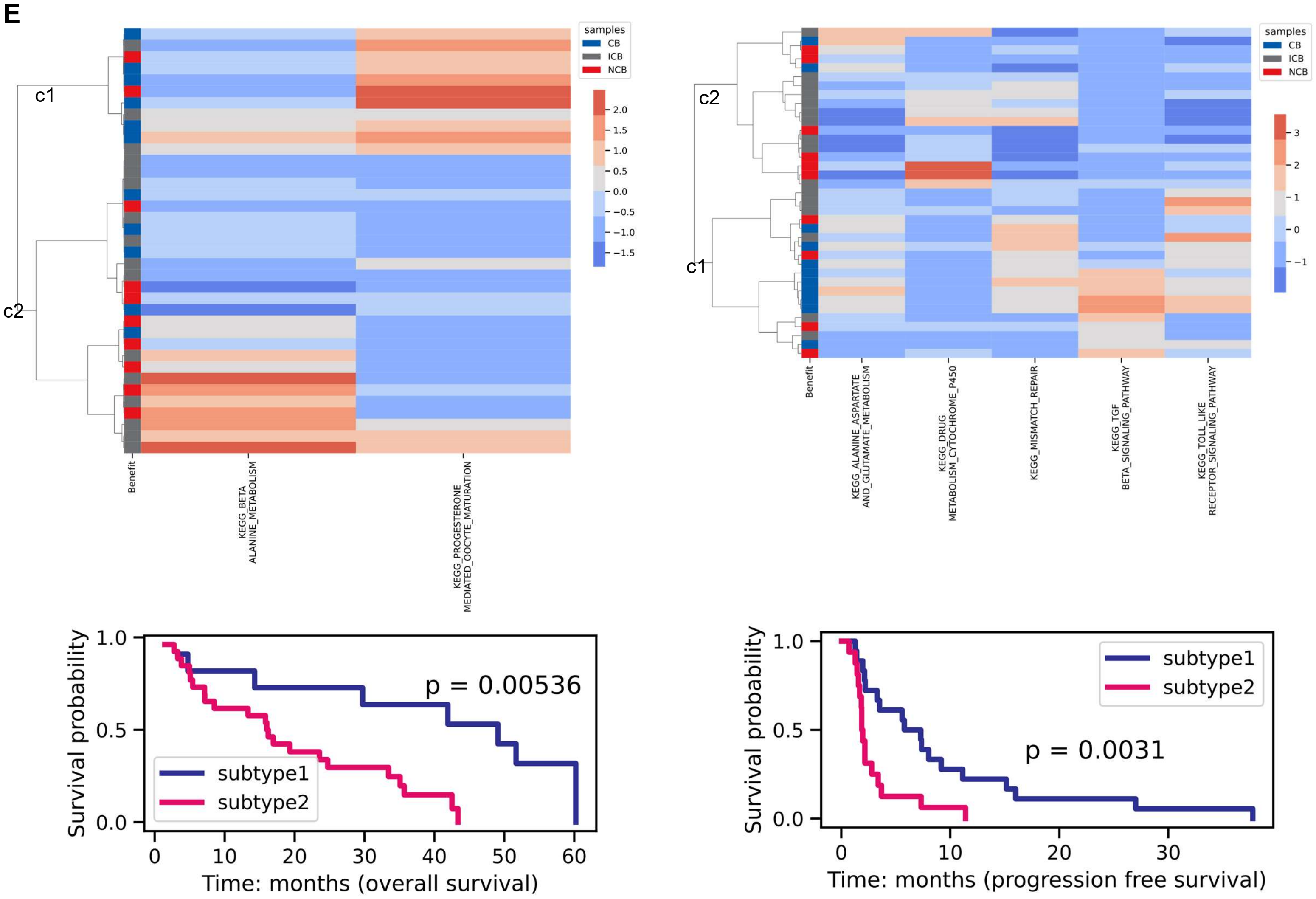


**Supplementary Figure S20. Clustermap and survival analysis of mE samples using pathway scores.** (A). 3 (OS)/ 8 (PFS) significant pathways based on GSVA. (B). 5 (OS)/ 3 (PFS) significant pathways based on entropy. (C). 9 (OS)/ 2 (PFS) significant pathways based on gene eigenvector centrality scores. (D). 8 (OS)/ 8 (PFS) significant pathways based on gene closeness centrality scores. (E). 2 (OS)/ 5 (PFS) significant pathways based on edge betweenness centrality score.

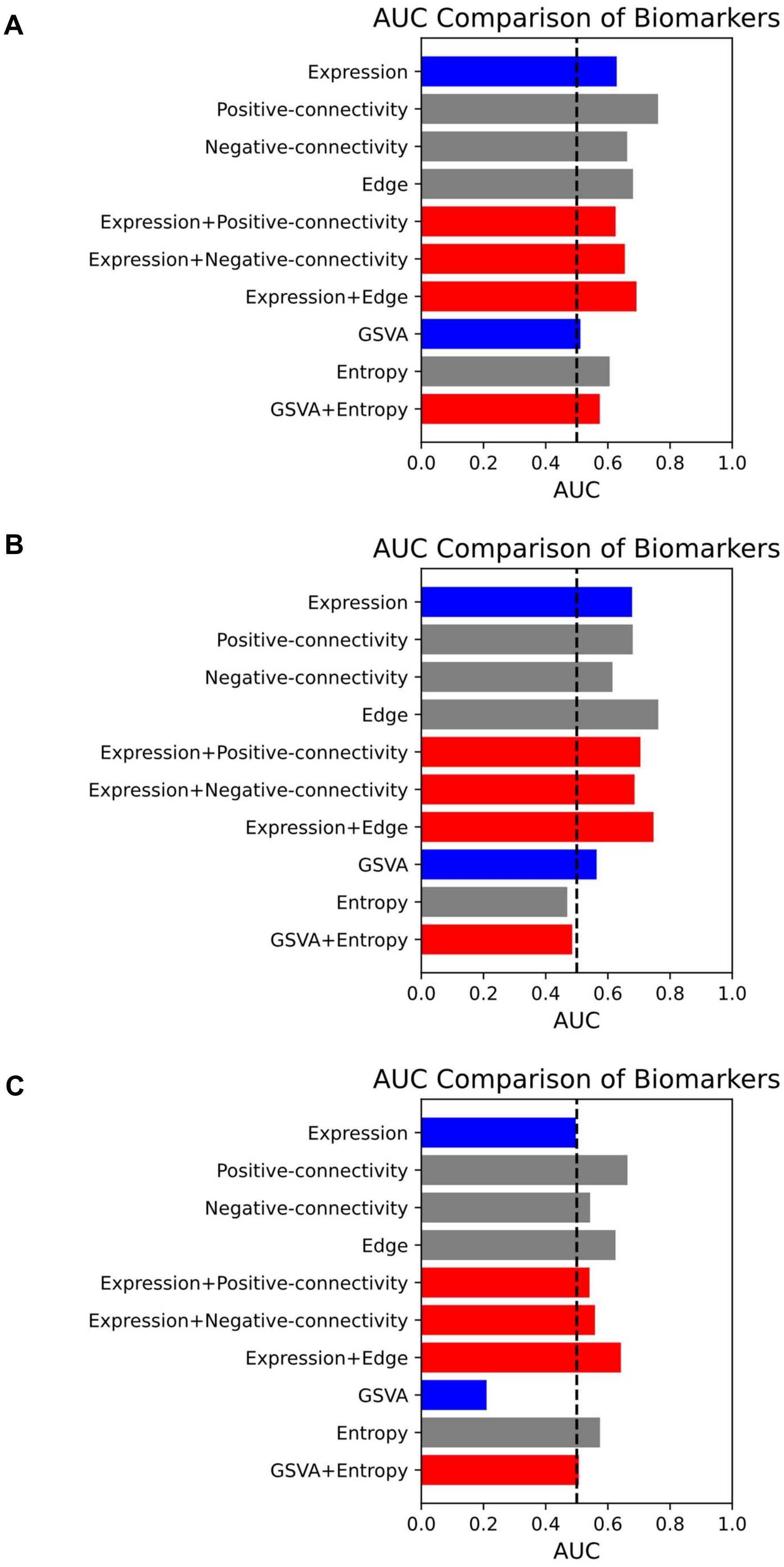


**Supplementary Figure S21. Leave-one-out cross validation (LOOCV) method evaluated the performance of ML models based on gene expression input, network feature input and their combination.** (A). Logistic regression (LR) model. (B). Random forest (RF) model (C). Support vector machine (SVC) model.

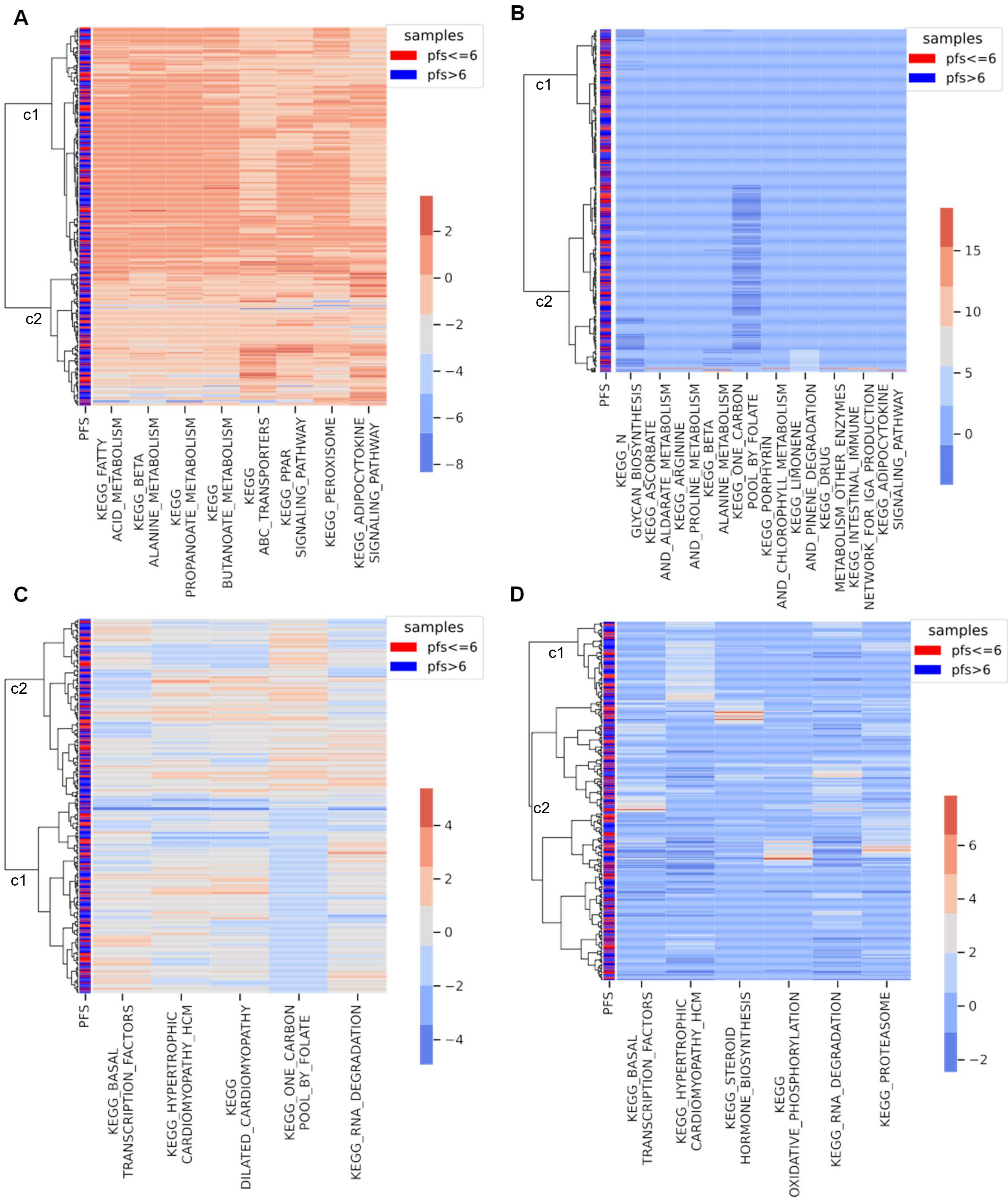


**Supplementary Figure S22. Clustering samples treated with Avelumab plus axitinib.** A. Edge entropy. B. Eigenvector centrality scores. C. Closeness centrality scores. D. Edge betweenness centrality scores.
